# CloneCoordinate: Open-source software for collaborative DNA construction

**DOI:** 10.1101/2025.07.04.663221

**Authors:** Ethan Jeon, Ziyang Shen, Santiago Christ, Evelyn Qi, Ida Fan, Nawon Lee, Odysseas Morgan, Madeline Ohl, Dylan Millson, Michelle Laker, Aspen Pierson, Emily Villegas Garcia, Yifan Zhang, Adeline Choi, Ashrita Iyengar, Rebecca Kim, Josh Lee, Linden Niedeck, Vanessa Oeien, Maliha Rashid, Nandini Seetharaman, Arnav Singh, Delaney Soble, Jenny Yu, Katherine Yu, Simms Berdy, Ellia Chang, Robin Kitazono, Sofija Ortiz, Dylan Taylor, B Thuronyi

**Affiliations:** Williams College, Williamstown, MA 01267, USA

## Abstract

Custom DNA constructs have never been more common or important in the life sciences. Many researchers therefore devote substantial time and effort to molecular cloning, aided by abundant computer-aided design tools. However, support for managing and documenting the construction process, and for effectively handling and reducing the frequency of setbacks, is lacking. To address this need, we developed CloneCoordinate, a free, open-source electronic laboratory notebook specifically designed for cloning and fully implemented in Google Sheets. By maintaining a real-time, automatically prioritized task list, a uniform physical sample inventory, and standardized data structures, CloneCoordinate enables productive, collaborative cloning for individuals or teams. We demonstrate how the information captured by CloneCoordinate can be leveraged to troubleshoot assembly problems and provide data-driven insights into cloning efficiency. CloneCoordinate offers a new and uniquely accessible model for how to carry out, and iteratively improve on, real-world DNA assembly.

## Introduction

Our ability to study, manipulate, and engineer living systems has never been greater, thanks in large part to advances in DNA synthesis and assembly technologies. Techniques that once required specialized training and equipment are now routine and supported by straightforward commercial reagents and kits. The cost and speed of both custom oligonucleotide synthesis and rapid-turnaround DNA sequencing have dropped by orders of magnitude.^1^ High-efficiency DNA assembly methods now abound and dramatically increase the scope of construct design that can be attempted with acceptable success rates.^2^ Consequently, work across molecular biology, biochemistry, chemical biology, synthetic biology, and the life sciences in general has increasingly leveraged novel genetic constructs to address research goals.

The prominent role of novel genetic material in research means that, for many groups, cloning has long been mission-critical work that constitutes a substantial fraction of overall time and effort. Though many commercial vendors or academic/nonprofit biofoundries offer complete, custom plasmids on a fee-for-service basis, the costs and/or turnaround times are such that in-house cloning is often preferred by academic labs, especially when rapid iteration on genetic designs is needed to advance research. The fact that cloning efficiency and scope can limit the rate of overall scientific progress for many labs shows that it is not a trivial problem and highlights the need for its continued improvement.

Major factors that make cloning challenging are (1) the logistical and organizational complexity, which scales more than linearly with the number and variety of constructs being built; and (2) the non-negligible and poorly predictable rate of failures or delays throughout the process, even when using common high-efficiency methods. This second factor also hampers the decision to outsource since time and cost cannot be known with certainty in advance for either outsourced or in-house cloning.

Lack of standardization and support for DNA construction also creates barriers to both entry and advancement in the discipline. The large number of disparate steps involved may be individually simple, but cumulatively require substantial training and time to complete even a single construct. The diversity of available approaches to cloning generates confusion for novices. Troubleshooting typically relies on heuristics arising from long experience, emphasized by an oral tradition and apprenticeship-based training. Acquiring the expertise needed for high productivity is therefore a difficult and time-consuming effort that each individual must make with little structural support.

While many effective software tools now support construct design and/or the planning of the build process (for example, generating primer, PCR, and assembly plans that include manual or robotics instructions),^3–7^ analogous tools for tracking the physical DNA construction process are extremely rare. With few exceptions, build *planning* tools do not cover the data collection, task management, or troubleshooting involved in build *execution*. The few that appear to do so are poorly accessible to academic labs or startups. Commercial platforms targeted toward industry users are closed source and costly,^8^ while non-profit alternatives are technically demanding to deploy and maintain and therefore probably best suited to integrated biofoundries.^9,10^

We note that although new DNA assembly methods (whether molecular, kit- or equipment-based, robotically automated, or a combination) are regularly published and their value demonstrated using small cloning test sets, the basic workflow of cloning has not substantially changed in decades (**Figure 1a, left**).^11,12^ Expanded technical capabilities have raised expectations and ambitions along with efficiency so that the overall success rate of cloning still constrains progress. We believe this problem is exacerbated by a disconnect between targets for innovation in cloning methodology and the challenges that commonly arise in real-world cloning experiences.

**Figure 1.**
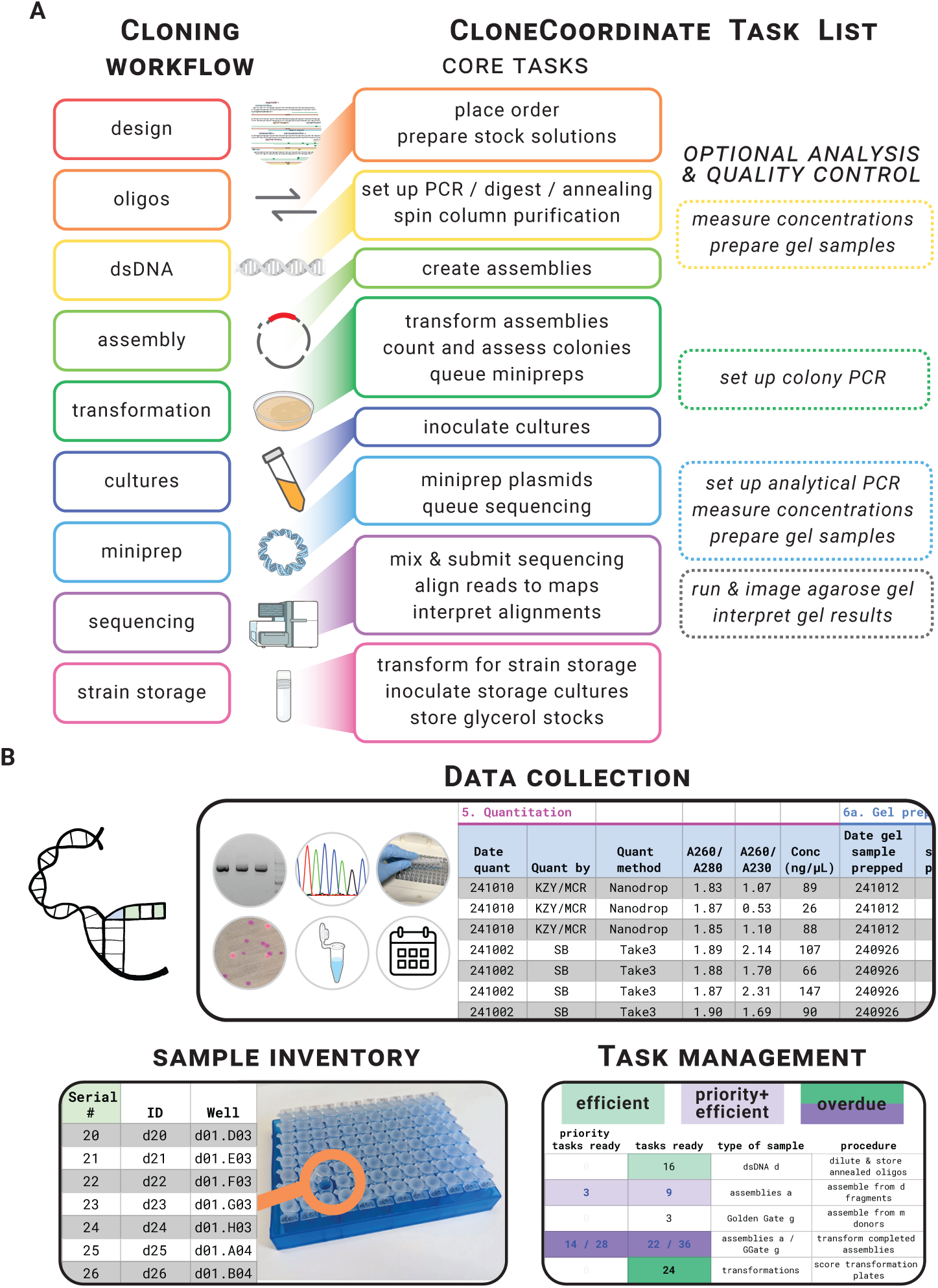
CloneCoordinate manages the cloning workflow and associated data, samples, and tasks. A, Cloning and CC workflows. The general steps of the cloning process are broken down by CC into a list of discrete and clearly delineated tasks, shown schematically here and displayed in detail on CC’s Dashboard tab (**Supplementary Figure 2**). **B, General CC features.** Data collected by CC includes, but is not limited to, tabulation of agarose gel electrophoresis, colony counts and colors, sequencing results, volumes and components used for all operations, instrument settings/conditions, and date stamps for all steps. The sample inventory assigns unique identifiers to physical samples that can be stored long-term in well plates for easy access. The task priority system directs work effectively by highlighting tasks on the Dashboard that meet user-set criteria for efficiency (based on number of samples to be processed), priority (based on manual flagging of specific constructs as high-priority), and elapsed time since the task was last done (with steps that have been done least recently marked for attention).

To meaningfully improve the cloning process, we consider it essential to regularly gather detailed and granular data about how it is actually done – across diverse constructs, researchers, contexts, and projects, and including failures and abandoned or repeated operations. Such data are indispensable for meaningful evaluation of alternative strategies or methods, for applying the design-build-test-learn cycle to cloning itself, and for applying machine-learning tactics which have shown enormous success in other fields.^13,14^

These data are largely unavailable, either because they are not captured at all, or because, in the absence of any accessible, standardized system, idiosyncratic approaches are used that mean the data are not FAIR (Findable, Accessible, Interoperable, and Reusable)^15^ and cannot be easily leveraged for improvement. Despite advances in standardizing construct representation,^16^ composition and derivation,^17^ publication of new constructs almost never includes detailed cloning data (and sometimes even construct sequences are omitted).^18^ Instead, methods are omitted or given only schematically without concrete details (e.g., concentrations and amounts of components, colony counts, phenotypic or genotypic success rates, etc.) at the level of individual constructs and samples that are essential for systematic analysis. This is understandable since the effort involved in comprehensively documenting cloning without software support is enormous and usually unrewarding.

A platform that seeks to support and capture data about cloning execution faces significant challenges. Any new approach to a familiar process must overcome the inertia of established practice and expertise. It must offer usability and efficiency benefits to make the overhead cost of adoption worthwhile, ideally for practitioners of any experience level. Both initial setup and ongoing operation must be as simple as possible, ideally at no cost and with minimal equipment requirements. Software with a graphical interface that does not require source code compilation or command line interactions is essential for broad accessibility. Operation and data storage should be transparent, accessible, and extensible for end users. Finally, high-quality, pre-built data analysis tools should quickly reward the investment of adopting and entering data into the platform by providing actionable information.

We have worked to implement these principles in development of cloning management software which we call CloneCoordinate. Our aim is to increase both the accessibility and effectiveness of cloning for the widest possible range of practitioners, while making its detailed documentation both manageable and rewarding, enabling collection of standardized cloning data sets at a meaningful scale.

## Results

### CloneCoordinate provides end-to-end task and data management coverage for cloning

CloneCoordinate (CC) is a free, open-source electronic lab notebook specialized for cloning that covers the entire DNA construction process (**Figure 1**). CC provides scaffolded data entry and logging for all cloning-related operations, a comprehensive physical sample inventory, and task and status tracking that updates live to reflect the progress (or setbacks) of lab work.

CC is implemented entirely within Google Sheets. The broadly familiar spreadsheet interface makes data entry straightforward. Sheets operates on any platform and practically any device, supports simultaneous multi-user editing, and automatically maintains version history. Installation is as simple as creating a copy (an “instance”) of the main CC Sheet (available at clonecoordinate.org) and sharing it with other users (**Supplementary Text 1**). Separate Sheets for data analysis/visualization and tools, also provided, can be connected to a CC instance by entering its URL. Users retain full control over their CC instance and its data through Google access permissions.

Each step in cloning and its associated samples and operations (e.g., oligos, transformations, minipreps) is tracked on a tab of CC (**Figure 1a** and **Supplementary Figure 1**). Operations are logged by filling in a set of designated columns, working from left to right, including dates, measurement results, volumes, and conditions used, etc. Required, optional, and informational (formula-filled) fields are marked with distinctive header colors (green, blue, and white, respectively). Data entries are validated automatically using consistency check formulas, format requirements, and/or drop-down menus, and formulas are protected from accidental modification.

CC implements a standardized inventory that assigns unique identifiers to all physical samples involved in the workflow (**Figure 1b**). These correspond to specific storage locations (by default, in 96- well plates) within a single collection shared by all users of a CC instance. This approach streamlines sample organization, storage, and retrieval, ensuring that materials needed for bench work are always ready to hand. It guarantees unambiguous specification of how each step in cloning is carried out and allows CC to track sample properties such as remaining volume, concentration, and analysis outcomes. CC can support most sample storage methods. We use 8-tube strips with attached caps arranged into 96-position racks and have stably maintained >7,000 physical samples for multiple years in only a few cubic feet of space.

CC provides a real-time, prioritized task list, shown on its Dashboard tab (**Supplementary Figure 2**), by drawing on data and inventory standardization (**Figure 1b**). This breaks the cloning workflow into a series of independent, narrowly scoped operations. As these steps are carried out and logged, the status of each sample is automatically updated and compiled into a to-do list. Tasks that are ready, representing steps in the cloning process across multiple constructs at different stages of completion, can be completed independently and in any order. CC is method-agnostic, so that any existing protocol, including physical automation (robotics) or even outsourcing, can be used to accomplish each operation, and the results entered into CC’s generalized data fields.

Participants are free to work linearly on advancing specific constructs, but CC also steers their effort toward tasks that provide good return on time investment. This is done using a customizable multilayer priority system (**Figure 1b)**. Tasks are highlighted when the number of samples ready for a given operation exceeds a user-customizable value. This promotes efficiency and economies of scale, e.g., by enabling effective multichannel pipette or liquid handler use, and/or distributing fixed overhead costs like setup or incubation time across more samples. Additionally, particular target constructs can be manually designated high-priority, marking all their associated operations with separate tallies and highlighting. Steps that have not been done for a relatively long time are also marked for attention to keep the overall pipeline balanced.

The sample statuses that generate the task list take step dependencies, customizable settings, and molecular biology logic into account (**Supplementary Table 1**). For example, a Gibson assembly becomes ready to do when all its component DNA fragments have been generated, while it becomes stalled if the PCR for a fragment is found to have failed (**Supplementary Figure 3**). CC can be configured to require optional analytical steps for sample quality control (**Figure 1a**) on a global and/or per-sample basis according to group and researcher preferences (**Supplementary Text 2**). CC version 1.0 supports many commonly used cloning flowcharts with potential for future expansion based on user feedback.

### CloneCoordinate enables new cloning paradigms for efficiency, collaboration, and labor allocation

CC allows researchers to reframe traditional cloning by productively managing and distributing interactions with the process in ways that are otherwise impractical or impossible.

Abstraction is a key principle for efficient research because it allows focus on high-level goals without distractions from the details of implementation. CC isolates the build execution phase of DNA construction and frees researcher cognitive load for other objectives. CC captures complete plans for cloning each target construct. It is design tool agnostic so that any method, from manual to fully automated design tools, can be used to generate build instructions specifying the unique assembly steps for each construct (**Supplementary Figure 1**). These instructions are then entered into CC and the construct becomes “queued” for construction.

CC does provide one tool to support construct design using the Golden Gate cloning method.^19–21^ CC’s Registry stores information about plasmids and other samples created (or obtained) as Golden Gate cloning part donors, including restriction enzyme and overhang sequences generated. This information is read by the “Golden Gate assembly queuer” Accessory Sheet (first reported as part of the Vnat Golden Gate Collection), which allows users to select a set of compatible parts from dropdown menus that will yield a complete plasmid according to Golden Gate assembly rules (**Supplementary Figure 4**). This tool greatly simplifies the Golden Gate design process and avoids part selection errors. Designs can be pasted into CC’s Golden Gate tab and are automatically matched to CC’s inventory of physical samples.

Once a construct is queued, all cloning steps (including any necessary troubleshooting; see below) are handled within CC until that construct is complete. CC provides information about part locations, availability, and properties (such as concentration) and automates calculations for assembly mixtures and other bench steps. The self-contained CC cloning process is therefore isolated by abstraction layers from the design and test phases. This makes it possible to queue and build many more constructs simultaneously than any individual designer could keep track of by hand and also allows pooling of cloning steps across many participants and/or projects without increasing complexity.

To connect the design and test phases regardless of how complex or prolonged the intervening build phase becomes, CC provides an Experiment tracker. While CC’s construct Registry captures construct metadata including design goals, the Experiments tab focuses on planned testing. Each Experiment links a set of constructs (e.g., a series of design variations together with positive and negative controls) to planned testing conditions, typically entered at the same time as the designs (**Supplementary Figure 5**). When CC detects that all required constructs are cloned, sequence-verified, and ready to use, the relevant researcher can be automatically alerted by email that the Experiment is ready to execute. In the meantime, the Experiment monitors the build status of each associated construct and will generate an email alert if any become stalled during cloning and require manual intervention. This approach allows isolation of the build phase from the design and test phases in time, attention, and even responsibility.

While most benefits of CC apply to any number of users, CC enables a collective approach to cloning (i.e., across a project team or entire laboratory) with unique benefits. Its single-inventory framework guarantees sample access and unambiguous instructions, making tasks accessible to any participant while also providing a complete audit trail for the build process. It removes barriers to reusing parts (e.g., sequencing primers, plasmid backbone digests) efficiently across constructs, projects, or individuals. It prevents information loss from siloing, revealing duplicated work and failure modes that cut across individuals, for example problems with shared equipment or reagents.

Collective cloning unlocks several economies of scale that reduce costs and increase throughput. With enough participants, cloning steps can be carried out continuously and seamlessly throughout the entire work day and work week without requiring any individual to contribute more than a few operations, freeing up their time for other purposes (**Figure 2**). When many target constructs are active in a cloning pipeline, each step can be carried out on a large batch (many samples per operation), bringing numerous advantages. Frictionless order pooling allows oligonucleotides to be purchased in mass-normalized 96- well-plate format, or sequencing reactions ordered at bulk rates. Shared equipment use can be optimized, e.g., combining PCRs in one thermocycler block, and labor-saving tools such as multichannel or electronic pipettes, robotics, and multiwell plates can be regularly brought to bear. For example, our lab routinely carries out 48 minipreps simultaneously from 4 mL cultures grown in two 24-well deep well plates, saving sample handling time compared to individual culture tubes and facilitating multichannel and electronic pipette use. This can be completed within 3 hours even by relatively inexperienced trainees, under 4 minutes per miniprep. Likewise, large-batch enzyme reactions reduce error rates, labor per sample, and reagent waste through efficient use of premixes (common reagents combined once and then divided rather than pipetted individually). CC’s “Assistant” tabs support this bench work, supplying customizable premix calculations, checklists for component addition, and color-coded sample locations to reduce mistakes (**Supplementary Figure 6** and **Supplementary Text 4**).

**Figure 2.**
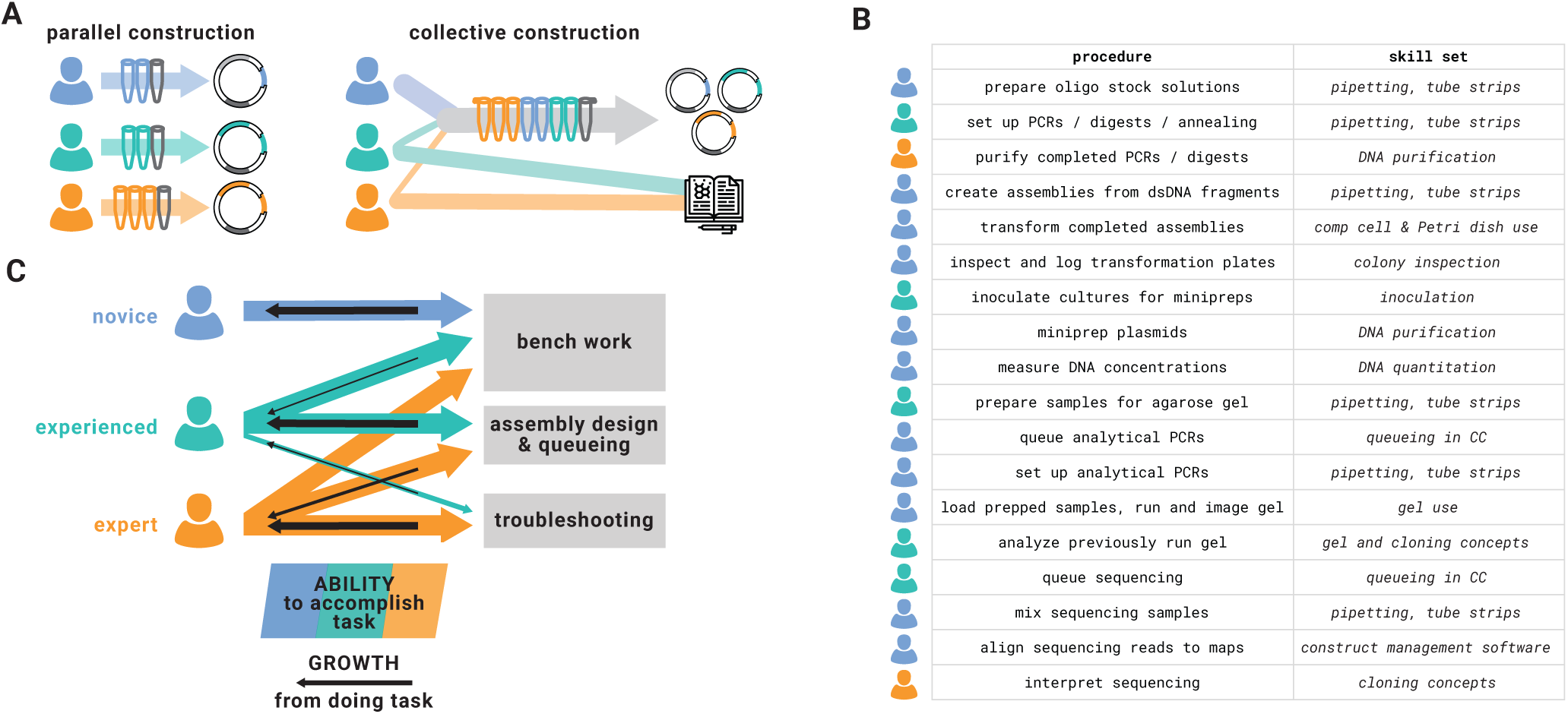
CloneCoordinate’s task management enables collective DNA construction and improves efficiency, accessibility, and participant development. A, Schematic illustration of parallel vs. collective DNA construction. Several individuals working independently to generate distinct target constructs must each carry out cloning operations on their own samples, including the fixed overhead effort needed to assemble materials and tools and set up equipment. If the same individuals can efficiently pool information about their designs and the operations needed to create them, one person at a time can handle each cloning operation. This requires only marginally increased effort to handle the additional samples, particularly by leveraging scale- efficient methods such as multichannel pipette use. In some cases, components common to multiple, related designs (gray tubes) can be reused, reducing redundancy. Effort and attention are freed for other research activities. CC makes this possible by providing unambiguous build instructions and sample IDs and coordinating tasks. **B, Abridged table of CC cloning procedures and skill sets involved.** Participant icons give a representative illustration of how labor might happen to be allocated from 3 hypothetical participants across a workflow during 1 round of cloning so that each step is covered. The skill set column highlights common and/or distinct requirements for each step to guide inexperienced participants in selecting work to do. No single participant necessarily needs to be capable of carrying out all steps. **C, Schematic illustration of individuals’ capabilities vs. opportunity for development as a function of expertise level.** A novice participant is most capable of doing hands-on tasks with clearly defined protocols, and also to find these tasks novel and developmentally rewarding. Conversely, they may be unable to queue new designs or troubleshoot assembly problems without assistance. At greater experience levels, participants no longer learn much from less complex work even though they may be able to accomplish it effectively, and their time is better spent on the most challenging tasks. Collectivized DNA assembly allows participants to change how they deploy their labor as they become more experienced, and this benefits both individual development and the group’s effectiveness.

CC divides the cloning workflow into as many discrete, self-contained operations as possible, minimizing the time and expertise requirements for each one. Rather than requiring each participant to learn and successfully execute every step in the workflow in order to be productive, CC presents a nonlinear, abstracted, incremental approach. An individual participant can make a useful contribution after learning any single step, the minimum possible barrier to entry. Cloning can proceed effectively so long as the group as a whole is capable of handling the complete workflow, regardless of whether any single individual can do so. (**Figure 2b**). Normally, dividing up tasks and communicating accurately about each sample in the cloning process would present prohibitive logistical challenges, but CC’s structure allows seamless integration and organization of work across any number of participants.

By facilitating division of labor and expertise, collective cloning through CC provides substantial accessibility and inclusion benefits, especially for research groups that involve, or rely on, trainee and/or part-time researchers, such as undergraduate or high school students. These include Course-based Undergraduate Research Experiences (CUREs);^22^ International Genetically Engineered Machine (iGEM) competition teams;^23^ research groups at primarily or exclusively undergraduate institutions;^24^ community biotechnology labs; and hosts of Summer Undergraduate Research Fellowships. Building new DNA constructs in a timely and efficient manner can pose great difficulties; these settings and coordination of labor can both make this possible and extend the opportunity for meaningful participation to more, and more diverse, contributors. Our undergraduate-only research group at Williams College, whose members almost never enter with prior cloning experience, work mostly part-time, and rarely finish their involvement with more than 6 months’ full-time research experience, has seen this reflected in our own experience.

Within a diverse group of participants, CC allows allocation of labor to support both overall efficiency and personal development (**Figure 2c**). Individuals can choose tasks where they have comparative advantage from their expertise. The most experienced researchers might focus primarily on construct design, troubleshooting, and experiments, rather than the build process itself. Participants can select work that will contribute to their growth, practicing tasks in their zone of proximal development and shifting their focus over time as they become increasingly capable. Experience levels are made transparent by a CC Accessory Sheet that provides Statistics on all CC contributions, (**Supplementary Figure 7**) making it easy to find experts to consult about a given step (and, in our experience, encouraging friendly competition between lab members). It also allows mentors to track trainee progress over time (on the basis of performance output, not only time input) and advise them on next steps.

Finally, we find that a collective approach to DNA construction also benefits the entire research group and its scientific output. Project scopes can reach far beyond any one individual’s DNA-building capacity. They can also authentically leverage contributions from group members in early training stages rather than being driven only by the most experienced senior students. Every member of our group, whatever their duration of involvement or level of experience, can claim some ownership of the lab’s research achievements because these rely directly and transparently on their labor.

### CloneCoordinate is robust toward setbacks and informs troubleshooting

The comprehensive and standardized data collected by CC can be used to keep the cloning process organized and efficient and to facilitate troubleshooting even when cloning steps or strategies fail. The per-step failure rate is a major driver of the difficulty of cloning many constructs simultaneously. A large cloning batch can easily be initiated where all constructs are synchronized and progress together from one step to the next, but some steps invariably fail for a fraction of the samples. Constructs quickly become desynchronized, spread out across multiple construction stages. Manual backtracking creates logistical and organizational complexity and can erode efficiencies of scale. However, CC remains efficient even when cloning does not succeed as planned. Rather than construct-by-construct or batch- by-batch organization that is subject to attrition upon operation failure, CC uses task-by-task organization natively. This allocates labor to efficient steps whether batches remain synchronized or not.

CC participants must still make decisions about how best to overcome failures that block progression of each construct and queue the necessary operations to resume building. To facilitate this, CC’s “Construct Tracker” tool documents the status and history of each construct. It integrates information across construction steps and generates reports detailing the samples, analysis results, and outcomes involved in each assembly attempt (**Figure 3a**), including any previous setbacks and troubleshooting attempts.

**Figure 3.**
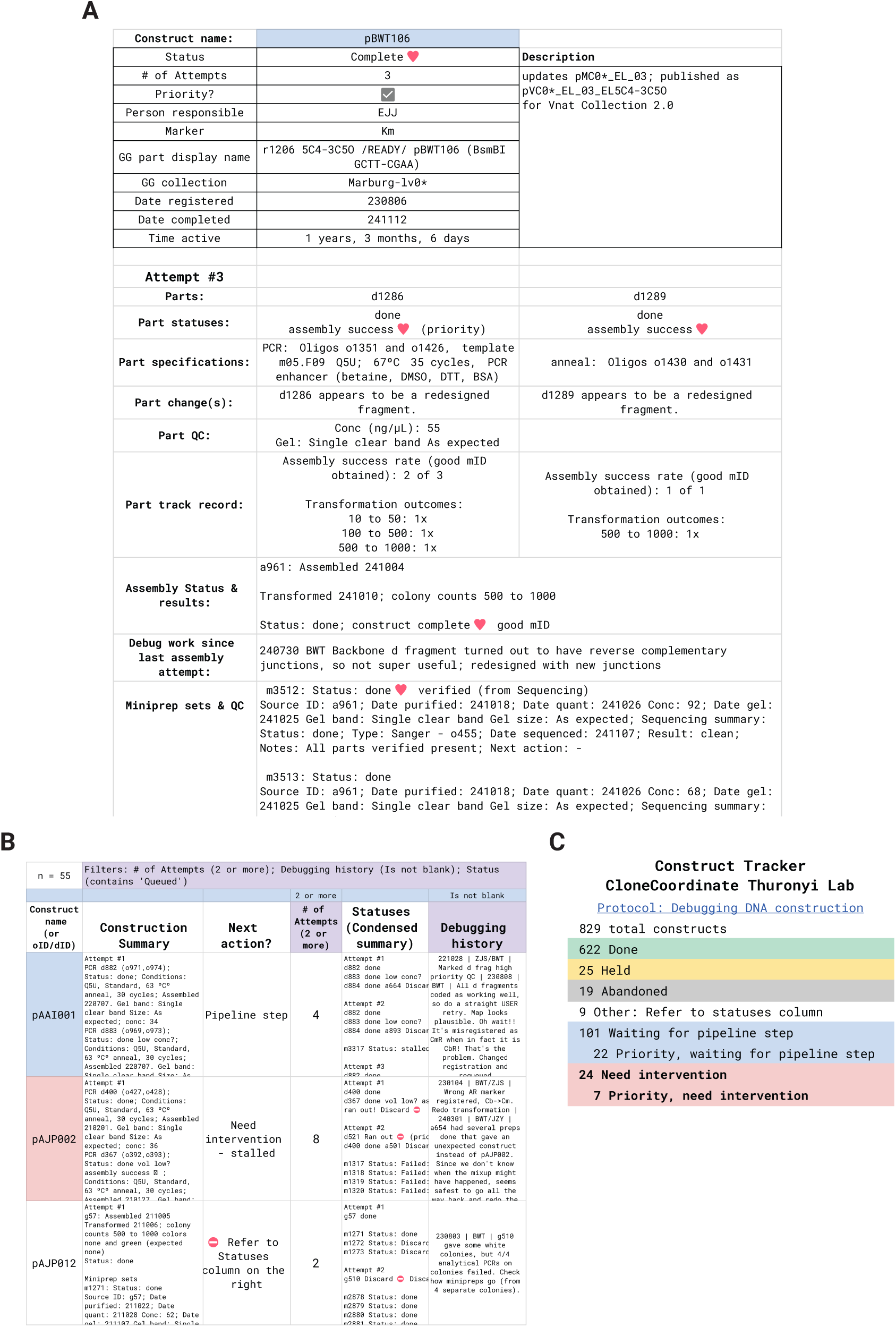
The CC Construct Tracker shows complete histories and troubleshooting information for each target construct. Screenshots are lightly modified for clarity. **A, Construct Inspector view**. This pane compiles information about all steps involved in building a plasmid across CC, organized by assembly attempt, to enable any participant to decide what next steps or new plans are needed. The construct shown was built successfully on a third attempt. The parts were redesigned after the discovery that the junction sequences were reverse complementary (a problem that CC now automatically detects and flags). One part (d1286) was reused in other constructs and successfully assembled in 2 other cases (all 3 of which gave colonies upon transformation). **B, Filtered Tracker view**. This pane lists each registered construct on its own row and provides a text summary of its history, a compilation of previous troubleshooting attempts, and the statuses of associated samples. Based on this information, the construct is classified as in progress (awaiting a CC pipeline step which is listed on the CC Dashboard) or stalled and needing manual intervention, such as queueing a new attempt. Other columns, not shown, tabulate construct properties, description, project information, and IDs of associated samples. The table can be filtered by various combinations of column values. **C, Construct Tracker Dashboard view**. This pane tabulates the overall construction statuses of all constructs and presents category totals. Tasks ready to do are tabulated according to the number of constructs needing them. Any participant can troubleshoot and restart constructs that “need intervention” so they can re-enter the CC pipeline.

Crucially, the Construct Tracker can incorporate and leverage information about outcomes from *other* constructs that may be relevant. For example, a PCR product for a plasmid backbone might be shared across several target constructs. If some of those constructs are built and successfully sequence-verified, this provides strong evidence that the PCR was sufficiently successful (independent of direct quality assessments such as concentration measurement or agarose gel electrophoresis) and is not the source of problems with a stalled construct that includes it. This information is incorporated into the PCR product’s Status field and tabulated in the Construct Tracker report. If it is unavailable, transformation colony counts can instead be used as a proxy for the part’s success. These tabulations represent our first steps toward leveraging CC data in more sophisticated ways to assist troubleshooting. Future goals include automated identification of common failure modes and data-informed recommendations for specific interventions.

The Construct Tracker presents a construct list that can be filtered by fields such as status, assembly method, project, or calculated properties such as number of previous assembly attempts, a proxy for troubleshooting difficulty (**Figure 3b**). To direct participant attention to needed troubleshooting, the Construct Tracker classifies constructs as in progress or needing intervention and tabulates the totals on a Dashboard (**Figure 3c**). This makes restarting work on stalled constructs a discrete participant task like those on the main CC Dashboard.

All participants have equal opportunity to troubleshoot any construct because all relevant information is available through the CC data compiled in the Construct Tracker’s history report. A participant can focus on constructs whose troubleshooting difficulty fits their experience level regardless of whether they personally designed them, or they can pursue the group’s collective cloning priorities, or their individual ones. After next steps are chosen and implemented in CC – e.g., retrying a PCR with different additives and queueing a new assembly attempt using the revised PCR – the troubleshooting rationale and plan are logged and dated in CC’s construct Registry and become part of the construct history.

### CloneCoordinate data and analytics allow broad-scale analysis of cloning effectiveness

Cloning is a complex multistep process that involves myriad decisions, from whether to purify PCR products before assembly, to how long to incubate DNA with competent cells. These decisions have diverse and disparate impacts on the labor, cost, and time efficiency of cloning. While the importance or unimportance of some factors is well known, the effects of most cloning practices on outcomes are obscure. Even minor factors can have significant cumulative effects, and factors can interact with each other or with properties of specific target constructs. Decision-making usually relies on individual researcher experience or widespread protocols, none of which are necessarily substantiated by data or controlled experimentation and may even be arbitrary. Lack of data makes it easy for bias or error to persist and stymies systematic improvement.

CloneCoordinate does not attempt to present any single “solution” to the multifactorial optimization problem of how to clone most effectively. However, by supporting collection of structured data, it makes optimization increasingly possible. Crucially, CC operates differently from studies that evaluate cloning using construct test sets.^6,25–28^ Rather than asking practitioners to assume that general findings about a method (however rigorously determined) will apply to their specific applications, CC data collection can evaluate whether the methods in use are effective *for the specific applications themselves*, an analysis that is relevant by definition.

To demonstrate that CloneCoordinate data can be leveraged to inform cloning process choices, we have assembled a data set from approximately 18 person-months of full-time cloning work, conducted across ∼6.3 total person-years of research by 34 undergraduate students (assuming 25% of total work time spent on cloning on average) over 6 calendar years at Williams College. We have completed 389 constructs, with 391 still in the queue as of this writing, through generation of >8,700 physical samples and logging of >44,000 discrete tasks. Although CC was under active development and changed significantly during this data collection period, we were able to capture or reconstruct the data needed for a strong foundation for analysis. Our data set, with sequence information and construct descriptions removed for privacy, is provided as a CC Sheet (**CloneCoordinate Thuronyi Lab 2019-2025 CC instance, Supplementary Data**).

To enable frictionless visualization of CC data, we developed several CC Analytics Sheets (CCAs). These can be connected to a CC instance and read data from it in close to real time. Each CCA Sheet implements a set of related, prebuilt analyses which are readily customizable without coding. For example, the “CCA Operations Timeline” tool can plot the cumulative total of up to 5 CC tasks (selected from dropdown menus) over time (**Supplementary Figure 8**), as well as the number of in-progress constructs and the calendar days required for construct completion over time. Most CCA Sheets investigate the relationship between two kinds of CC data, with customizable filtering and classification of each, aiming to find correlation with successful cloning outcomes and/or to evaluate the effectiveness of particular cloning practices.

As an illustration, we created a CCA Sheet to investigate the relationship between storage time at 4 °C on the viability of DNA assembly mixtures. We hypothesized that prolonged storage before transformation might reduce colony counts or the likelihood of obtaining a sequence-verified clone. Storage prior to transformation is a routine part of our lab practices; therefore, of our 920 USER assemblies, >500 happened to be stored for ≥2 days before transformation, 267 stored ≥8 days, and even 92 samples stored for ≥51 days. We do not heat-inactivate the USER enzyme mixture (uracil DNA glycosylase, endonuclease VIII, and DpnI), leaving the possibility of DNA degradation during storage. Likewise, we have conducted 562 Golden Gate assemblies, 105 of which were stored for ≥18 days before transformation. No formal experiment needed to be conceived or implemented to collect this data set; rather, it was readily available because date and outcome information for these steps is logged in CC.

We created “CCA Transformation outcomes vs storage time” to analyze USER and Golden Gate assembly data. Storage duration can be found by comparing assembly and transformation dates. A ballpark estimate of colony counts is logged for each transformation as part of the “plate scoring” task, and the entries are structured into specific categories (e.g., 10 to 50 colonies) by selection from a dropdown menu. The existence of a sequence-verified miniprep arising from an assembly is annotated in that Assembly’s Status field, because the miniprep ID and its sequencing outcomes are linked to the assembly ID. The compiled data allow construction of stacked bar plots that relate storage duration to colony counts and to successfully obtaining the target plasmid (**Figure 4a**). Our data shows little dependence of colony count or eventual success on storage time for either USER or Golden Gate assemblies under our practices, with a modest trend toward lower colony count with longer storage time. Others can easily use this CCA Sheet with their own CC instance to determine whether their practices give similar results.

**Figure 4.**
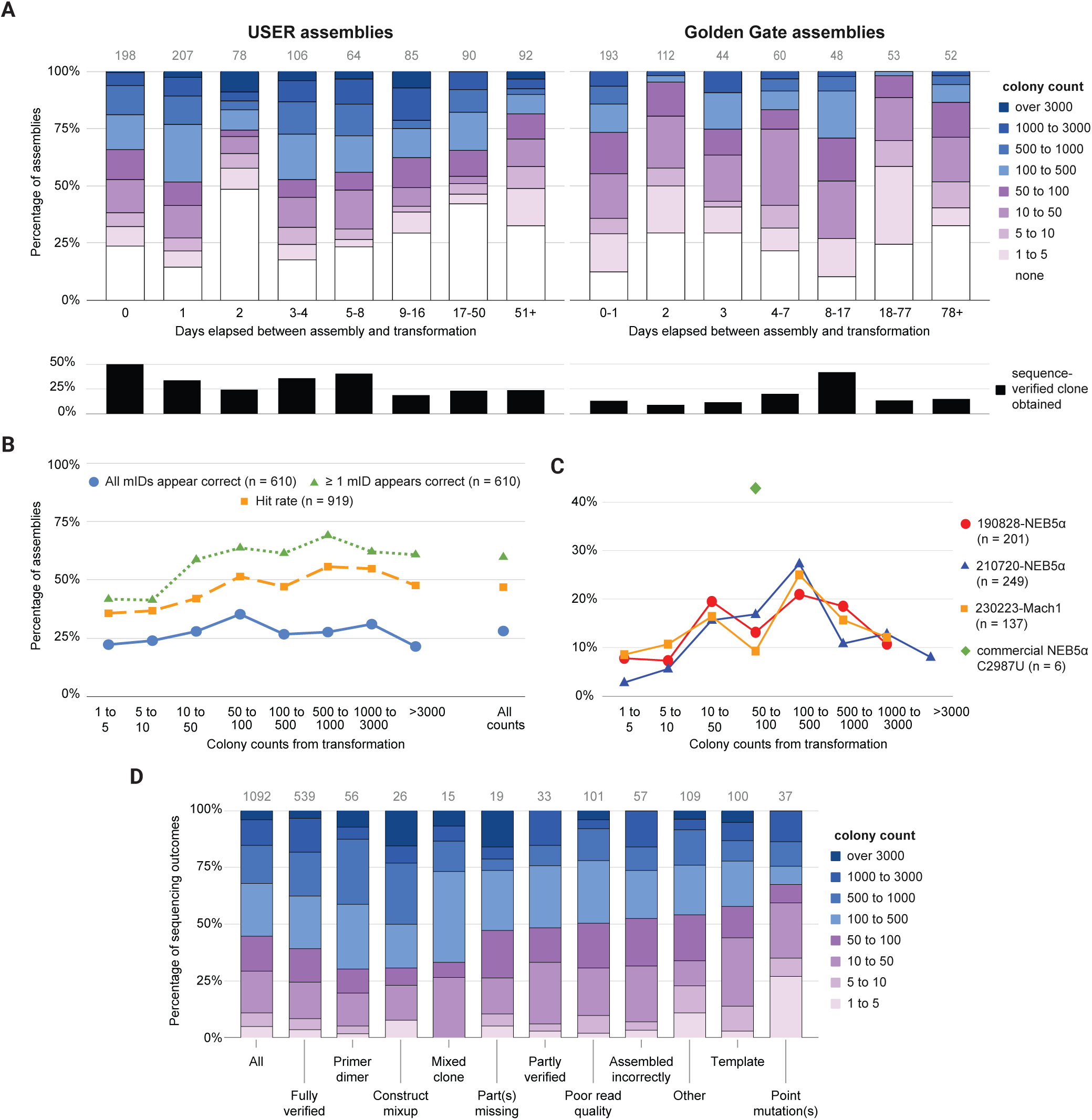
CC Analytics Sheets provide customizable analyses of CC data to inform cloning practices. The plots shown are generated by the indicated CCA Sheets reading from the Thuronyi Lab 2019-2025 CC Instance (**Supplementary Data**) and are only lightly edited for clarity and to fit the figure’s layout. **A, CCA Transformation outcomes vs storage time** output for USER assemblies (left) and Golden Gate assemblies (right). The top stacked bar plots show colony count distributions obtained from transforming assemblies (across all methods and conditions) that had been previously stored at 4 °C for the indicated durations. Colony counts estimates are logged in CC using the quasi-logarithmic categories shown in the legend at right. Colony counts above and below 100 are shown in shades of blue or purple respectively. The bottom plots show the percentage of the assemblies above that eventually yielded a sequence-verified clone. Counts of assemblies contributing to each bar are shown at the top in gray. In the CCA Sheet, the histogram bins update automatically to reflect the data distribution of the linked CC instance. **B and C, CCA Miniprep outcome vs colony count** outputs, reporting data for USER assemblies (99%) and Gibson assemblies (1%) that resulted in at least one miniprepped colony. **B** shows the percentage of *n* total assemblies for which all (blue circles) or at least one (green triangles) clone appears to be correct, based on sequencing and agarose gel electrophoresis data, as a function of colony count (left) or across all assemblies (far right). The estimated empirical hit rate (orange squares) is the percentage of assessed clones that appear to be correct for each of *n* assemblies; this accounts for variable numbers of clones assessed. No filters were applied. The corresponding data table is shown in **Supplementary Figure 11**. **C** shows the percentage of *n* total assemblies giving the indicated colony count ranges after transformation using 3 different batches of in-house prepared chemically competent cells (see Methods) and one commercial product (New England Biolabs product C2987U). Because we reserve the commercial cells for retransformation of low-colony-count assemblies, usage is low and peaks at moderate colony count. Because only assemblies that resulted in minipreps are included, there is no category for no colonies. Percentages based on ≥5 assemblies are shown on the plot; no other filters were applied. The corresponding data tables are shown in **Supplementary Figure 12**. **D, CCA Sequencing outcomes vs colony count** output, reporting data for USER assemblies. Each sequencing submission (Sanger or whole-plasmid nanopore) are logged in CC with read interpretation categories such as those shown on the x-axis. The colony counts for the transformation leading to each sequenced clone are shown in the stacked bar plot. The 1117 sequencing submissions in the data set represent 690 unique clones from 514 unique USER assemblies. Categories with ≥10 observations are shown on the plot. The “All” category combines the submissions shown in the other categories. Categories are further described in Methods. Bar colors and observation counts are as in panels A and B.

An important decision in cloning is how many colonies one should evaluate to eventually obtain a sequence-correct plasmid clone. In principle, high-fidelity PCR and high-efficiency assembly methods can give transformation plates where nearly all colonies are identical and correct – that is, the “hit rate” is near 100% – so that checking more than one is a waste of resources. However, hit rate can vary greatly, and usually multiple colonies must be checked to successfully obtain at least one correct clone. If the hit rate is low, checking even dozens of colonies is futile. In the absence of reliable phenotypic screening, it is very useful to be able to estimate the hit rate for a given assembly before choosing how many colonies to examine.

We considered that colony count might be correlated with hit rate and investigated this for our cloning so far using the “CCA Miniprep outcomes vs colony count” Analytics Sheet. Colony count is easy to assess, and intuition suggests it should be directly related to assembly efficiency, since the number of “correct” colonies should depend on the amount of correctly assembled DNA. On the other hand, “incorrect” colonies can come from a variety of genetic sources, and how these vary with total colony count is unclear. To estimate hit rate from available CC data, we wished to use as much available information about each transformed assembly as possible. Sequencing provides the least ambiguous data on clone correctness but is relatively expensive. CC supports a variety of workflows and strategies besides sequencing for checking colonies. We typically miniprep plasmid from 3-4 phenotypically correct colonies, assess the DNA by agarose gel electrophoresis (where migration of the primary supercoiled plasmid band strongly correlates with plasmid size, see **Supplementary Figure 9**), and follow up with sequencing. Sequencing results allow classification of unsequenced clones having similar or different gel results as probably correct or incorrect and let us analyze many more clones than we sequence. We initially focused our analysis on our primary assembly method, USER, and our speculative-miniprep- followed-by-gel workflow.

Our results (**Figure 4b)** show that, perhaps surprisingly, the success rate of our cloning practices so far seems to be largely independent of colony count. We can consider any assembly that yields at least one correct clone (across the combined assembly, transformation, and clone assessment processes) to be successful. 60% of assemblies on average met this criterion, with the percentage dipping only modestly lower at very low colony counts. Only 28% of assemblies gave only presumed- correct clones, with little variation across colony count. We see a modest positive trend in empirical hit rate (estimated number of presumed correct clones as a percentage of those checked, across all assemblies) with increasing colony count. Our average empirical hit rate estimate (47%) validates our initially arbitrary practice of checking 3 to 4 clones per assembly, which would give a 85% to 92% chance on average of obtaining a correct clone at the average hit rate. Contrary to our expectations, the data do not suggest that we should vary the number of colonies we check based on the total colony count for these assemblies.

The colony count statistics in “CCA Miniprep outcome vs colony count” can also be filtered by date, showing that our overall cloning success rate was higher in the first third of assemblies we conducted than in the remainder; or by assembly inputs, showing, as expected, that USER assemblies with 3 or 4 parts are less successful and produce fewer colonies than those with 1 or 2, and that including annealed oligos as an assembly component modestly increases the hit rate (**Supplementary Figure 10**). Filtering by competent cell batch and plotting the overall colony count distribution across all transformed assemblies (**Figure 4c**) allows comparison of competence under actual cloning conditions.

We also wondered whether incidence of certain assembly or transformation failure modes might be correlated with colony count. In particular, template contamination, where a PCR template with the same resistance marker as the target construct survives DpnI digestion and gives rise to a colony, and/or primer dimer insertion, where a primer heterodimer is elongated by DNA polymerase and assembles in place of the intended PCR product, could each be expected to inflate total colony counts and reduce the hit rate. CC sequencing classifications – entered during the “sequencing interpretation” task and limited to a discrete set of options – allow us to examine this correlation (**Figure 4c**). We found that primer dimer insertion did indeed skew toward higher colony count as compared to sequencing of correct constructs, while template contamination was actually somewhat more common at lower colony count. Interestingly, point mutations were substantially enriched at very low colony count, perhaps because a high mutation rate can inactivate plasmid replication or antibiotic resistance. Mixed clones (where sequencing showed the presence of two or more species differing at some bases, suggesting a multiple transformant colony) were, as expected, enriched at higher colony counts.

We do not intend to present these results as dispositive about cloning practices in general, or to suggest that these specific analyses are the most important or impactful ones that could be done. Our point is that CC-captured data can easily reveal to adopters actionable trends in their own cloning that are typically invisible when working at the level of individual constructs. Each CCA Sheet described here can be copied and connected to a CC instance to immediately provide the same analysis results, and additional analytics can be developed using CC data structures following these examples.

## Discussion

We believe CloneCoordinate can benefit a wide constituency of scientists, in training and professional, who build DNA as part of their research. It can make cloning practical in contexts where it would otherwise be prohibitively challenging; expand its accessibility, inclusion, and developmental value; and increase productivity and efficiency.

Our own experience relying exclusively on collective DNA construction using CC in our undergraduate research group over the past ∼6 years has demonstrated its value to us. Despite structural challenges such as low average participant experience and frequent competing time commitments, we have completed about 400 constructs. We invested an estimated average of 11 person-hours of active work for each construct completed, a significant fraction of which also contributed to ∼400 as-yet incomplete targets and ∼200 targets abandoned due to changes in research direction. We carried out this work while each of our group’s members (with the exception of the faculty advisor) developed their cloning experience from minimal to moderate during the course of their participation, while the group functioned productively as a collective unit of ∼4-7 students at a time. We believe CC was absolutely required for this coordinated approach, which in turn was essential for our productivity. It also enabled the involvement in DNA construction of a much larger number of trainees, at a lower average experience level, than would otherwise be practical: 34 students, including 10 completing undergraduate theses and 19 working full-time during the summer. We believe that hands-on exposure to DNA building greatly enhances understanding of, and is a powerful inducement to further participation in, synthetic biology and life sciences research.

We also predict that CC will enhance the productivity of experienced, full-time DNA builders, since it offers experiment registration and tracking, build phase abstraction, task batching, and standardized inventory and documentation. These capabilities provide time and effort multipliers even for an individual using CC alone.

To facilitate its adoption and accessibility, we aim to make CC maximally customizable and transparent without requiring users to recode its formulas. To support diverse cloning practices, we provide Settings (**Supplementary Text 2**) that can fit CC’s task management logic to many common workflows (for example, verification steps such as analytical PCR or sequencing can be mandated and their timing relative to minipreps adjusted). We invite feedback and development requests (as well as volunteer development partners), particularly from groups whose practices are less well accommodated by CC. To document how CC works, we provide a guide (**Supplementary Text 1**), logic flow charts (**Supplementary Table 1**) and explanations for each Setting (**Supplementary Text 2**).

The current implementation of CC is not yet performance-optimized, so instances with many thousands of entries become less responsive. CC is best suited to cloning throughputs of <100 constructs per month. Large or very high-throughput groups might opt for multiple parallel CC instances, e.g., divided by project area. We aim to improve performance in future versions and provide tools to assist periodic data archiving and migration of active and frequently-used entries to a fresh CC instance. We have created scripts to import all existing CC data into a new CC instance, enabling version updates.

The open source code, native spreadsheet format, and standard data structures of CC make it possible for anyone to develop and share compatible accessory Sheets, scripts, or software, and to easily access raw data. We have provided examples through our CC Accessory Sheets and documented our own coding conventions and practices (**Methods, General Sheets coding practices**). Depending on interest, tools could be readily created to convert the build instruction outputs of design tools such as j5, Benchling, or Geneious to CC queueing; to import and structure instrument data for the appropriate CC fields; and/or to export CC information to other formats, such as robot instructions for individual cloning tasks.

We exercise no control over users’ CC data or instances, though they are subject to Google’s terms of service, privacy policy, and server availability. Google maintains version histories and supplementary offline data backup as a CSV file is straightforward. We anticipate creating tools for automatic data anonymization (removal of sequences, descriptions, and construct names), unique ID assignment, and export to a standardized format, allowing CC users to share their cloning results with the broader community and aggregate them into large and general datasets.

We have established a paradigm for CC data collection and its real-time analysis using a small set of CC Analytics Sheets, which we aim to expand and continue to share freely. We will also focus on implementing algorithmic, data-driven recommendations for cloning troubleshooting through the CC Construct Tracker tool. Ideally these will be based on, or even automatically update from, analysis of CC data. Ultimately, CC’s purpose is to facilitate DNA assembly and improve its efficiency. We anticipate that the standardized data collection that CC enables will make this increasingly possible as the tool is adopted.

## Author Contributions

**Conceptualization:** B W. Thuronyi.

**Data curation:** Ethan Jeon and B W. Thuronyi.

**Formal analysis:** Ethan Jeon, Ziyang Shen, and B W. Thuronyi.

**Funding acquisition:** B W. Thuronyi.

**Investigation:** Ethan Jeon, Ziyang Shen, Santiago Christ, Evelyn Qi, Ida Fan, Nawon Lee, Odysseas Morgan, Madeline Ohl, Dylan Millson, Michelle Laker, Aspen Pierson, Emily Villegas Garcia, Yifan Zhang, Adeline Choi, Ashrita Iyengar, Rebecca Kim, Josh Lee, Linden Niedeck, Vanessa Oeien, Maliha Rashid, Nandini Seetharaman, Arnav Singh, Delaney Soble, Jenny Yu, Katherine Yu, Simms Berdy, Ellia Chang, Robin Kitazono, Sofija Ortiz, Dylan Taylor, and B W. Thuronyi.

**Methodology:** Ethan Jeon, Ziyang Shen, and B W. Thuronyi.

**Project administration:** B W. Thuronyi.

**Resources:** B W. Thuronyi.

**Software:** Ethan Jeon, Ziyang Shen, Santiago Christ, Evelyn Qi, Ida Fan, Nawon Lee, and B W. Thuronyi.

**Supervision:** B W. Thuronyi.

**Validation:** Ethan Jeon, Ziyang Shen, Santiago Christ, Evelyn Qi, Ida Fan, Nawon Lee, Odysseas Morgan, Madeline Ohl, Dylan Millson, Michelle Laker, Aspen Pierson, Emily Villegas Garcia, Yifan Zhang, Adeline Choi, Ashrita Iyengar, Rebecca Kim, Josh Lee, Linden Niedeck, Vanessa Oeien, Maliha Rashid, Nandini Seetharaman, Arnav Singh, Delaney Soble, Jenny Yu, Katherine Yu, Simms Berdy, Ellia Chang, Robin Kitazono, Sofija Ortiz, Dylan Taylor, and B W. Thuronyi.

**Visualization:** Ethan Jeon, Ziyang Shen, Santiago Christ, Evelyn Qi, Ida Fan, and B W. Thuronyi.

**Writing - original draft:** B W. Thuronyi.

**Writing - review & editing:** Ethan Jeon, Ziyang Shen, Ida Fan, Odysseas Morgan, Madeline Ohl, Aspen Pierson, Adeline Choi, Arnav Singh, Jenny Yu, Robin Kitazono, Dylan Taylor, and B W. Thuronyi.

## Acknowledgments

We thank Williams College and Thuronyi Lab alumni Michelle Garcia ‘21, Abu Sangare ‘23, Xavi Segal ‘23, and Russell Blakey ‘23 for their involvement in the group and feedback on CC. We acknowledge Matthew Baya, Gerol Petruzella and Tattiya Maruco of the Williams College Office of Information Technology for Apps Script and web hosting advice. B Thuronyi thanks Dr. Mea Cook for coworking and advice. This work was supported by Williams College and by NSF RUI Award 2401332 to B Thuronyi.

## Sources of graphical elements for Figures

Elements not cited are the authors’ original work.

**Figure 1**

NIAID Visual & Medical Arts. (10/7/2024):

DNA. NIAID NIH BIOART Source. bioart.niaid.nih.gov/bioart/123 Petri Dish. NIAID NIH BIOART Source. bioart.niaid.nih.gov/bioart/404

Eppendorf Tube. NIAID NIH BIOART Source. https://bioart.niaid.nih.gov/bioart/143 Next Gen Sequencer. NIAID NIH BIOART Source. bioart.niaid.nih.gov/bioart/386 Plasmid. NIAID NIH BIOART Source. bioart.niaid.nih.gov/bioart/411

Cryo Blood Vial. NIAID NIH BIOART Source. bioart.niaid.nih.gov/bioart/87

Google Sheets logo: https://commons.wikimedia.org/wiki/File:Google_Sheets_2020_Logo.svg Calendar by lailistudio fromNoun Project (CC BY 3.0). https://thenounproject.com/browse/icons/term/calendar/

Falcon tube by Alexander Bates, scidraw.io, https://10.0.20.161/zenodo.4421153 PCR thermocycler by Manon Chauvin.

https://commons.wikimedia.org/w/index.php?title=File:Thermocycler_T100.jpg&oldid=572865443

**Figure 2**

PCR tube strips by DataBase Center for Life Science (DBCLS). https://commons.wikimedia.org/w/index.php?curid=116852479 Plasmid with insert by Michael Jeltsch. https://commons.wikimedia.org/w/index.php?curid=51664406 Lab notebook icon by monkik from Noun Project (CC BY 3.0). https://thenounproject.com/browse/icons/term/lab-notebook/

## Materials and Methods

### General Sheets coding practices

Formulas use lowercase for built-in functions (these are bolded in this Methods text) and uppercase for custom-defined Named Functions. Direct references to ranges are in uppercase. Where possible and practical, Named Ranges (e.g., Registry_Construct_name) are used instead of direct references (Registry!C:C) for clarity. Sheets has no native code commenting capabilities, so formulas are commented, where possible, using comment clauses within formulas. These consist of FALSE,”comment text”, pairs in **ifs** or comment1,”comment text” pairs in **let**. Any formula can be commented by wrapping it in **let**(comment,”comment text”, (formula)).

Some function choices are preferred in CC code for performance reasons. Lookups by ID (e.g., “a20”) preferably use **index** and the Named Function ID_TO_INDEX, which converts the ID to a row number by extracting the serial number (20) and adding the value of the Named Range settings_NumHeaderRows (2 in CC v1.0). **Index**/**xmatch** is an acceptable alternative and is used for indexing by values other than ID. **Filter** is strongly preferred over **query** for performance reasons to retrieve multiple matches. **Ifs** or nested **if** statements are preferred because they are short-circuit evaluated in Sheets, while **and** and **or** are not. **Ifs** cannot be used with array inputs or outputs, but nested **if** is equivalent. **Countif** is preferred to **counta**/**iferror**/**filter** where equivalent. Where possible, formulas in each row that filter/match against an entire column are avoided and instead **regexmatch** or **search** a single helper cell on the “(Lookup lists and tabulations)” Sheet that compiles the filter/match output (however, the performance of these strategies has not been rigorously compared).

In general, use of helper columns is minimized in favor of nested formulas (possibly including array processing with **byrow** or **bycol**) because Sheets performance is influenced by Sheet size and total Spreadsheet size is capped. One exception is that dynamically populated dropdown menu choices must be drawn from a range of cells using Data Validation, so in this case, a helper range is used for each cell with a dropdown. Helper columns are also sometimes needed to avoid circular references, especially when evaluating Status formulas. Where practical, a single formula in the header row(s) is used to populate all rows in a Sheet by outputting an array. This allows users to view the entire output of the formula (even if it exceeds the cell dimensions) for a given row by selecting the appropriate cell, which would otherwise show only that row’s formula. This strategy is also used on Assist tabs to retrieve and display information from the relevant source tab. Where the ability to view the entire output is important but use of a single formula with an array output is cumbersome, a helper column is used that contains each row’s formula, and the user-facing column is populated with a single formula outputting the array of the formula column, e.g., ={Z:Z}.

### CC Data structures

Columns (and cells) in CC have a binary designation as either user data entry or formula output. User data entry columns have a blue (#c9daf8) or green (#d9ead3) header color and are part of a Named Range. These specific header colors are integral to the user data entry column definition and are referenced by Apps Scripts. For Sheets that track user-generated entities (e.g., Experiments, Registry, Oligos …), each user data Named Range spans an entire column so that added rows are automatically included. For this reason, the Named Range includes the header rows (the number of which is stored in Named Range settings_NumHeaderRows). User data is assumed to never contain formulas and is copied as values by Apps Scripts. Formula output columns have white column headers and are always populated by formulas (in the header row with an array output or with a formula in each row) below the header row. Formula output columns may or may not be defined as Named Ranges. When a formula output column is referred to by any formula outside its containing tab, it should be assigned and referred to using a Named Range, which greatly simplifies reference searching.

Accessory Sheets and the Apps Script-assisted workflow for data migration / version updating both rely on data import using **importrange** to retrieve the contents of user data Named Ranges. Therefore, Named Range names and data range definitions must remain unmodified by users. While new Named Ranges can be added by users in principle, name space collisions from future CC updates should be anticipated and avoided.

Full-column Named Ranges for user data entry are named with the tab name followed by a descriptive name for the column in sentence case with underscore separation (e.g., dsDNA_d_Certify_correctly_queued). Single-cell or irregularly sized Named Ranges on the Settings & admin tab are named with settings_ followed by the abbreviation for the affected tab and then a setting description in camel case (e.g., settings_dTemplateMiniprepMinVolumeLeft). These conventions facilitate Named Range selection in formulas through autocomplete.

### Accessory Sheet specification

CloneCoordinate Accessory or Analytics (CCA) Sheets are structured to allow user data and some formula data to be read from a connected CC instance. The Imported-data tab contains a cell where the CC URL can be entered, and a series of **importrange** formulas to import relevant data from that URL by Named Range. Each Imported-data column is itself designated as a Named Range with the same name as the range its formula imports from CC. Code that references Named Ranges is therefore portable between CC and CCA(s). The Imported-data tab can be regenerated, updated, or trimmed (to avoid importing data not actually referenced within the CCA Sheet) using Apps Script. If single-column or irregular Named Ranges (from Settings & admin) must be imported, these are manually referenced on an optional Additional imported data tab. All calculations specific to the CCA’s operation are housed in a Formulas tab which can read in full-column ranges from the Imported-data tab (e.g., “={dsDNA_d_ID}”) as needed. Separation of the Formulas and Imported-data tabs allows simple Apps Script changes to Imported-data, such as rebuilding to include newly added Named Ranges or streamlining to remove any that are unused by the CCA Sheet. Other tabs provide data visualizations based on Formulas calculations and user settings to modulate the output.

A new CCA can be created by making a copy of an existing one or of the CCA Template Sheet. All information flow is from CC to CCAs; CC itself uses **importrange** only on the Dashboard to display CC updates/news information from a public Sheet. CCAs should operate independently, never in chains.

### Apps Scripts

Optional scripts are supplied with CC to repair/replace/update the data validation and formatting, and/or the formulas, on all rows of a user data entry tab based on those used in the first row below the headers. This is important to repair problems sometimes caused by copy/paste or cut/paste operations, and to populate new rows added by users. A Migration script is also provided along with the update edition of new CC versions to facilitate copying of user data and settings from a CC instance already in use. The update edition is preset with **importrange** formulas in appropriate cells so that user data can be imported from a CC instance; the Migration script copies the imported values and writes them permanently to the new version.

The Authorization Scope for Apps Scripts attached to each CC Sheet was manually limited to the current document only using the “ * @OnlyCurrentDoc” JsDoc annotation in a file-level comment.^29^

### Software and User Support

Links to all software are provided at clonecoordinate.org and below. Software is based on the Google Sheets architecture and can be accessed only through Google infrastructure in a web browser, although offline editing can be set up once a copy is established. Users are invited to [join the Slack workspace](https://docs.google.com/forms/d/e/1FAIpQLSc_sZ0UCExPnGI9pnKqzHocbSLykopWCEJaKc_R6tPh0Ep0jA/viewform?usp=sharing&ouid=109298485705044436089) at clonecoordinate.slack.com to interact with the development team and other CC users.

#### CloneCoordinate

The Thuronyi Lab CloneCoordinate instance (version 0.99) containing cloning data from 2019-2025 can be viewed at the following link. Sequence information and construct descriptions have been removed to maintain privacy for ongoing unpublished research. https://docs.google.com/spreadsheets/d/1wZwXvLzPqq4h6Fzb-27fKORN6zZ60yFWyrfPe6eCVts

A link to a blank copy of CloneCoordinate can be obtained using this automated form (all questions are optional) or on request to: https://docs.google.com/forms/d/e/1FAIpQLSf4krmMykxG_7Nfxe9BOy0DYSe6lG_TXIzahH6ImPm4pvRDyw/viewform?usp=header

CloneCoordinate Issues (bug reporting, feature ideas and improvement plans) have been tabulated in the public GitHub repository https://github.com/bthuronyi/CloneCoordinate beginning in August 2023 and users are invited to contribute.

#### CloneCoordinate Accessory Sheets

An inventory of CCA Sheets with links to blank copies and copies linked to the Thuronyi Lab 2019-2025 CC Instance can be found at the following:

https://docs.google.com/spreadsheets/d/1Btno2VytKgC-cLb7Uxt_Vres0n_p1SWHupQ-D5l0nao

### Data analysis

#### Figure 4a, CCA Transformation outcomes vs storage time

“Assembly” analysis shown in Figure 4a is set to display data from USER assemblies; this can be changed to Gibson or Ligation types in the Settings tab of the CCA Sheet. To generate each chart, assembly dates, transformation dates, assembly Status, and colony counts are retrieved from CC. If multiple transformations occurred, only the most recent is included. The user Setting for “assembly type to display” causes the retrieved data to be **filter**ed by assembly type. Status records whether a sequence- verified miniprep is eventually obtained from a given assembly by including a “♥” character. Days elapsed between assembly and transformation are calculated. Histogram bins represent deciles of the days elapsed, calculated using **percentile** (to determine the bin cutoffs) and **frequency** (to create the histogram counts) formulas. Because time of transformation and assembly is recorded only to the nearest day, storage durations are quantized and will typically fall into fewer than 10 bins. To generate the plot x-axis category labels, the bins are parsed to account for ones that span multiple days elapsed. The last bin cutoff is not actually given by **percentile** but instead is displayed as the text of the largest bin cutoff from **percentile** plus 1 with a “+” appended, and **frequency** ignores this “bin” but produces an output alongside it corresponding to values larger than any of the other bin cutoffs.

The histogram bins are then used in **frequency** formulas operating on **filter**ed assemblies whose colony counts match each option in turn in the “Colony counts” dropdown menu. Currently these options are hard-coded into the CCA Sheet and will need to be updated if other options are made available. These frequencies are displayed as a normalized stacked bar plot.

#### Figure 4bc, CCA Miniprep outcome vs colony count

To generate data tables and charts, minipreps with an assembly as their origin are retrieved from CC. If a transformation ID is not referenced as the miniprep source, the ID most likely to be relevant is estimated by comparing the transformation dates for that assembly with the inoculation date of the miniprep and choosing the transformation with the largest serial number that occurred before that date. Colony counts and other parameters for the transformation assigned to each miniprep are then retrieved, along with assembly parameters for its source.

Minipreps are classified as appearing incorrect, ambiguous, or correct (in terms of their assembly process having put together all intended parts) based on agarose gel electrophoresis results and sequencing, aggregated across minipreps derived from each assembly. Gel band analysis is logged and processed in CC in the “Minipreps_m_gel_success” column. Tabulated values are compared with flags set in “settings_GelCodingQualityFlags” in Settings & admin. Briefly, the miniprep is considered likely correct if the gel lane shows a single band (or a supercoiled and relaxed band) and its position relative to the ladder is as expected for a plasmid of the targeted size. If the band is absent, smeared, clearly at the wrong size or multiple bands are present (not consistent with supercoiled/relaxed), the miniprep is considered likely incorrect. This classification is then adjusted based on sequencing results.

For miniprep sets with sequencing available, the sequencing alignment interpretation classification for the read of a given miniprep with the largest serial number is retrieved. If this indicates a problem (anything other than verification of some/all parts or read failure), the miniprep is presumed to be incorrect. In addition, if gel band interpretation alone suggested this and/or other minipreps from the same assembly were correct, and this is refuted by sequencing of any miniprep, the entire set is reclassified as incorrect. If a given miniprep was sequenced and found to be correct (fully sequence verified), it is classified as correct, but classifications for other minipreps based on gel band interpretation are not changed.

These determinations, wherever they can be made, are used to classify the minipreps derived from each assembly as all likely incorrect, ≥1 correct but not all correct, or all correct. These classifications are the input for the “Data set” tables in the CCA Sheet and are counted after being filtered by various user-customizable criteria.

The empirical hit rate is calculated for each assembly by dividing the number of presumed correct minipreps (or the number of sequence verified minipreps derived from the assembly, whichever is larger) by the total number of minipreps and converted to a percentage. Minipreps not classified as correct or incorrect contribute to the denominator for the calculation, but the hit rate is not calculated, and therefore does not contribute to the plots or the average, if any gel band data is ambiguous or if no gel band data is available.

#### Figure 4d, CCA Sequencing outcomes vs colony count

Sequencing interpretation classifications are retrieved from CC and those derived from minipreps which in turn were derived from assemblies are selected and their associated transformation information retrieved. If a transformation ID is not referenced as the miniprep source, the ID most likely to be relevant is estimated by comparing the transformation dates for that assembly with the inoculation date of the miniprep and choosing the transformation with the largest serial number that occurred before that date.

The selected sequencing interpretation classifications are further refined using **filter** and **query** to those that match the assembly types and procedures specified in the Settings tab. For each sequencing interpretation classification, the number of occurrences of each colony count classification is tabulated and used to construct a frequency table, the categories for which are then sorted by the fraction of transformations having ≤100 colonies. The plot shows a normalized histogram for the sorted frequency table.

The categories listed on the x-axis of the plot in Figure 4d are as follows: All, all sequencing submissions. Fully verified (“All parts verified present” in CC), sequencing that shows a clone is correctly assembled with all parts present and no detected errors. Primer dimer (“Primer dimer in place of part” in CC), sequencing shows that a primer homo- or heterodimer inserted into the clone in place of the intended PCR product. Construct mixup (“Mixup? Matches other map” in CC), sequencing corresponds to a target plasmid design other than the expected one, presumably based on inadvertent mislabeling or incorrect selection of samples or contamination at some point in the assembly process. Mixed clone (“Mixed clone: signs of 2 overlapping sequences for part, not all, of read” in CC), indications of a miniprep containing multiple distinct plasmids, likely due to a multiple transformation event. Part(s) missing (“Part(s) missing completely” in CC), misassembly involving absence of one or more parts (not template contamination). Partly verified (“Some parts verified present” in CC), determination that some parts are present and no errors are detected so far, but further sequencing is needed. Assembled incorrectly (“Evidence of misassembly” in CC), indication that parts are present but not assembled as desired, e.g., duplicated or in the wrong order or orientation. Other, special situation described in a free text field. Poor read quality (“Sequencing quality too low to use” in CC), sequencing quality too low to gain reliable information about the clone. Template (“Template or part with same marker, not assembled construct” in CC), sequenced plasmid is not the desired construct but is instead a PCR template or other part originally added to the assembly mix that has survived digestion or been reassembled intact. Point mutation(s), substitution mutations observed. CC also provides further categories not shown in Figure 4d due to low counts in the underlying data set: Mixup? Part present from source not listed as part of this construct; Primer dimer in addition to part; 1-2 base insertion(s) or deletion(s); Large (3+ base) insertion(s) or deletion(s); Transposable element present; PCR amplified undesired sequence instead of desired one.

### Cloning

#### Sample storage

DNA samples and bacterial strain cryogenic stocks were stored in Thermo Scientific EasyStrip Plus Tube Strips with Attached Flat Caps (thermofisher.com, cat. no. AB2000) arrayed in 96-position polypropylene racks with covers, such as from MUHWA (Amazon.com, cat. no. MH-90115TR5) or Axygen Scientific (cat. no. R-96-PCR-FSP). Each tube strip was labeled with a sample type abbreviation (o, d, a, g, or m), plate number, and column number. Oligonucleotide backup stocks were stored at -20 °C, all other DNA samples (including assembly mixtures) were stored at 4 °C, and cryogenic stocks at - 80 °C.

PCR tube strips (200 µL with attached caps) were purchased from Hyundai Micro Co. (Amazon.com, cat. no. HBP002CF8D). Microcentrifuge (Eppendorf) tubes were purchased from VWR (1.5 mL, Axygen Scientific, cat. no. 10011-702) or Fisher Scientific (2 mL, 30,000 x g centrifugal force tolerance, cat. no. 05-408-141). 96-well (cat. no. 204353-100) and 24-well (cat. no. 202061-100) deep- well plates were purchased from Agilent Technologies.

#### Chemicals and reagents

*Water.* For *in vitro* DNA handling, HyClone water (Cytiva cat. no. SH3053803) was used. Buffers were prepared using 18 MOhm resistivity ultrapure water (Thermo Scientific GenPure Pro UV-TOC, model no. 50131948) and media was prepared using deionized water, then sterilized by autoclave (125 °C, ≥30 min per L in the largest container) or filtration (0.2 µm aPES membrane, Nalgene cat. no. 565- 0020).

*Chemicals.* Ethidium bromide solution (10 mg/mL, cat. no. E1510), boric acid (cat. no. B7901), lithium acetate dihydrate (cat. no. L6883), DL-dithiothreitol (molecular biology grade, cat. no. D9779), betaine (5 M solution, cat. no. B0300), dimethyl sulfoxiode (cat. no. 276855), and guanidinium hydrochloride (98%, cat. no. 50950) were purchased from MilliporeSigma. Tris (tris(hydroxymethyl)aminomethane) base (cat. no. T60040), Tris-HCl (cat. no. T60040), sodium chloride (cat. no. S23020), agar (bacteriological grade, cat. no. A20030), SOB Broth (cat. no. S25000) and LB Broth (Lennox, cat. no. L24060) were purchased from Research Products International (rpicorp.com). Carbenicillin (Cb, cat. no. C-103), spectinomycin (Sp, cat. no. S-140), tetracycline hydrochloride (Tc, cat. no. T-101), chloramphenicol (Cm, cat. no. C-105), kanamycin sulfate (Km, cat. no. K-120), and deoxynucleotide triphosphates solution (D-900-10) were purchased from Gold Biotechnology. Agarose (cat. no. EM-2125) was purchased from VWR.

*Antibiotic stocks.* 1000x stocks were prepared as follows: Cb, 100 mg/mL in 50% ethanol (dissolved in 0.5 volumes water, then absolute ethanol added); Sp, 100 mg/mL in water; Tc, 100 mg/mL in ethanol; Cm, 30 mg/mL in ethanol; Km, 50 mg/mL in water. Stocks were 0.2 µm filter sterilized and stored at -20 °C or -80 °C for longer periods.

*Reagents.* Q5U Hot Start High-Fidelity DNA Polymerase (cat. no. M0515), USER enzyme (cat. no. M5505), DpnI (cat. no. R0176), BsaI-HFv2 (cat. no. R3733), T4 DNA ligase (cat. no. M0202), Gibson assembly mix (cat. no. E2611), NEBuilder HiFi DNA Assembly Master Mix (cat. no. E2621), and ET SSB (cat. no. M2401) were purchased from New England Biolabs. PhusionU Hot Start High-Fidelity DNA Polymerase (cat. no. F555L) was purchased from ThermoFisher Scientific.

#### Bacterial strains and plasmids

Routine cloning was carried out using NEB 5-alpha (New England Biolabs, cat. no. C2987) or Mach1 T1^R^ (ThermoFisher Scientific, cat. no. C862003) *E. coli* strains.

Rather than reporting new DNA constructs, this work describes a general method for cloning, so the attached CC data set does not contain specific sequences.

#### Double-stranded DNA

##### Oligonucleotides

Oligos were ordered from Integrated DNA Technologies at 25 nmol scale, shipped dry in PCR plates normalized to 5 or 10 nmol per well. Oligos were resuspended at 50 µM concentration in **storage buffer** (10 mM Tris-HCl, pH 8.1) to create a primary stock which was stored at -20 °C in storage tube strips. A working stock at 10 µM concentration in storage buffer was created from the primary stock and stored at 4 °C in storage tube strips.

##### Oligonucleotide annealing

6 µL of each oligo was mixed with 8 µL of 2.5x annealing buffer (50 mM NaCl + 10 mM Tris-HCl, pH 8.1) giving a 20 µL solution containing 20 mM NaCl, 10 mM Tris-HCl, pH 8.1, and 3 µM each oligo. This solution was heated to 95 °C and cooled to 30 °C at a rate of 0.2 °C/sec using a PTC-200 Thermal Cycler (MJ Research), then diluted 50-fold with storage buffer and stored at 4 °C.

##### PCR

PCR mixtures contained 3 µL each primer (600 nM final), 33.5 µL of the appropriate premix, 0.5 µL DNA polymerase, approximately 0.5 µL of the indicated DNA template, and 10 µL of water and/or additives for a total volume of 50 µL in PCR tube strips. Premixes contained 1.5x manufacturer polymerase buffer and 300 µM each dNTP (giving 1x and 200 µM final concentrations). Additives included 1–2.5 µL DMSO (2–5% final), 0.5 µL extreme thermostable single-stranded DNA binding protein (New England Biolabs M2401), or 5x CES PCR enhancer^30^ (with the 5x stock containing 2.7 M betaine, 6.7 mM DTT, 6.7% DMSO and 55 µg/mL BSA, prepared from solutions of 5 M betaine, 200 mM DTT, 20 mg/mL BSA, and neat DMSO and stored at -20 °C). Thermal cycling was carried out using a PTC-200 Thermal Cycler (MJ Research) with a default program of 98 °C initial denaturation for 30 seconds followed by 30 cycles of 98 °C for 15 seconds, 63 °C annealing for 30 seconds, 72 °C elongation for 15 seconds per kb of the longest PCR product, then a final elongation of 72 °C for 5 minutes and a hold at 12 °C. Actual annealing temperatures, cycle counts, and elongation times are recorded for each PCR in CC. PCRs were stored at 4 °C after thermal cycling prior to purification.

##### Purification of PCR products

PCR products were diluted 5-fold with **binding buffer** (44.5 wt% GuHCl, 22.0 wt% isopropanol, 33.5 wt% MilliQ water) or buffer PB (Qiagen cat. no. 19066) and loaded onto silica spin columns (Epoch Life Sciences Econospin, item 1910, or GenCatch, item 2160). Vacuum was applied and the columns were washed with 0.7 mL **ethanol wash buffer** (1.21 wt% 1 M Tris pH 7.4, 74.1 wt% absolute ethanol, 24.7 wt% MilliQ water) or buffer PE (Qiagen 19065), then centrifuged for 1 min at 21,000 x g to remove residual wash buffer, and eluted with storage buffer or buffer EB (Qiagen cat. no. 19086), typically 50 µL, again by centrifugation as before.

#### Assemblies

Actual volumes, concentrations, and assembly scales are recorded for each assembly directly in CC. The following procedures were typically used:

##### USER assemblies

Assemblies were typically prepared at 10 µL scale as follows (“USER - Standard v1” conditions): dsDNA fragments and HyClone water were mixed in approximately equimolar ratios to a final concentration of 7.5 ng/kb/µL (1 ng/kb/µL = 1.623 nM) and 5 µL volume. A 2x premix consisting of 2.5 µL HyClone water, 1 µL 10x Cutsmart buffer (New England Biolabs cat. no. B6004), 0.75 µL DpnI, and 0.75 µL USER enzyme mix was prepared and 5 µL added to each assembly. For the “USER - Extra DpnI v1” condition, 1 µL DpnI and 0.5 µL USER enzyme per assembly were used in the premix. The mixture was incubated at 37 °C in a thermal cycler with heated lid for 60 minutes, then stored at 12 °C or 4 °C until transformation. Actual volumes, concentrations, and assembly scales are recorded for each assembly in CC.

##### Gibson assemblies

Mixtures were prepared as above but with 5 µL 2x Gibson assembly mixture rather than the USER premix, and were incubated at 50 °C for 60 minutes.

##### Golden Gate assemblies

Mixtures were prepared as above but with 1 µL 10x T4 DNA ligase buffer (New England Biolabs, cat. no. B0202SVIAL), 0.5 µL T4 DNA ligase (400 U/µL) and 0.5 µL type IIS restriction enzyme (usually BsaI-HFv2) added directly to the mixture of parts rather than as a premix. The mixture was placed in a thermal cycler and subjected to 60 (“GGATE1” or “GGATE60”) or 120 cycles of 37°C for 5 minutes and 16°C for 5 minutes, followed by 37°C for 1 hour, and 80°C for 10 minutes, or incubated statically on the benchtop (“GGATEBENCH”) or at 37 °C or 25 °C as indicated in CC.

#### Transformations and competent cells

##### Preparation of chemically competent cells

A culture of the appropriate *E. coli* strain was grown overnight in LB medium at 37 °C with shaking (225 rpm), then diluted 50-fold in a baffled shake flask and grown to late log phase (OD600 0.3 - 0.5) either entirely at 37 °C or shifted for ∼2 hours to room temperature before collection. Cells were cooled on ice then centrifuged at 4 °C and 4,500 x g for 15 minutes in a Thermo Scientific Sorvall Legend XTR centrifuge. The supernatant was removed and the pellet was resuspended in 1 volume of ice cold LB medium per 20 mL initial culture volume. To 1 volume of the suspension was added 1 volume ice cold **2x TSS solution**^31^ (200 g/L PEG 8000, 12 g/L magnesium chloride, 10 vol% DMSO, 20 g/L LB medium) and mixed by inversion. The resulting 1x TSS cell suspension was separated into 0.5 mL portions, snap frozen in liquid nitrogen, and stored at -80 °C. Before use, 1 volume (0.5 mL) of **KCM solution** (100 mM potassium chloride, 30 mM calcium chloride, 50 mM magnesium chloride) was added to each tube of cells on ice. 100 µL of the resulting suspension was used for each transformation.

##### Transformation

Specific volumes, recovery media, recovery times and amounts plated are recorded for each transformation in CC. A typical transformation using the “standard” procedure was as follows. 5 µL of assembly mixture were mixed with chemically competent cells in a PCR tube strip and incubated on ice for 2-3 min, then heat shocked at 42 °C for 90 (labmade TSS cells) or 30 (New England Biolabs cells) in a thermal cycler and returned to ice for 2 minutes. The cells were recovered by adding the mixture to 400 µL SOC (3.603 g/L D-glucose, 0.186 g/L potassium chloride, 2.4 g/L magnesium sulfate, 0.5 g/L sodium chloride, 20 g/L casein enzymatic hydrolysate) for 60 min at 37 °C in a 96-well deep well plate. 150 µL of the suspension was spread using sterile glass beads on ∼15-30 mL LB agar containing the appropriate antibiotic in a 100x15 mm Petri dish and incubated at 37 °C overnight.

For the “maximum care” procedure, cells were incubated on ice for 30 min after addition of DNA but before heat shock. The recovery was carried out by adding 1 mL SOC to the cell suspension, then the entire mixture was centrifuged for 6 min at 3,000 x g at room temperature and most of the supernatant discarded. The cell pellet was resuspended in the remaining medium (∼50 µL) and spread on a plate.

#### Isolation of plasmid DNA (miniprep)

##### Inoculation and culture growth

24-well plates containing 4 mL LB medium per well with the appropriate antibiotic were prepared and fitted with breathable sealing film (Breathe-EASIER, Diversified Biotech, cat. no. BERM-2000, or 3M Micropore Surgical Tape, cat. no. 1530-3). Colonies from transformation plates were picked using sterile toothpicks or pipette tips. Inoculated cultures were grown for 16 - 48 hours at 37 °C with shaking (225 rpm). Plates were centrifuged at 2250 x g for 10 min in an Eppendorf 5804R centrifuge equipped with a swinging bucket rotor. The supernatant was discarded by inversion into a waste container and blotting on paper towels. 24-well plates were reused by soaking in 10-fold diluted household bleach overnight, then rinsing and cleaning in a laboratory dishwasher (Labconco FlaskScrubber) and autoclaving.

##### Plasmid purification

Pellets were resuspended in 250 µL buffer P1 (Qiagen cat. no. 19051) containing RNAse A (100 µg/mL, New England Biolabs cat. no. T3018), then transferred to 2 mL high- speed microcentrifuge tubes. Cells were lysed and the lysate neutralized with Qiagen buffers P2 (cat. no. 19052) and N3 (cat. no. 19064) according to the manufacturer’s instructions. The tubes were spun for 6 min at 30,000 x g in an Eppendorf 5430R microcentrifuge and the supernatant loaded on QIAPrep Spin 2.0 silica columns (cat. no. 27115) affixed to a vacuum manifold. Columns were washed with 0.5 mL binding buffer (44.5 wt% GuHCl, 22.0 wt% isopropanol, 33.5 wt% MilliQ water) or buffer PB (Qiagen cat. no. 19066), then with 0.7 mL ethanol wash buffer (1.21 wt% 1 M Tris pH 7.4, 74.1 wt% absolute ethanol, 24.7 wt% MilliQ water) or buffer PE (Qiagen cat. no. 19065), then centrifuged for 1 min at 21,000 x g to remove residual wash buffer, and eluted with storage buffer or buffer EB (Qiagen cat. no. 19086), typically 100 µL, again by centrifugation as before, and transferred to storage tube strips.

#### Quality control and DNA analysis

##### Agarose gel electrophoresis

1% agarose gels were prepared in lithium acetate borate buffer (1x LAB, prepared from a 25x stock containing 250 mM boric acid and 250 mM lithium acetate)^32,33^ containing 5 µg/mL ethidium bromide and run at 250 V, typically for 30 min, in a HR-2525 High-Resolution/High- Throughput Electrophoresis Box (IBI Scientific cat. no. IB57000). Gels were imaged using a SynGene G:Box system with UV transillumination.

##### Spectrophotometric DNA quantification

DNA concentration was determined spectrophotometrically using a Biotek Synergy H1 plate reader fitted with a BioTek Take3 plate (Agilent), or with a Thermo Scientific NanoDrop 2000 Spectrophotometer, according to the manufacturer’s instructions.

##### Analytical PCR

PCRs (20 µL) were prepared in an ice-cooled aluminum block and were composed of 10 µL Taq Mastermix (Syd Labs cat. no. MB067-EQ2G), 1 µL of each 10 µM primer stock, 0.3 µL of colony stock or miniprep template, and 7.7 µL of Hyclone water. Thermal cycling was carried out using a PTC-200 Thermal Cycler (MJ Research) with 95 °C initial denaturation for 30 seconds followed by 30 cycles of 95 °C for 30 seconds, 55 °C annealing for 30 seconds, 68 °C elongation for 3 minutes, then a final elongation of 68 °C for 5 minutes and a hold at 12 °C. Analytical PCR products were analyzed by agarose gel electrophoresis.

##### Sequencing

Sanger (Quintara Biosciences) or nanopore (Plasmidsaurus) sequencing was prepared as shown in CC and according to the provider’s instructions and submitted by overnight mail.

#### Strain storage

Sequence-verified plasmids were transformed into *E. coli* using labmade chemically competent cells according to the procedure described above. For high-throughput plating, 10 µL of the recovery mixture was transferred by multichannel pipette and spread by gravity onto LB agar strips (about 3.6 mL) with antibiotics prepared in 8-channel polypropylene reservoir plates (Agilent, cat. no. 204367-100). After overnight growth, a single colony from each channel was inoculated in 1 mL LB medium with appropriate antibiotic and grown overnight in a 96-well deep-well plate fitted with breathable sealing film at 37 °C with shaking at 225 rpm. 300 µL of 60 wt% glycerol was added and 200 µL of the mixture transferred to replicate working and backup storage tube strips for preservation at -80 °C. 8-channel reservoirs were reused by scraping out the agar strips with a spatula, then soaking in 10-fold diluted household bleach overnight, rinsing and cleaning in a laboratory dishwasher (Labconco FlaskScrubber) and autoclaving.

