## Supplementary Information for "CloneCoordinate: Open-source software for collaborative DNA construction"

**Design — Constructs 1 and 2 conception & design**

| | | |
|---|---|---|
| **Tab** | Experiments | |
| **Sample ID** | e1 | |
| **Type** | control | test |
| **Draws on** | r1 | r2 |

**Queueing phase — Construct 1 (r1) via Gibson assembly**

| | | | | | | | | | | | | | | |
|---|---|---|---|---|---|---|---|---|---|---|---|---|---|---|
| **Tab** | Registry | Oligos o | | | | dsDNA d | | | | | | Assemblies a | | |
| **Sample ID** | r1 | o1 | o2 | o3 | o4 | d1 | | | d2 | | d3 | a1 | | |
| **Type** | construct | primer | | | | PCR | | | anneal oligos | | digest | Gibson | | |
| **Draws on** | external design tools | external design tools | | | | o1 | o2 | m20 | o3 | o4 | m21 | d1 | d2 | d3 |

**Queueing phase — Construct 2 (r2) via Golden Gate assembly**

| | | | | | | |
|---|---|---|---|---|---|---|
| **Tab** | Registry | Golden Gate g | | | | |
| **Sample ID** | r2 | g1 | | | | |
| **Type** | construct | part donor plasmid | | | PCR | gBlock |
| **Draws on** | external design tools | r3, m4 | r4, m5 | r5, m6 | r6, d4 | r7, o5 |
| | | Golden Gate Assembly Queuer | | | | |

**Correct clone isolation phase — Construct 1 (r1) straight to minipreps**

| | | | | | | | | | |
|---|---|---|---|---|---|---|---|---|---|
| **Tab** | Transformations t | *skip optional analytical PCR step* | Minipreps m | | | Sequencing | | | Strains |
| **Sample ID** | t1 | | m1 | m2 | m3 | serial # | | | s1 |
| **Type** | chem comp | | silica spin column | | | Sanger | Sanger | Sanger | cryostock |
| **Draws on** | a1 | | t1 | | | m1, o5 | m2, o5 | m2, o6 | m2 |

**Correct clone isolation phase — Construct 2 (r2) analytical PCR on colony stocks**

| | | | | | |
|---|---|---|---|---|---|
| **Tab** | Transformations t | Analytical PCR | Minipreps m | Sequencing | Strains |
| **Sample ID** | t2 | serial # | m7 | serial # | s2 |
| **Type** | electroporation | | magnetic bead (SPRI) | nanopore | cryostock |
| **Draws on** | g1 | t2 colony stocks 1-8 o7, o8 | t2 colony stock 6 | m7 | m10 |

**Analysis / quality control steps**

| | | | |
|---|---|---|---|
| **Tab** | dsDNA d | Minipreps m | Analytical PCR |
| **Type** | Agarose gel electrophoresis | | |
| | UV-vis concentration measurement | | |

**Supplementary Figure 1. Example construct queueing diagram.** The diagram shows how two new constructs for a simple experiment might be queued in CC, including example sample identifiers. Both constructs are first designed using existing tools of the user's choice. Construct 1 is queued for a 3-piece Gibson assembly, which includes a PCR (from an existing plasmid template), a pair of annealed oligos, and a restriction digest of an existing plasmid backbone. Construct 2 is queued as a 5-piece Golden Gate assembly which includes 3 existing precloned parts (liberated by type IIS digest of plasmids), an existing PCR product, and a gBlock (ordered as an oligonucleotide and used directly in Golden Gate), each of which is Registered as supplying a Golden Gate part with compatible junctions. The Golden Gate assembly is planned using the Golden Gate Assembly Queuer tool. Next, the constructs are assembled and transformed. Colonies could be miniprepped directly (construct 1) and multiple clones checked by e.g., Sanger sequencing, or colony stocks could be prepared and assessed by analytical PCR (construct 2), then a single clone verified by nanopore sequencing. Verified clones are transformed and stored as strain cryostocks. The Experiment becomes ready to do as soon as both constructs r1 and r2 are sequence-verified.

**CloneCoordinate Thuronyi Lab 2019-2025** archived copy

Recommended (efficient) tasks are highlighted in green and high-priority tasks in lavender + blue. More intense color means the task has not been done in a long time.

| # of priority tasks ready | # of tasks ready | type of sample | procedure | status text shown when ready | # queued | days since last done (25-351) | good # to do | time estimate | protocol quality | task # | type | skill set |
|---|---|---|---|---|---|---|---|---|---|---|---|---|
| 0 | 3 | oligos o | make backup and working stocks | ready to stock | 2 | 96 | anything ready | 1 h | good | 1 | bench | pipetting, tube strips |
| 0 / 0 / 0 | 0 / 0 / 0 | dsDNA d PCR / anneal / digest | set up PCR / annealing / digest | ready to set up | 6 | 52 | 8, 16, or 24 | 1.5 h / 16 | good | 2 | bench | pipetting, tube strips |
| 0 / 0 | 0 / 0 | dsDNA d PCR / digest | purify completed PCRs / digests | ready to clean up | | 46 | up to 24 | 45 min / 16 | fair | 3 | bench | DNA purification |
| 0 | 0 | dsDNA d | dilute & store annealed oligos | ready to dilute & store | | 139 | anything ready | 20 min / 16 | good | 4 | bench | pipetting, tube strips |
| 4 | 10 | assemblies a | assemble from d fragments | ready to assemble | 23 | 63 | 8, 16, or 24 | 1.5 h / 16 | good | 5 | bench | pipetting, tube strips |
| 0 | 0 | Golden Gate g | assemble from m donors | ready to assemble | 0 | 56 | 8 | 2 h / 16 | good | 6 | bench | pipetting, tube strips |
| 1 / 3 | 1 / 3 | assemblies a / GGate g | transform completed assemblies | ready to transform | | 27 | 8, 16, or 24 | 3 h / 16 | good | 7 | bench | comp cell & Petri dish use |
| 0 | 0 | transformations | score transformation plates | ready to score plates | | 26 | anything ready | 2 min each | good | 8 | bench | colony inspection |
| 0 / 0 | 0 / 0 | assemblies a / GGate g | queue preps of successful transformations/assemblies | ready to queue miniprep | | | anything ready | 10 min | good | 9 | bench | queueing in CC |
| 15 / 7 | 18 / 11 | minipreps m plates / strains | inoculate liquid culture | ready to inoculate for miniprep | 3 | 26 | 24 or 48 | 1h / 24 | good | 10 | bench | inoculation |
| 0 | 0 | minipreps m | prep plasmids after overnight growth (2 day growth max) | ready to miniprep | | 25 | 1-2 plates of 24 | 2 h / 1 plate; 3 h / 2 plates | excellent | 11 | bench | DNA purification |
| 0 / 20 | 5 / 21 | dsDNA d / minipreps m | quantitate using Take3 | ready to quant | | 62 | 16, 32, 64... | 30 min for 32 | good | 12 | bench | DNA quantitation |
| 0 / 20 | 3 / 21 | dsDNA d / minipreps m | gel sample prep | ready to prep gel sample | | 57 | 8, 16, 32... | 2 min each | good | 13 | bench | pipetting, tube strips |
| 0 / 0 | 0 / 0 | transformations / minipreps m | queue analytical PCR | ready to queue analytical PCR | | 351 | as needed | 2 min each | fair | 14 | remote | queueing in CC |
| 0 | 0 | analytical PCR | set up analytical PCR | ready to set up | 0 | 350 | 8, 16, 32... | 1 h for 16 | fair | 15 | bench | pipetting, tube strips |
| 4 / 0 | 11 / 0 | prepped samples / analytical PCRs | load prepped samples, run gel, image gel | ready to run gel | | 116 | >80 samples = full gel | 1.5 h | good | 16 | bench | gel use |
| 0 / 0 / 0 | 0 / 0 / 0 | d / m / analytical PCR | analyze previously run gel | ready to code gel | | 108 | anything ready | 15 min per row | poor | 17 | remote | gel and cloning concepts |
| 0 | 0 | minipreps m (# constructs) | queue sequencing | ready to queue sequencing | | 35 | anything ready | 5 min each | good | 18 | remote | queueing in CC |
| 0 / 0 | 0 / 0 | sequencing Sanger / whole-plasmid | mix sequencing samples | ready to mix | 3 / 1 | 35 | anything ready | 20 min / 8 | good | 19 | bench | pipetting, tube strips |
| 2 | 3 | sequencing | align sequencing reads to maps | ready to align | | 32 | anything ready | 2 min each | fair | 20 | remote | construct management software |
| 0 | 5 | sequencing | interpret sequencing | ready to interpret | | 32 | anything ready | variable | fair | 21 | remote | construct management software, cloning |
| 12 | 147 | minipreps m | transform for strain storage | ready to transform for strain storage | | 134 | 8 (16 at most) | 3 h / 8 to 16 | incomplete | 22 | bench | comp cell & Petri dish use |
| 0 | 3 | minipreps m | inoculate overnight for storage | ready to inoculate for strain storage | | 133 | 8, 16, or 24 | 30 min / 8 | fair | 23 | bench | inoculation |
| 0 | 0 | strains s | store overnight culture as new strain | ready to store | | 132 | 8, 16, or 24 | 30 min / 8 | good | 24 | bench | pipetting, tube strips |
| | 0 | experiments e | test and learn from constructs | ready to start | 4 | 293 | anything ready | variable | n/a | 25 | bench | scientific method |

0 oligos are ready to order, length ≤ 60
0 oligos are ready to order, length > 60
0 dsDNA or ultramers are ready to order
0 sequencing rxns are queued/mixed for Sanger
0 sequencing rxns are ready to submit for Sanger
0 sequencing rxns are queued/mixed for whole-plasmid sequencing
0 sequencing rxns are ready to submit for whole-plasmid sequencing

**CC News**
250610
(0 days ago)

- Version 1.0 of CC will shortly be available as a public release! See clonecoordinate.org for details.
- News posts for all CC instances will now be pulled from a public Sheet so they can be updated remotely

**Supplementary Figure 2. CC Dashboard.** The task list reflects the CloneCoordinate Thuronyi Lab 2019-2025 archived CC instance. The task count thresholds for highlighting and the contents of the “good # to do,” “time estimate,” and “protocol quality” columns can be configured in CC Settings. Each row in the “procedure” column can be hyperlinked to protocols, also specified in Settings.

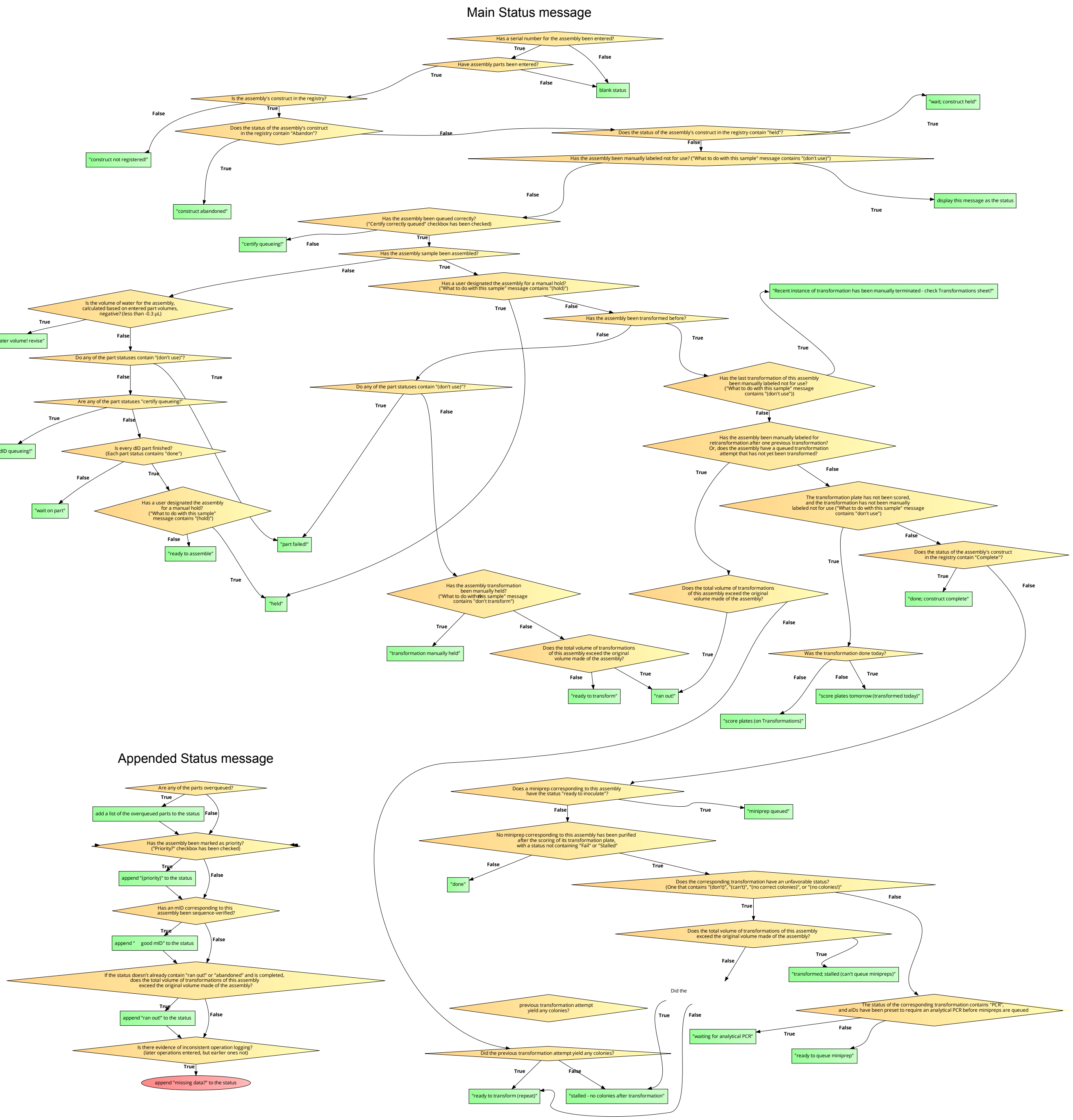

**Supplementary Figure 3. Flow chart for Assemblies Status code.** Zoom in to view details. The flow chart presents a simplified version of the Sheets formulas that set the Status field for Assemblies. The complete code can be viewed in CC itself. Interactive flow charts for each Status formula are linked from **Supplementary Table 1** and narrative descriptions are in **Supplementary Text 3**.

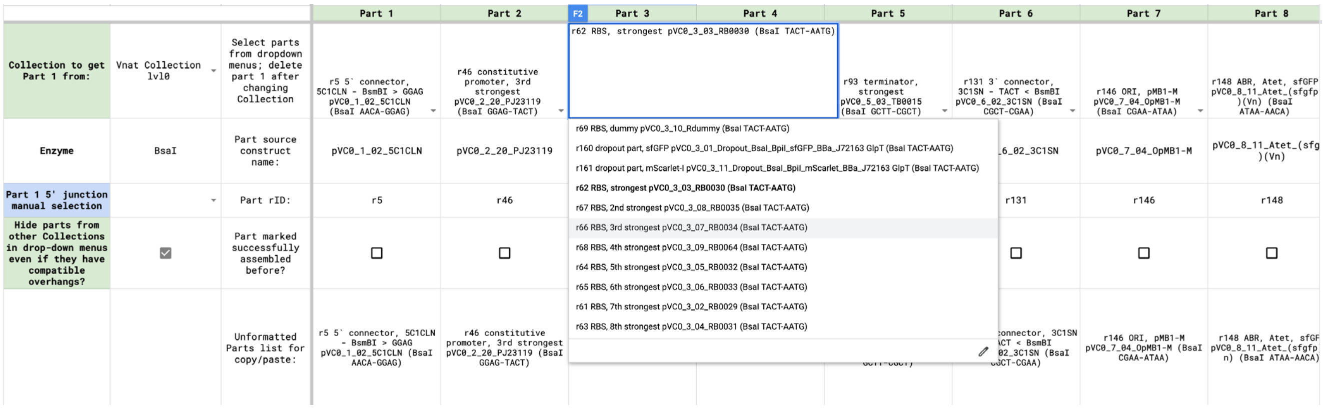

**Supplementary Figure 4. CCA Golden Gate Assembly Queuer tool screenshot.** An example assembly using parts from the Vnat Collection (Faber et al., Expanding genetic engineering capabilities in *Vibrio natriegens* with the Vnat Collection. Nucleic Acids Research (**2025**), manuscript in proof) is shown. Working from left to right, parts are chosen from dropdown menus that populate dynamically to display only those parts from the linked CC Registry that will assemble with the previous part, that is, they share the same type IIS enzyme and the 3' junction sequence of the previous part matches the 5' junction sequence of the next part (meaning that the overhang generated at the 3' end of the previous part will be reverse complementary to the overhang generated at the 5' end of the next part). Restriction enzyme, 5' and 3' junction sequences, part descriptive text, sort order, and Golden Gate collection are set in the CC Registry. When connected to a full CC instance (rather than a stand-alone Registry as shown here), the dropdown options will also indicate whether each part is currently in stock (ready to use) based on miniprep inventory. The parts list generated by the tool can be pasted into the Golden Gate g tab of CC to queue an assembly. Palindromic or repeated junctions within the assembly are flagged automatically in the Queuer tool and junction validity is checked when queueing the assembly in CC.

| Experiments | | | 1. Sample properties | | 2a. Queueing, general | | | | |
|---|---|---|---|---|---|---|---|---|---|
| Serial # | ID | Status | What to do with this experiment? | General notes | Date queued | Queued by | Experiment title | Short description / experimental question | Notebook entry / link |
| 1 | e1 | Complete ♥ | | | 240719 | EJJ | Promoter test | Which promoters have good maximum induction? | https://benchling.com/s/etr-drDfjghaWVObM4kdHcWTj?m=slm-k5sKvz0dsmkr387f |
| 2 | e2 | Complete ♥ | | | 240731 | BWT | Strain build | Create new strain | https://benchling.com/s/etr-drDfjghfdjksdgjkhEWdHcWTj?m=slm-kcs,lf38SsmR5uEcZ |
| 3 | e3 | ready to start | | | 240731 | BWT | Burden | Assess metabolic burden for new strain | https://benchling.com/s/etr-drsjdwelkObM4kdHcWTj?m=slm-k5sKvz0dfs394R5uEcZ |
| 4 | e4 | Required ID is marked ⛔! | | | 240802 | ERQ | Output | Measure pathway output for new strain | https://benchling.com/s/etr-drDfjghaWVObcmkfjkeWTj?m=slm-k5sKvfmskfjdk325 |
| 5 | e5 | Wait on constructs | | | 240920 | BWT | Homolog test | Replace pathway enzyme with homolog | https://benchling.com/s/etr-drDfjghafdkljEfgeWTj?m=slm-k5sKvfmskfjdk325 |

| | | | | | 2b. Queueing, constructs/IDs | | | | |
|---|---|---|---|---|---|---|---|---|---|
| Serial # | ID | Status | Project | Assay(s) needed / notes | Describe control condition(s) | Control construct(s) / IDs needed | Describe experimental condition(s) | Experimental construct(s) / IDs needed | Status needed to consider ID(s) to be ready to use |
| 1 | e1 | Complete ♥ | Project A | Measure log phase sfGFP expression in plate reader | empty plasmid (no sfGFP); constitutive sfGFP | pBWT001, pBWT002 | promoter variants; induce with 1 mM IPTG | pBWT003, pBWT004, pBWT005 | |
| 2 | e2 | Complete ♥ | Project B | standard transformation procedure | | | transform pBWT006 into s25 to create s26 | pBWT006, s25 | |
| 3 | e3 | ready to start | Project B | Measure growth curves, OD700 in plate reader | empty strain | s25 | measure pathway burden | pBWT006, s26 | done |
| 4 | e4 | Required ID is marked ⛔! | Project B | GCMS on 24h production culture, standard procedure | empty strain, current top producer strain | s25, s6 | test new pathway variants | s26, s27, s28 | done |
| 5 | e5 | Wait on constructs | Project C | clone constructs for testing later | | | | pBWT008, pBWT009, pBWT010 | |

| | | | | 3. Execution | | |
|---|---|---|---|---|---|---|
| Serial # | ID | Status | Statuses | Experiment ready? | Notification threshold | Email to notify when ready (or status changes) |
| 1 | e1 | Complete ♥ | pBWT001: Complete ♥<br>pBWT002: Complete ♥<br>pBWT003: Complete ♥<br>pBWT004: Complete ♥<br>pBWT005: Complete ♥ | Complete | turn off email notifications for now | |
| 2 | e2 | Complete ♥ | pBWT006: Complete ♥<br>s25: OK, done | Complete | notify when any Status changes (may generate lots of emails) | |
| 3 | e3 | ready to start | pBWT006: Complete ♥<br>s25: OK, done<br>s26: OK, done | Ready | notify when any Status changes (may generate lots of emails) | |
| 4 | e4 | Required ID is marked ⛔! | s26: OK, done<br>s27: Contaminated / heterogeneous! ⛔<br>s28: currently unused (empty) | Required ID is marked ⛔! | notify when experiment is ready or stalled (few emails) | |
| 5 | e5 | Wait on constructs | pBWT008: Queued<br>pBWT009: Queued<br>pBWT010: Complete ♥ | Wait on constructs | | |

**Supplementary Figure 5. CC Experiments tab example.** This tab tracks Experiments which can reference constructs and/or CC sample IDs (such as strains, shown here) as control and/or experimental conditions. It allows CC users to connect their experimental plans to CC's cloning progress so that work can be carried out as soon as the required materials are ready. Users can specify a particular Status that the referenced IDs must have before the Experiment is ready to do (here, strains must be "done," marked complete). Constructs must be completed (cloned) for an Experiment to be ready. If constructs are held or abandoned or any listed samples have problems (indicated by the ⛔ character), the Experiment status changes to reflect this (e.g., for e4, a required strain was found to be faulty). In addition, the Experiments tab is read into the CCA Construct Tracker Sheet, which maintains information about whether each Queued construct is proceeding through the build process or has become stalled, needing manual intervention to troubleshoot problems. This CCA Sheet has Conditional Notifications which, if enabled, will generate emails to the listed address(es) when constructs become stalled and/or IDs change status according to the selected choices, facilitating prompt attention to problems. The entries are fictitious and for illustrative purposes only.

**A**

**Assist: dsDNA**

Plates needed: m05 m06 m07 m37 o10 o12 o13 o15 o16
Premix A: Q5U-MM-1.5x 110.6 µL; Q5U 1.65 µL; 5x CES 31.4 µL; ET-SSB 1.65 µL
Premix B: Q5U-MM-1.5x 73.7 µL; Q5U 1.1 µL; 5x CES 22 µL
Premix C: Q5U-MM-1.5x 73.7 µL; Q5U 1.1 µL; water 22 µL

| Sort # | Strip pos'n | ID | Status | Type | Scale (µL) | Queue notes | Premix | | Additional component(s) | | | | | | | | | | |
|---|---|---|---|---|---|---|---|---|---|---|---|---|---|---|---|---|---|---|---|
| 1 | s1-1 | d1365 | ready to set up | PCR | 50 µL | | | | Q5U-MM-1.5x | 33.50 µL | ☐ | Q5U | 0.500 | ☐ | water | 7.50 | ☐ | DMSO 2.50 | ☐ |
| 2 | s1-2 | d1373 | ready to set up | PCR | 50 µL | | Premix A 43.5 µL | ☐ | | | | | | | | | | | |
| 3 | s1-3 | d1372 | ready to set up | PCR | 50 µL | | Premix A 43.5 µL | ☐ | | | | | | | | | | | |
| 4 | s1-4 | d1376 | ready to set up | PCR | 50 µL | | Premix B 43.5 µL | ☐ | | | | | | | | | | | |
| 5 | s1-5 | d1368 | ready to set up | PCR | 50 µL | | Premix B 43.5 µL | ☐ | | | | | | | | | | | |
| 6 | s1-6 | d1367 | ready to set up | PCR | 50 µL | | Premix C 43.5 µL | ☐ | | | | | | | | | | | |
| 7 | s1-7 | d1366 | ready to set up | PCR | 50 µL | | Premix C 43.5 µL | ☐ | | | | | | | | | | | |
| 8 | s1-8 | d1374 | ready to set up | PCR | 50 µL | | Premix A 43.5 µL | ☐ | | | | | | | | | | | |

**Assist: dsDNA**

Blocks needed: 3
63° anneal, 30 cycles (6)
63° anneal, 35 cycles (1)
67° anneal, 35 cycles (1)

**Elongation times (minutes:seconds):**
**63° anneal, 30 cycles - 0:21**
**63° anneal, 35 cycles - 0:32**
**67° anneal, 35 cycles - 0:33**

| Sort # | Strip pos'n | ID | Oligo 1 (well) | µL | | Oligo 2 (well) | µL | | Input DNA well | Input DNA volume (µL) | | # of cycles | Anneal temp (°C) | Required elong (s) | Notes / misc conditions |
|---|---|---|---|---|---|---|---|---|---|---|---|---|---|---|---|
| 1 | s1-1 | d1365 | o16.D03 | 3.00 | ☐ | o16.G03 | 3.00 | ☐ | m06.D12 | 0.500 | ☐ | 30 | 63 | | |
| 2 | s1-2 | d1373 | o16.D03 | 3.00 | ☐ | o16.E03 | 3.00 | ☐ | m07.D01 | 0.500 | ☐ | 30 | 63 | | |
| 3 | s1-3 | d1372 | o15.A08 | 3.00 | ☐ | o15.D04 | 3.00 | ☐ | m05.F07 | 0.500 | ☐ | 30 | 63 | | |
| 4 | s1-4 | d1376 | o10.E04 | 3.00 | ☐ | o12.F09 | 3.00 | ☐ | m37.B05 | 0.500 | ☐ | 35 | 63 | | |
| 5 | s1-5 | d1368 | o15.B11 | 3.00 | ☐ | o15.G01 | 3.00 | ☐ | m05.A10 | 0.500 | ☐ | 35 | 67 | | |
| 6 | s1-6 | d1367 | o16.F03 | 3.00 | ☐ | o16.G03 | 3.00 | ☐ | m06.D12 | 0.500 | ☐ | 30 | 63 | | |
| 7 | s1-7 | d1366 | o16.F03 | 3.00 | ☐ | o16.E03 | 3.00 | ☐ | m06.D12 | 0.500 | ☐ | 30 | 63 | | |
| 8 | s1-8 | d1374 | o13.H04 | 3.00 | ☐ | o13.A05 | 3.00 | ☐ | m07.E12 | 0.500 | ☐ | 30 | 63 | | |

**B**

**Assist: Golden Gate**

**Premix enabled: see instructions in rows below. Components in premix are grayed out in table.**
**m plates needed: 19 24 31 36 37 38**

| Sort # | Strip pos'n | ID | Premix component | µL in premix | | µL water | | µL Premix | | Well, Part 1 | µL 1 | | Well, Part 2 | µL 2 | | Well, Part 3 | µL 3 | | Well, Part 4 | µL 4 |
|---|---|---|---|---|---|---|---|---|---|---|---|---|---|---|---|---|---|---|---|---|
| 1 | s1-1 | g729 | *water* | *28.0* | ☑ | 0.146 | ☐ | 9.02 | ☐ | *m31.D08* | *0.226* | ☑ | ~~m37.C05~~ | ~~0.527~~ | | ~~m36.A09~~ | ~~0.308~~ | | ~~m24.G06~~ | ~~0.098~~ |
| 2 | s1-2 | g730 | *m37.C05* | *2.93* | ☑ | 0.203 | ☐ | 9.02 | ☐ | *m31.E08* | *0.226* | ☑ | ~~m37.C05~~ | ~~0.527~~ | | ~~m36.A09~~ | ~~0.308~~ | | ~~m24.G06~~ | ~~0.098~~ |
| 3 | s1-3 | g731 | *m36.A09* | *1.711* | ☑ | 0.220 | ☐ | 9.02 | ☐ | m36.A08 | 0.226 | ☐ | ~~m37.C05~~ | ~~0.527~~ | | ~~m36.A09~~ | ~~0.308~~ | | ~~m24.G06~~ | ~~0.098~~ |
| 4 | s1-4 | g732 | m24.G06 | 0.544 | ☐ | | | 9.02 | ☐ | m37.C06 | 0.226 | ☐ | ~~m37.C05~~ | ~~0.527~~ | | ~~m36.A09~~ | ~~0.308~~ | | ~~m24.G06~~ | ~~0.098~~ |
| 5 | s1-5 | g733 | m37.D05 | 2.74 | ☐ | | | 9.02 | ☐ | m36.A08 | 0.226 | ☐ | ~~m37.C05~~ | ~~0.527~~ | | ~~m36.A09~~ | ~~0.308~~ | | ~~m24.G06~~ | ~~0.098~~ |
| | | | m38.F01 | 2.89 | ☐ | | | | | | | | | | | | | | | |
| | | | m24.H08 | 3.00 | ☐ | | | | | | | | | | | | | | | |
| | | | NEB 10x T4 ligase buffer | 5.55 | ☐ | | | | | | | | | | | | | | | |
| | | | T4 ligase | 2.78 | ☐ | | | | | | | | | | | | | | | |

**Assist: Golden Gate**

| Sort # | Strip pos'n | ID | Well, Part 5 | µL 5 | | Well, Part 6 | µL 6 | | Well, Part 7 | µL 7 | | Well, Part 8 | µL 8 | | µL NEB 10x T4 ligase buffer | | µL T4 ligase | | Enzyme | µL RE | |
|---|---|---|---|---|---|---|---|---|---|---|---|---|---|---|---|---|---|---|---|---|---|
| 1 | s1-1 | g729 | ~~m37.D05~~ | ~~0.493~~ | | ~~m38.F01~~ | ~~0.520~~ | | ~~m24.H08~~ | ~~0.541~~ | | m37.E05 | 0.355 | ☐ | ~~1.000~~ | ☐ | ~~0.500~~ | ☐ | BsaI | 0.250 | ☐ |
| 2 | s1-2 | g730 | ~~m37.D05~~ | ~~0.493~~ | | ~~m38.F01~~ | ~~0.520~~ | | ~~m24.H08~~ | ~~0.541~~ | | m38.H04 | 0.298 | ☐ | ~~1.000~~ | ☐ | ~~0.500~~ | ☐ | BsmBI | 0.250 | ☐ |
| 3 | s1-3 | g731 | ~~m37.D05~~ | ~~0.493~~ | | ~~m38.F01~~ | ~~0.520~~ | | ~~m24.H08~~ | ~~0.541~~ | | m38.D01 | 0.281 | ☐ | ~~1.000~~ | ☐ | ~~0.500~~ | ☐ | BsaI | 0.250 | ☐ |
| 4 | s1-4 | g732 | ~~m37.D05~~ | ~~0.493~~ | | ~~m38.F01~~ | ~~0.520~~ | | ~~m24.H08~~ | ~~0.541~~ | | m37.D11 | 0.501 | ☐ | ~~1.000~~ | ☐ | ~~0.500~~ | ☐ | BsaI | 0.250 | ☐ |
| 5 | s1-5 | g733 | ~~m37.D05~~ | ~~0.493~~ | | ~~m38.F01~~ | ~~0.520~~ | | ~~m24.H08~~ | ~~0.541~~ | | m19.F06 | 0.405 | ☐ | ~~1.000~~ | ☐ | ~~0.500~~ | ☐ | BsaI | 0.250 | ☐ |

**Supplementary Figure 6. Assistant tabs examples.** The Assistant tabs are populated by user-selected samples from the corresponding tab in order of assigned Sort #. Where applicable, premixes are calculated for samples that are ready for a given task (**Supplementary Text 4**). Check boxes allow users to track which components have been added. **A, Asst: dsDNA showing PCRs. B, Asst: Golden Gate.**

**A**

| Statistics All time | Everyone | B / BWT | Arnav / ADS | Evelyn / ERQ | Katherine / KZY | Maliha / MCR | Santiago / STC | Simms / SB | Sofija / SO |
|---|---|---|---|---|---|---|---|---|---|
| Comprehensive achievement? | | | | 🥳🦍🎉 | | | | | |
| Oligos o queued | 1466 | 228 | 4 | 39 | 0 | 0 | 0 | 3 | 0 |
| Oligos o stocked | 1461 | 32 | 4 | 129 | 0 | 0 | 0 | 0 | 0 |
| dsDNA d queued | 1373 | 299 | 18 | 43 | 16 | 4 | 11 | 0 | 0 |
| dsDNA d PCRs run | 1061 | 24 | 23 | 75 | 11 | 0 | 55 | 4 | 0 |
| dsDNA d annealed | 292 | 10 | 0 | 28 | 11 | 6 | 1 | 0 | 1 |
| dsDNA d purified | 1041 | 70 | 28 | 121 | 25 | 13 | 53 | 4 | 5 |
| Assemblies a queued | 1045 | 265 | 0 | 43 | 30 | 3 | 14 | 0 | 0 |
| Assemblies a assembled | 957 | 15 | 11 | 102 | 7 | 23 | 21 | 0 | 3 |
| Golden Gate g assembled | 585 | 69 | 0 | 126 | 0 | 13 | 25 | 4 | 0 |
| Transformations | 1686 | 148 | 27 | 90 | 64 | 54 | 16 | 28 | 16 |
| Plates scored | 1634 | 288 | 50 | 56 | 33 | 22 | 22 | 12 | 6 |
| Minipreps m queued | 3728 | 955 | 48 | 214 | 16 | 107 | 24 | 0 | 0 |
| Minipreps m inoculated | 3635 | 417 | 72 | 277 | 24 | 69 | 24 | 24 | 0 |
| Minipreps m done | 3575 | 386 | 72 | 355 | 0 | 45 | 47 | 24 | 24 |
| Analytical PCRs run | 390 | 0 | 0 | 7 | 0 | 0 | 0 | 0 | 0 |
| Sequencing queued | 1616 | 596 | 0 | 87 | 0 | 3 | 31 | 2 | 0 |
| Sequencing prepared | 1600 | 166 | 14 | 101 | 6 | 6 | 32 | 5 | 0 |
| Sequencing aligned | 1536 | 698 | 0 | 45 | 23 | 22 | 0 | 0 | 0 |
| Sequencing interpreted | 1486 | 731 | 0 | 46 | 0 | 0 | 0 | 0 | 0 |
| Transformations (strain) | 200 | 58 | 17 | 16 | 0 | 31 | 0 | 0 | 0 |
| Inoculate strain | 195 | 53 | 25 | 13 | 0 | 49 | 0 | 0 | 0 |
| Strains s stored | 795 | 316 | 25 | 19 | 0 | 23 | 0 | 0 | 0 |
| dsDNA d / Miniprep m Quantified | 4326 | 185 | 70 | 363 | 18 | 27 | 97 | 12 | 0 |
| Gel samples prepped | 3636 | 206 | 92 | 226 | 28 | 14 | 37 | 0 | 3 |
| Gels run | 104 | 9 | 0 | 10 | 5 | 1 | 1 | 1 | 0 |
| Gel samples coded | 3980 | 584 | 0 | 442 | 61 | 0 | 83 | 0 | 6 |
| Constructs debugged | 235 | 174 | 0 | 14 | 24 | 4 | 15 | 0 | 0 |
| Constructs active | 778 | 91 | 0 | 50 | 0 | 0 | 1 | 4 | 0 |
| Constructs completed | 389 | 43 | 0 | 8 | 0 | 0 | 0 | 2 | 0 |
| % Constructs complete | 50% | 47% | | 16% | | | 0% | 50% | |
| Queued constructs in portfolio | 390 | 51 | 0 | 49 | 0 | 0 | 1 | 2 | 0 |
| % portfolio complete | | 77% | | 17% | | | 0% | 50% | |

**B**

| Statistics By date range | Show operations that occurred on or after date: | | 240900 | Show operations that occurred on or before date: | | 250101 | |
|---|---|---|---|---|---|---|---|
| | Everyone | B / BWT | Arnav / ADS | Evelyn / ERQ | Katherine / KZY | Maliha / MCR | Santiago / STC |
| Oligos o queued | 0 | 0 | 0 | 0 | 0 | 0 | 0 |
| Oligos o stocked | 33 | 0 | 0 | 0 | 0 | 0 | 0 |
| dsDNA d PCRs run | 39 | 0 | 3 | 0 | 0 | 0 | 31 |
| dsDNA d annealed | 12 | 0 | 0 | 0 | 5 | 6 | 0 |
| dsDNA d purified | 39 | 4 | 8 | 0 | 0 | 0 | 31 |
| Assemblies a assembled | 45 | 0 | 10 | 0 | 0 | 0 | 17 |
| Golden Gate g assembled | 4 | 0 | 0 | 0 | 0 | 0 | 4 |
| Transformations | 76 | 0 | 27 | 1 | 32 | 24 | 0 |
| Plates scored | 66 | 8 | 50 | 0 | 0 | 0 | 8 |
| Minipreps m inoculated | 96 | 24 | 72 | 0 | 24 | 24 | 0 |
| Minipreps m done | 112 | 40 | 72 | 0 | 0 | 0 | 0 |

**C**

| | Leader, 240900 - 250101 |
|---|---|
| Oligos o queued | |
| Oligos o stocked | Emily / EVG with 33 |
| dsDNA d PCRs run | Santiago / STC with 31 |
| dsDNA d annealed | Maliha / MCR with 6 |
| dsDNA d purified | Santiago / STC with 31 |
| Assemblies a assembled | Santiago / STC with 17 |
| Golden Gate g assembled | Santiago / STC with 4 |
| Transformations | Katherine / KZY with 32 |
| Plates scored | Arnav / ADS with 50 |
| Minipreps m inoculated | Arnav / ADS with 72 |
| Minipreps m done | Arnav / ADS with 72 |
| Analytical PCRs run | |
| Sequencing queued | Santiago / STC with 15 |

**Supplementary Figure 7. CCA Statistics output. A, All time task counts for active group members.** This view shows the number of tasks (operations on individual samples) completed by group members who are currently marked active. Jointly completed tasks are credited to all contributors. The highest total for each task is in bold with light gray background. Members such as Evelyn Qi who have completed every task at least once have their contributions shown in lavender and earn celebratory emojis in the top row (“Comprehensive achievement?”). **B, Date range filtered view (abridged)**. Viewers can adjust the start and/or end dates for which to total tasks. **C, Date range leaderboard (abridged)**. This view shows the top contributors and their totals within the selected period. Tasks with no entry were not carried out.

**A**

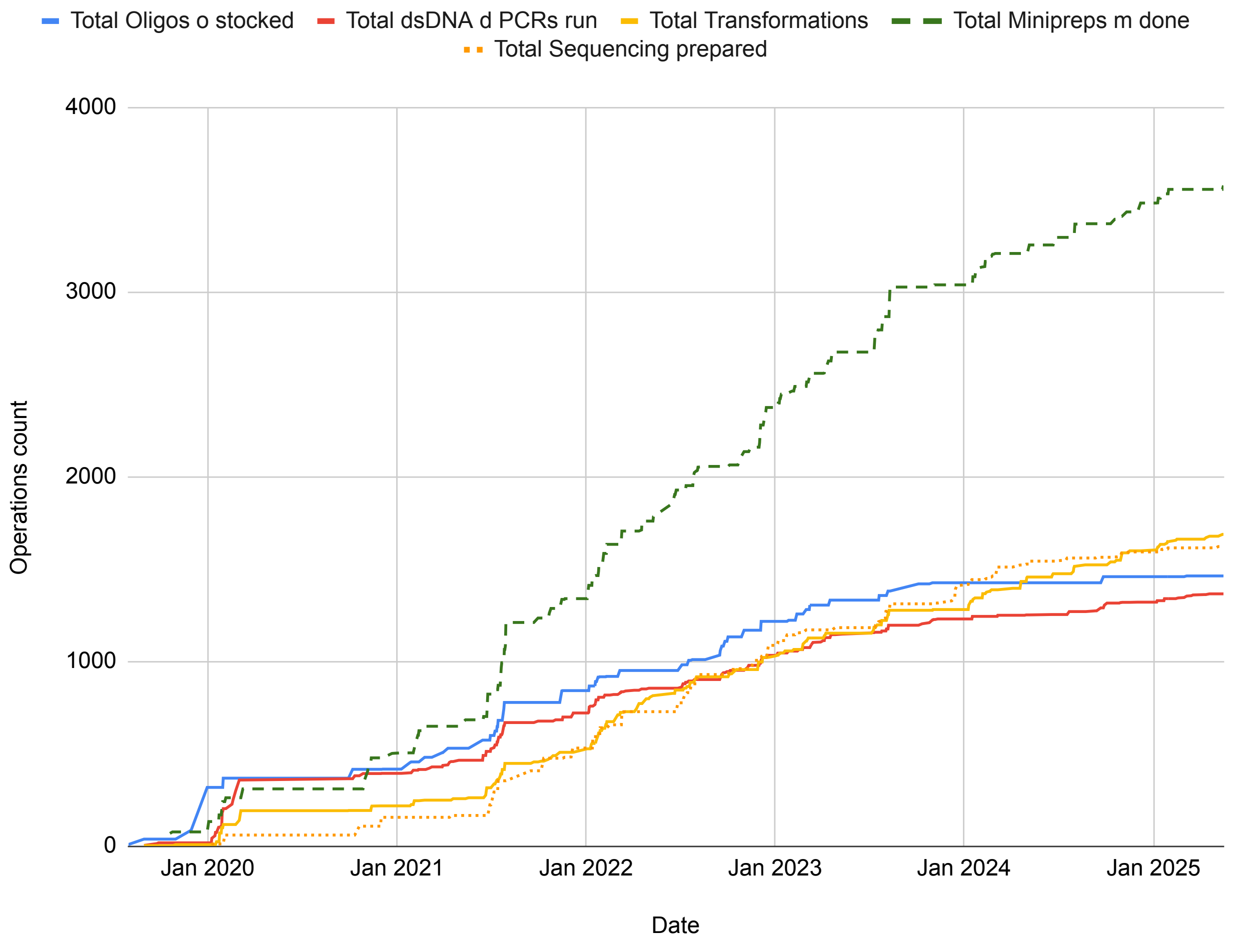

**B**

| Calculate and display this data set on the chart tab? | Operation to tabulate | Cumulative total so far | Average monthly total | Average annual total | Average days per operation |
|---|---|---|---|---|---|
| ☑ | Oligos o stocked | 1461 | 21 | 250 | 1 |
| ☑ | dsDNA d PCRs run | 1364 | 19 | 234 | 1.6 |
| ☑ | Transformations | 1686 | 24 | 289 | 1.3 |
| ☑ | Minipreps m done | 3575 | 51 | 613 | 0.6 |
| ☑ | Sequencing prepared | 1620 | 23 | 278 | 1.3 |

**Supplementary Figure 8. CCA Operations timeline output.** Up to 5 tasks can be shown at once on this CCA Sheet, including any of those listed on the CC Dashboard as well as construct registration, debugging, and completion. **A, Operations timeline plot.** The cumulative count of selected tasks over time is shown. **B, Operations table.** Totals and monthly, annual, and days per operation (in calendar time, not business days) statistics are tabulated for the selected tasks.

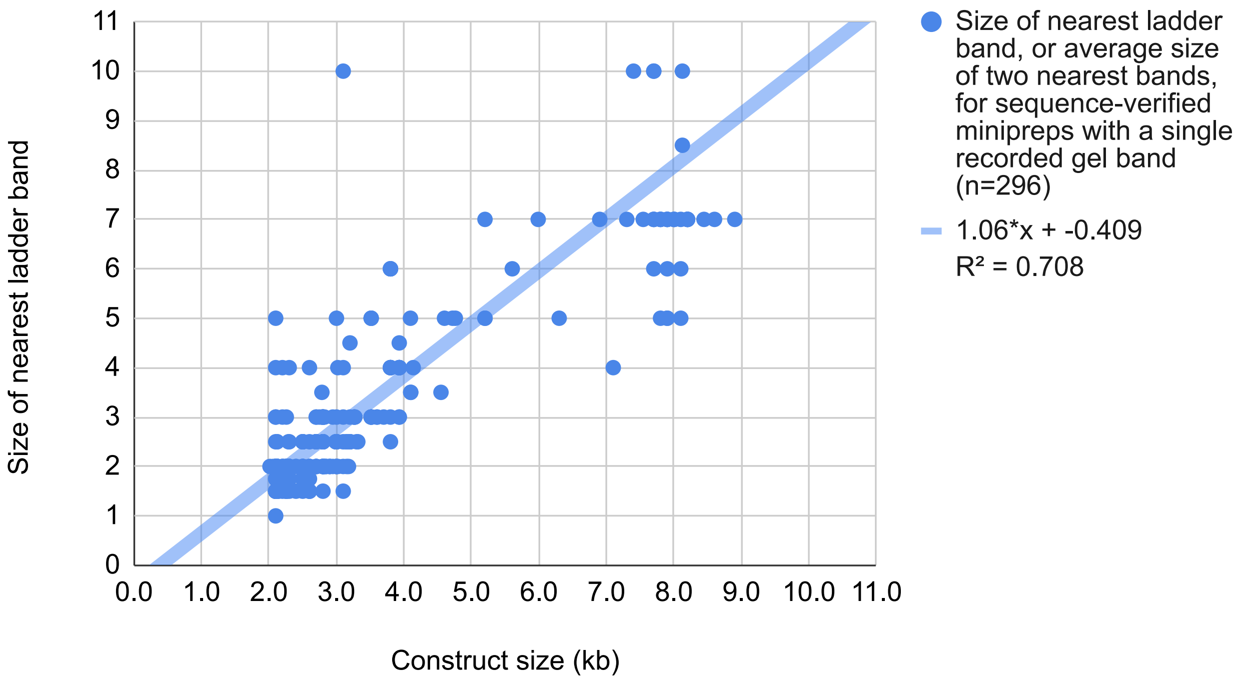

**Supplementary Figure 9. CCA Plasmid gel sizing output.** The plot shows the size of the nearest band or bands in the sizing ladder selected for agarose gel electrophoresis of sequence-verified minipreps vs. their known size in kilobase pairs. The ladder band size options are quantized. When 2 bands are selected in CC, their average size is used. The fit equation and the square of the Pearson correlation coefficient are shown.

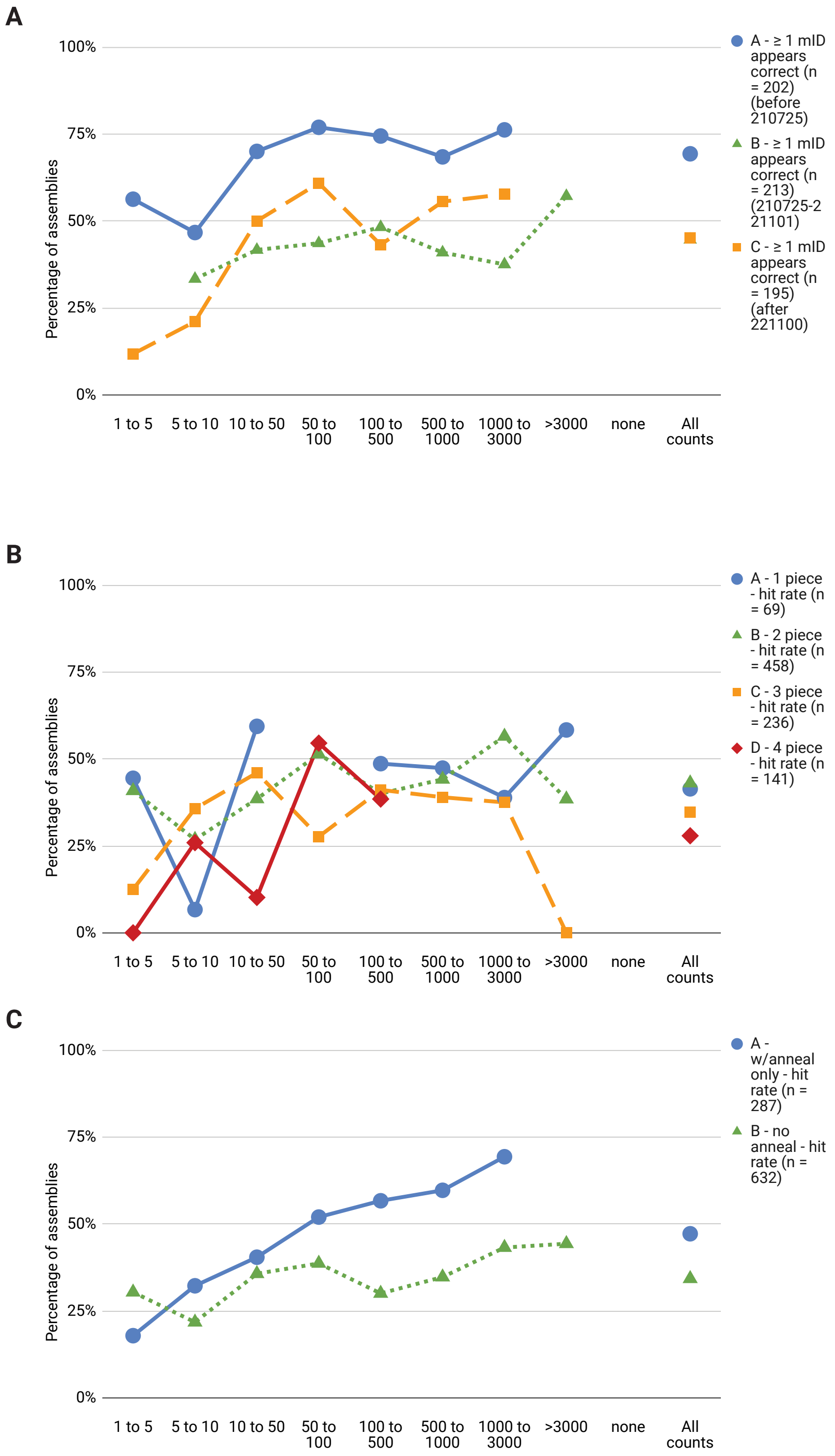

**Supplementary Figure 10. Additional analyses from CCA Miniprep outcome vs colony count**. The CCA Sheet reports data for USER assemblies (99%) and Gibson assemblies (1%) that resulted in at least one miniprepped colony. **A, Success rate by assembly transformation date**. The plot shows the percentage of assemblies for which at least one clone appears to be correct, based on sequencing and agarose gel electrophoresis data, as a function of colony count (left) or across all assemblies (far right). Percentages based on ≥10 assemblies are shown. The three series are filtered by transformation date so as to divide assemblies approximately into thirds (transformation before July 25, 2021, blue circles; from July 25, 2021 through November 1, 2022, green triangles; after November 1, 2022, orange squares). The overall success rates are 69%, 45% and 45%. **B, Hit rate by number of parts in assembly.** The plot shows the percentage of assessed clones that appear to be correct for each assembly (the empirical hit rate). Percentages based on ≥5 assemblies are shown. Each series shows assemblies with 1, 2, 3, or 4 separate DNA parts. The overall hit rates are 41%, 43%, 35%, and 28% respectively. **C, Hit rate by presence of annealed oligonucleotide part(s).** The plot shows the empirical hit rate for assemblies that contain (blue circles) or do not contain (green triangles) dsDNA fragments with 3’ cohesive end overhangs (generated by annealing partially reverse-complementary oligonucleotides) in addition to PCR products (which are prepared using dU-containing primers, then digested by the USER enzyme mixture to create 3’ cohesive end overhangs). The overall hit rates are 47% for assemblies that contain annealed oligonucleotides and 34% for assemblies that do not.

| | | Colony counts following transformation of assembly | | | | | | | | | |
|---|---|---|---|---|---|---|---|---|---|---|---|
| | | 1 to 5 | 5 to 10 | 10 to 50 | 50 to 100 | 100 to 500 | 500 to 1000 | 1000 to 3000 | >3000 | none | **All counts** |
| **Assembly count by proportion of mIDs appearing correctly assembled based on gel coding** | No mIDs appear correct | 20 | 27 | 45 | 34 | 64 | 35 | 28 | 11 | | **264** |
| | Ambiguous or unclear if any mIDs appear correct | 3 | 4 | 2 | 3 | 5 | 2 | 3 | 1 | | **23** |
| | At least one mID appears correct | 5 | 4 | 28 | 20 | 41 | 26 | 18 | 10 | | **152** |
| | All mIDs appear correct | 8 | 11 | 29 | 31 | 40 | 24 | 22 | 6 | | **171** |
| | Total | **36** | **46** | **104** | **88** | **150** | **87** | **71** | **28** | | **610** |
| | % of assemblies, all mIDs appear correct | 22% | 24% | 28% | 35% | 27% | 28% | 31% | 21% | | **28%** |
| | % of assemblies, ≥ 1 mID appears correct | 36% | 33% | 55% | 58% | 54% | 57% | 56% | 57% | | **53%** |
| | % of minipreps appearing correct (hit rate) | 26% | 27% | 38% | 44% | 41% | 44% | 51% | 44% | | **40%** |
| | # minipreps for hit rate calc | 50 | 87 | 158 | 129 | 246 | 128 | 91 | 30 | | 919 |

**Supplementary Figure 11. Data table for main text Figure 4b.**

**Show assemblies using competent cell batch:** 190828-NEB5α

| | | Colony counts following transformation of assembly: 1 to 5 | 5 to 10 | 10 to 50 | 50 to 100 | 100 to 500 | 500 to 1000 | 1000 to 3000 | >3000 | none | All counts |
|---|---|---|---|---|---|---|---|---|---|---|---|
| **Assembly count by proportion of mIDs appearing correctly assembled based on gel coding** | No mIDs appear correct | 6 | 3 | 11 | 3 | 11 | 4 | 3 | 1 | | **42** |
| | Ambiguous or unclear if any mIDs appear correct | 0 | 2 | 1 | 0 | 0 | 2 | 1 | 0 | | **6** |
| | At least one mID appears correct | 3 | 3 | 11 | 11 | 17 | 21 | 10 | 2 | | **78** |
| | All mIDs appear correct | 7 | 7 | 17 | 13 | 15 | 11 | 8 | 1 | | **79** |
| | Total | **16** | **15** | **40** | **27** | **43** | **38** | **22** | **4** | | **205** |
| | % of assemblies, all mIDs appear correct | 44% | 47% | 43% | 48% | 35% | 29% | 36% | | | **39%** |
| | % of assemblies, ≥ 1 mID appears correct | 63% | 67% | 70% | 89% | 74% | 84% | 82% | | | **77%** |
| | % of minipreps appearing correct (hit rate) | 48% | 66% | 46% | 72% | 53% | 68% | 68% | | | **55%** |
| | # minipreps for hit rate calc | 13 | 10 | 40 | 12 | 51 | 11 | 10 | | | 147 |

**Show assemblies using competent cell batch:** 210720-NEB5α

| | | Colony counts following transformation of assembly: 1 to 5 | 5 to 10 | 10 to 50 | 50 to 100 | 100 to 500 | 500 to 1000 | 1000 to 3000 | >3000 | none | All counts |
|---|---|---|---|---|---|---|---|---|---|---|---|
| **Assembly count by proportion of mIDs appearing correctly assembled based on gel coding** | No mIDs appear correct | 5 | 8 | 16 | 20 | 32 | 14 | 16 | 7 | | **118** |
| | Ambiguous or unclear if any mIDs appear correct | 0 | 1 | 1 | 2 | 3 | 0 | 2 | 1 | | **10** |
| | At least one mID appears correct | 2 | 2 | 15 | 10 | 21 | 10 | 6 | 9 | | **75** |
| | All mIDs appear correct | 0 | 3 | 7 | 10 | 12 | 3 | 8 | 3 | | **46** |
| | Total | **7** | **14** | **39** | **42** | **68** | **27** | **32** | **20** | | **249** |
| | % of assemblies, all mIDs appear correct | 0% | 21% | 18% | 24% | 18% | 11% | 25% | 15% | | **18%** |
| | % of assemblies, ≥ 1 mID appears correct | 29% | 36% | 56% | 48% | 49% | 48% | 44% | 60% | | **49%** |
| | % of minipreps appearing correct (hit rate) | 21% | 42% | 42% | 41% | 37% | 35% | 39% | 43% | | **39%** |
| | # minipreps for hit rate calc | 9 | 24 | 58 | 67 | 113 | 51 | 52 | 20 | | 394 |

**Show assemblies using competent cell batch:** 230223-Mach1

| | | Colony counts following transformation of assembly: 1 to 5 | 5 to 10 | 10 to 50 | 50 to 100 | 100 to 500 | 500 to 1000 | 1000 to 3000 | >3000 | none | All counts |
|---|---|---|---|---|---|---|---|---|---|---|---|
| **Assembly count by proportion of mIDs appearing correctly assembled based on gel coding** | No mIDs appear correct | 6 | 11 | 12 | 4 | 9 | 7 | 5 | 1 | | **55** |
| | Ambiguous or unclear if any mIDs appear correct | 3 | 1 | 0 | 1 | 2 | 0 | 0 | 0 | | **7** |
| | At least one mID appears correct | 2 | 2 | 6 | 2 | 12 | 5 | 6 | 0 | | **35** |
| | All mIDs appear correct | 1 | 1 | 5 | 6 | 12 | 10 | 6 | 2 | | **43** |
| | Total | **12** | **15** | **23** | **13** | **35** | **22** | **17** | **3** | | **140** |
| | % of assemblies, all mIDs appear correct | 8% | 7% | 22% | 46% | 34% | 45% | 35% | | | **31%** |
| | % of assemblies, ≥ 1 mID appears correct | 25% | 20% | 48% | 62% | 69% | 68% | 71% | | | **56%** |
| | % of minipreps appearing correct (hit rate) | 34% | 13% | 40% | 48% | 58% | 63% | 62% | | | **43%** |
| | # minipreps for hit rate calc | 17 | 35 | 36 | 15 | 33 | 23 | 16 | | | 175 |

**Show assemblies using competent cell batch:** commercial NEB5α C2987U

| | | Colony counts following transformation of assembly: 1 to 5 | 5 to 10 | 10 to 50 | 50 to 100 | 100 to 500 | 500 to 1000 | 1000 to 3000 | >3000 | none | All counts |
|---|---|---|---|---|---|---|---|---|---|---|---|
| **Assembly count by proportion of mIDs appearing correctly assembled based on gel coding** | No mIDs appear correct | 1 | 1 | 1 | 2 | 1 | 0 | 0 | 0 | | **6** |
| | Ambiguous or unclear if any mIDs appear correct | 0 | 0 | 0 | 0 | 0 | 0 | 0 | 0 | | **0** |
| | At least one mID appears correct | 0 | 1 | 0 | 2 | 2 | 0 | 0 | 0 | | **5** |
| | All mIDs appear correct | 0 | 0 | 0 | 2 | 1 | 0 | 0 | 0 | | **3** |
| | Total | **1** | **2** | **1** | **6** | **4** | **0** | **0** | **0** | | **14** |
| | % of assemblies, all mIDs appear correct | | | | 33% | | | | | | **21%** |
| | % of assemblies, ≥ 1 mID appears correct | | | | 67% | | | | | | **57%** |
| | % of minipreps appearing correct (hit rate) | | | | 50% | | | | | | **50%** |
| | # minipreps for hit rate calc | | | | 8 | | | | | | 8 |

**Supplementary Figure 12. Data tables for main text Figure 4c.**

### Supplementary Data

The Thuronyi Lab CloneCoordinate instance (version 0.99) containing cloning data from 2019-2025 can be viewed at the following link. Sequence information and construct descriptions have been removed to maintain privacy for ongoing unpublished research.
https://docs.google.com/spreadsheets/d/1wZwXvLzPqq4h6Fzb-27fKORN6zZ60yFWyrfPe6eCVts

### Supplementary Table 1: Status Code Flow Charts

| Tab | Status code flow chart link (interactive) | PDF link |
|---|---|---|
| Registry | https://app.code2flow.com/86X12HmG9w3E | CC Registry Status 1.0.pdf |
| Oligos o | https://app.code2flow.com/4CSdIpQljbjG | CC Oligos Status 1.0.pdf |
| dsDNA d | https://app.code2flow.com/Q8S9sS540ndl | CC dsDNA Status 1.0.pdf |
| Assemblies a | https://app.code2flow.com/Q3EgDMcVU07m | CC Assemblies Status 1.0.pdf |
| Golden Gate g | https://app.code2flow.com/sFQrMyTGuZ6G | CC Golden Gate Status 1.0.pdf |
| Transformations t | https://app.code2flow.com/b3npv35CZszM | CC Transformations Status 1.0.pdf |
| Minipreps m | https://app.code2flow.com/TmXxtrjmmZFR | CC Minipreps Status 1.0.pdf |
| Analytical PCR | https://app.code2flow.com/MeHbrNDmh5IA | CC Analytical PCR Status 1.0.pdf |
| Sequencing | https://app.code2flow.com/IT8JSkGMTBcH | CC Sequencing Status 1.0.pdf |
| Strains s | https://app.code2flow.com/Y9cFAxMXfGaA | CC Strains Status 1.0.pdf |

### Supplementary Text 1: CloneCoordinate Guide and User Manual

A user manual for CloneCoordinate is available at the following link:
https://docs.google.com/document/d/1Rky-_AYQLumPcQ87oZRaAi_a7sBCuaPWeaZY4-rezh8/

### Supplementary Text 2: CloneCoordinate Settings Documentation for v1.0

Settings documentation can be found at the following link, and is reproduced below.
https://docs.google.com/spreadsheets/d/1ifN-zsyOPhla7OHiAEViADX1Y4AJ4UqNp-3klbNFTJM/

| Tab(s) affected | Setting name | Description | Named Range designation in CC |
|---|---|---|---|
| **Settings highlighted in purple at the top are the ones you will almost certainly want to adjust when moving into CC for the first time. You can try the default values for the others initially and then change them as needed.** | | | |
| Dashboard | Lab name | What is your lab / group's name? (appears on Dashboard) | Settings_DashboardLabName |
| Dashboard | Protocol links list | Links to lab-specific protocols for each cloning task | Settings_DashboardProtocolLinksList |
| Registry | Project list | List of projects you are working on | Settings_RegistryProjectList |
| Assemblies a | Assembly recipes conditions | Components and volumes needed for different assembly recipes that you use | Settings_aAssemblyRecipesConditions |
| dsDNA d | Cleanup methods | Ways to purify PCR products you use | Settings_dCleanupMethods |
| dsDNA d | Default cleanup method | Default cleanup method | Settings_dDefaultCleanupMethod |
| dsDNA d | PCR polymerase components table | Components and volumes needed for each PCR DNA polymerase | Settings_dPCRPolymeraseComponentsTable |
| dsDNA d | PCR polymerase list | List of DNA polymerases you use for PCRs | Settings_dPCRPolymeraseList |
| dsDNA d | PCR variation components table | Components and volumes needed for each type of PCR additive | Settings_dPCRVariationComponentsTable |
| dsDNA d | PCR variation list | List of variations (additives) you use for PCRs | Settings_dPCRVariationList |
| dsDNA d | Thermocyclers incubators list | List of thermocyclers and/or incubators you use for PCRs and/or DNA assemblies | Settings_dThermocyclersIncubatorsList |
| Minipreps m | Purification methods | List of methods you use to purify plasmid DNA from bacterial cultures (miniprep) | Settings_mPurificationMethods |
| Transformations t | Competent cell batches | List of competent cell batches to show in Transformations dropdown menu - add new batches here | Settings_TransformationsCompetentCellBatches |
| Transformations t | Default comp cell batch | Default comp cell batch (remember to update this when a new batch is made!) | Settings_TransformationsDefaultCompCellBatch |
| All | Storage plate size | How do you want your physical samples to be stored (other than strains)? This setting affects where people are directed to find samples in the o, d, a, g, and m tabs and several Assistant tabs.<br><br>Enter your choice using the following format:<br>"#w" = # well plate<br>"b#" = box with # spaces<br>"#r#c" = plate with # rows, # cols<br><br>If you're using custom-sized boxes or plates (b# or #r#c) that don't appear in the dropdown menu, you'll need to type in your desired plate/box size using the specified format and ignore the data validation error. | Settings_SampleStoragePlateSize |
| Analytical PCR | PCR aID require queueing after transformation | For aID transformations: If analytical PCR is called for based on the priority levels set below (aID status shows "ready to queue analytical PCR"), require at least 1 analytical PCR to be queued and completed before the aID is marked "ready to queue miniprep"? | Settings_AnalyticalPCRaIDRequireQueueingAfterTransformation |
| Analytical PCR | PCR gID require queueing after transformation | For gID transformations: If analytical PCR is called for based on the priority levels set below (gID status shows "ready to queue analytical PCR"), require at least 1 analytical PCR to be queued and completed before the gID is marked "ready to queue miniprep"? | Settings_AnalyticalPCRgIDRequireQueueingAfterTransformation |
| Analytical PCR | PCR no abandoned constructs | If a construct is marked abandoned in the Registry, prevent analytical PCRs related to that construct from being marked "ready" for any operation? | Settings_AnalyticalPCRNoAbandonedConstructs |
| Assemblies a | Colony analytical PCR threshold | aID colony analytical PCR threshold | Settings_aIDColonyAnalyticalPCRThreshold |
| Assemblies a | Default scale | Assembly default scale | Settings_aDefaultScale |
| Assemblies a | Gibson exact match bases | Gibson number of exactly matching bases required to validate fragment overlap | Settings_aGibsonExactMatchBases |
| Assemblies a | Gibson premix fraction | Gibson Assembly premix is this fraction of the total assembly volume | Settings_aGibsonPremixFraction |
| Assemblies a | Gibson search distance | Gibson number of bases at ends of fragments to consider when searching for overlaps | Settings_aGibsonSearchDistance |
| Assemblies a | Ligation premix fraction | Ligation Assembly premix is this fraction of the total assembly volume | Settings_aLigationPremixFraction |
| Assemblies a | Miniprep analytical PCR threshold | aID-derived miniprep analytical PCR threshold | Settings_aIDMiniprepAnalyticalPCRThreshold |
| Assemblies a | Miniprep sequencing require analytical PCR? | For mIDs derived from aIDs: If analytical PCR is called for based on the priority levels set below (mID status shows "ready to queue analytical PCR"), require at least 1 analytical PCR to be queued and completed before the mID is marked "ready to queue sequencing"? | Settings_aIDMiniprepSequencingRequireAnalyticalPCR? |
| Assemblies a | Overage floor | Overage floor - make at least this much extra premix (µL) | Settings_aOverageFloor |

| Tab(s) affected | Setting name | Description | Named Range designation in CC |
|---|---|---|---|
| Assemblies a | Overage multiplier | Overage multiplier - call for this fraction of the amount of premix required (unitless) | Settings_aOverageMultiplier |
| Assemblies a | Premix bias | Premix bias: call for a premix unless NOT premixing saves at least this many pipetting steps. If bias is negative, premix will not be called for even when it saves some steps. | Settings_aPremixBias |
| Assemblies a | Store in plates? | Store completed assemblies in storage plates (checked) or temporarily as labeled individual samples (unchecked)? | Settings_aStoreInPlates? |
| Assemblies a | Target nku gibson | Gibson Target molar concentration of each DNA fragment (1 ng/kb/μL for dsDNA = 1.5 nM) | Settings_aTargetNkuGibson |
| Assemblies a | Target nku ligation | Ligation Target molar concentration of each DNA fragment (1 ng/kb/μL for dsDNA = 1.5 nM) | Settings_aTargetNkμLigation |
| Assemblies a | Target nku USER | USER Target molar concentration of each DNA fragment (1 ng/kb/μL for dsDNA = 1.5 nM) | Settings_aTargetNkuUSER |
| Assemblies a | USER premix fraction | USER Assembly premix is this fraction of the total assembly volume | Settings_aUSERPremixFraction |
| Assemblies a, Golden Gate g | Volume rounding | How do you want to round calculated assembly part volumes, when possible? This affects Assemblies a and Golden Gate g. Not yet implemented for CC v1.0. | Settings_aVolumeRounding |
| Dashboard | Good number to do list | How many of a task should someone do at once for efficiency (and manageability) | Settings_DashboardGoodNumberToDoList |
| Dashboard | Highlighting thresholds list | How many samples ready needed for tasks to be highlighted on the Dashboard as efficient choices for work | Settings_DashboardHighlightingThresholdsList |
| Dashboard | Lab leader name | Lab leader: always displays at the left of stats boards. You can leave this blank if you don't want to set a lab leader. | Settings_DashboardLabLeaderName |
| Dashboard | Lab member names | Lab member names (used by CCA Statistics) | Settings_DashboardLabMemberNames |
| Dashboard | Member active? | Is this group member currently working? Used by CCA Statistics. | Settings_DashboardMemberActive? |
| Dashboard | Protocol quality list | Estimate of how effectively the protocol can lead someone through the task - how much hands-on training should they have first? | Settings_DashboardProtocolQualityList |
| Dashboard | Remove departed? | Eventually remove departed members from roster? | Settings_DashboardRemoveDeparted? |
| Dashboard | Time estimate list | How long might it take to do the recommended number of samples for a task? | Settings_DashboardTimeEstimateList |
| dsDNA d | Agarose gel ladder band sizes | DNA ladder (agarose gels) band sizes in kb; use _1_ to indicate a more intense band at 1 kb | Settings_dAgaroseGelLadderBandSizes |
| dsDNA d | Agarose gel ladder choices | List of DNA ladders you use, with corresponding entries in ladder band sizes table | Settings_dAgaroseGelLadderChoices |
| dsDNA d | Anneal buffer volume | Buffer per reaction (μL) | Settings_dAnnealBufferVolume |
| dsDNA d | Anneal default scale | Standard scale, total reaction volume (μL) | Settings_dAnnealDefaultScale |
| dsDNA d | Anneal gel quant? | Allow gel sample prep and quantitation for annealed oligo samples? | Settings_QCAnnealGelQuant? |
| dsDNA d | Anneal min len | Minimum number of bases required for two oligos to be used as annealing inputs (error below this threshold) | Settings_dAnnealMinLen |
| dsDNA d | Anneal oligo volume | Oligo volume, each oligo (μL) | Settings_dAnnealOligoVolume |
| dsDNA d | Anneal warn len | Number of bases below which to show a warning for annealed oligos | Settings_dAnnealWarnLen |
| dsDNA d | Anneal water volume | Water per reaction (μL) | Settings_dAnnealWaterVolume |
| dsDNA d | Default elution volume | Default elution volume after purification (μL) | Settings_dDefaultElutionVolume |
| dsDNA d | dID gel sample threshold | dID gel sample threshold | Settings_QCdIDGelSampleThreshold |
| dsDNA d | dID quant threshold | dID quant threshold | Settings_QCdIDQuantThreshold |
| dsDNA d | Digest buffer list | Buffer used by each restriction enzyme you work with | Settings_dDigestBufferList |
| dsDNA d | Digest buffer volumes list | Volume of each buffer needed per standard scale reaction | Settings_dDigestBufferVolumesList |
| dsDNA d | Digest buffers table | Buffer used by each restriction enzyme you work with | Settings_dDigestBuffersTable |
| dsDNA d | Digest enzyme list | List of restriction enzymes you use for digests | Settings_dDigestEnzymeList |
| dsDNA d | Gel dID priority | Designate gel sample prep as a Dashboard task for dIDs when the sample "gel priority" column is set to | Settings_QCGeldIDPriority |
| dsDNA d | Low digest conc cutoff | Low concentration cutoff for digests - automatically mark PCRs below this value as failed and unusable | Settings_QCLowDigestConcCutoff |
| dsDNA d | Low digest conc warn | Warning concentration cutoff for digests - flag digests with concentrations below this value as potential failures | Settings_QCLowDigestConcWarn |
| dsDNA d | Low PCR conc cutoff | Low concentration cutoff for PCRs - automatically mark PCRs below this value as failed and unusable | Settings_QCLowPCRConcCutoff |

| Tab(s) affected | Setting name | Description | Named Range designation in CC |
|---|---|---|---|
| dsDNA d | Low PCR conc warn | Warning concentration cutoff for PCRs - flag PCRs with concentrations below this value as potential failures | Settings_QCLowPCRConcWarn |
| dsDNA d | Max samples per block | Maximum number of samples per PCR block | Settings_dMaxSamplesPerBlock |
| dsDNA d | Min volume to still use | Minimum estimated volume remaining of dID for sheet to validate it as available (µL) | Settings_dMinVolumeToStillUse |
| dsDNA d | Min volume warn | Volume remaining of dID for sheet to show a warning (µL) | Settings_dMinVolumeWarn |
| dsDNA d | PCR annealed oligo nku | default nkµ for annealed oligos (multiply by 1.5 for nmolar) | Settings_dPCRAnnealedOligoNku |
| dsDNA d | PCR assumed conc | Assumed concentration of an unquantitated PCR product (ng/µL) | Settings_dPCRAssumedConc |
| dsDNA d | PCR default anneal temp | Default anneal temperature | Settings_dPCRDefaultAnnealTemp |
| dsDNA d | PCR default cycle count | Default cycle count | Settings_dPCRDefaultCycleCount |
| dsDNA d | PCR default elongation per kb | Default elongation time per kb (seconds) | Settings_dPCRDefaultElongationPerKb |
| dsDNA d | PCR default polymerase | Default polymerase (from table at right) | Settings_dPCRDefaultPolymerase |
| dsDNA d | PCR default premix volume per PCR | Premix volume per PCR (everything except oligos and template) | Settings_dPCRDefaultPremixVolumePerPCR |
| dsDNA d | PCR default scale | Standard scale for PCRs AND digests, total reaction volume | Settings_dPCRDefaultScale |
| dsDNA d | PCR default variation | Default variation / additive name (from table at right) | Settings_dPCRDefaultVariation |
| dsDNA d | PCR minimum elongation time | Minimum elongation time (seconds) | Settings_dPCRMinimumElongationTime |
| dsDNA d | PCR overage floor | Overage floor - make at least this much extra premix | Settings_dPCROverageFloor |
| dsDNA d | PCR overage multiplier | Overage multiplier - call for this fraction of the amount of premix required (unitless) | Settings_dPCROverageMultiplier |
| dsDNA d | PCR premix allowed? | Allow Asst: dsDNA to call for premixing reaction components? | Settings_dPCRPremixAllowed? |
| dsDNA d | PCR premix bias | Premix bias: call for a premix even when premixing requires up to this many extra pipetting steps. | Settings_dPCRPremixBias |
| dsDNA d | PCR primer volume | Primer volume for this scale, each primer | Settings_dPCRPrimerVolume |
| dsDNA d | PCR template volume | Template volume - 0 if template volume is neglected | Settings_dPCRTemplateVolume |
| dsDNA d | Quant dID priority | Designate quantitation as a Dashboard task for dIDs when the sample "quant priority" column is set to | Settings_QCQuantdIDPriority |
| dsDNA d | Template miniprep min volume left | Minimum estimated volume remaining of mID for sheet to suggest using that mID as a template for PCR | Settings_dTemplateMiniprepMinVolumeLeft |
| dsDNA d, Minipreps m | Gel sample volume | Upper bound DNA gel sample volume used | Settings_QCGelSampleVolume |
| Golden Gate g | Default assembly scale | Default assembly scale | Settings_gDefaultAssemblyScale |
| Golden Gate g | Default buffer vol | Default buffer volume (per default assembly scale) | Settings_gDefaultBufferVol |
| Golden Gate g | Default ligase vol | Default ligase volume (per default assembly scale) | Settings_gDefaultLigaseVol |
| Golden Gate g | Default r e vol | Default restriction enzyme volume (per default assembly scale) | Settings_gDefaultREVol |
| Golden Gate g | Ligase buffer name | What to call the ligase BUFFER used in Golden Gate assemblies? (e.g. specific product name if desired) | Settings_gLigaseBufferName |
| Golden Gate g | Ligase name | What to call the ligase used in Golden Gate assemblies? (e.g. specific product name if desired) | Settings_gLigaseName |
| Golden Gate g | Min donor stock conc | Minimum stock concentration required for a miniprep to be designated for use as a Golden Gate donor (nkµ) | Settings_gMinDonorStockConc |
| Golden Gate g | Min nku GG | Warn in Status when an assembly has a final concentration lower than this for any of its parts (1 ng/kb/µL = 1.5 nM) | Settings_gMinNkuGG |
| Golden Gate g | Miniprep min conc | Minimum miniprep concentration allowed for sheet to suggest using that mID as a gID component | Settings_gMiniprepMinConc |
| Golden Gate g | Oligo donor default concentration | Concentration to use for an oligo that is registered as a Golden Gate donor (ng/µL) | Settings_gOligoDonorDefaultConcentration |
| Golden Gate g | Premix allowed? | Allow Asst: Golden Gate to call for premixing reaction components? | Settings_gPremixAllowed? |
| Golden Gate g | Premix bias | Premix bias: call for a premix even when premixing requires up to this many extra pipetting steps. | Settings_gPremixBias |
| Golden Gate g | Premix max enzyme conc factor | What fold concentration can enzymes and buffer be allowed to be in a premix, relative to their 1x final concentrations? (e.g. 2x = twice as concentrated) | Settings_gPremixMaxEnzymeConcFactor |
| Golden Gate g | Premix overage floor | Overage floor - make at least this much extra premix | Settings_gPremixOverageFloor |

153

| Tab(s) affected | Setting name | Description | Named Range designation in CC |
|---|---|---|---|
| Golden Gate g | Store in plates? | Store completed Golden Gate assemblies in storage plates (checked) or as labeled individual samples (unchecked)? | Settings_gStoreInPlates? |
| Golden Gate g | Target nku golden gate | Default target molar concentration for each part in assemblies | Settings_gTargetNkuGoldenGate |
| Golden Gate g | Wait time before transforming | Golden Gate assemblies are listed as available to transform this long after the date they are listed as assembled (0 = available immediately) | Settings_gWaitTimeBeforeTransforming |
| Golden Gate g, Analytical PCR | Colony analytical PCR threshold | gID colony analytical PCR threshold | Settings_gIDColonyAnalyticalPCRThreshold |
| Golden Gate g, Minipreps m | Gel problem bars use as donor | If a sequence-verified miniprep has agarose gel electrophoresis analysis that suggests a potential problem (e.g. multimerization), prevent that prep from being suggested as a Golden Gate part source? | Settings_gGelProblemBarsUseAsDonor |
| Golden Gate g, Minipreps m, Analytical PCR | Miniprep analytical PCR threshold | gID-derived mID analytical PCR threshold | Settings_gIDMiniprepAnalyticalPCRThreshold |
| Golden Gate g, Minipreps m, Analytical PCR | Miniprep sequencing require analytical PCR? | For mIDs derived from gIDs: If analytical PCR is called for based on the priority levels set below (mID status shows "ready to queue analytical PCR"), require at least 1 analytical PCR to be queued and completed before the mID is marked "ready to queue sequencing"? | Settings_gIDMiniprepSequencingRequireAnalyticalPCR? |
| Minipreps m | Automatic ready to transform option | What newly sequence-verified mIDs should be marked as "ready to transform for strain storage" automatically? | Settings_mAutomaticReadyToTransformOption |
| Minipreps m | Culture age | If cultures are older than this, discard and grow new cultures for minipreps | Settings_mCultureAge |
| Minipreps m | Default culture volume | Default culture volume | Settings_mDefaultCultureVolume |
| Minipreps m | Default media | Default media: | Settings_mDefaultMedia |
| Minipreps m | Default purification method | Default purification method | Settings_mDefaultPurificationMethod |
| Minipreps m | Enforce max one prep stored? | Prevent more than one sequence-verified mID from being "ready to transform for strain storage?" even if multiple mIDs are manually set as (store this prep)?<br><br>(If checked, only the first such mID will be set as ready to transform.) | Settings_mEnforceMaxOnePrepStored? |
| Minipreps m | Gel mID priority | Designate gel sample prep as a Dashboard task for mIDs when the sample "gel priority" column is set to | Settings_QCGelmIDPriority |
| Minipreps m | Low miniprep conc cutoff | Low concentration cutoff for minipreps- automatically mark mID below this value as failed and unusable | Settings_QCLowMiniprepConcCutoff |
| Minipreps m | Low miniprep conc warn | Warning concentration cutoff for minipreps - flag mIDs with concentrations below this value as potential failures | Settings_QCLowMiniprepConcWarn |
| Minipreps m | Media options | List of growth media you use for minipreps | Settings_mMediaOptions |
| Minipreps m | MID gel sample threshold | mID gel sample threshold | Settings_QCmIDGelSampleThreshold |
| Minipreps m | MID quant threshold | mID quant threshold | Settings_QCmIDQuantThreshold |
| Minipreps m | Min volume left | Minimum estimated volume remaining of mID for sheet to suggest using that mID as a gID component | Settings_mMinVolumeLeft |
| Minipreps m | Plate age | If plates are older than this, show warning in status for "ready to inoculate" | Settings_mPlateAge |
| Minipreps m | Quant mID priority | Designate quantitation as a Dashboard task for mIDs when the sample "quant priority" column is set to | Settings_QCQuantmIDPriority |
| Minipreps m | Standard elution volume | Standard elution volume | Settings_mStandardElutionVolume |
| Minipreps m | Warn volume left | Estimated volume remaining of mID for sheet to warn about using an mID as a gID component | Settings_mWarnVolumeLeft |
| Minipreps m, Assemblies a | AID analytical PCR after transformation priority | Designate analytical PCR as a Dashboard task for aID-derived transformations when the sample "analytical PCR priority" column is set to | Settings_QCaIDAnalyticalPCRAfterTransformationPriority |
| Minipreps m, Assemblies a | AID analytical PCR mID priority | Designate analytical PCR as a Dashboard task for aID-derived mIDs when the sample "analytical gel priority" column is set to | Settings_QCaIDAnalyticalPCRmIDPriority |
| Minipreps m, dsDNA d | Quantitation volume | Upper bound DNA quantitation volume used (for nanodrop, Take3, Qubit, etc.) | Settings_QCQuantitationVolume |
| Minipreps m, Golden Gate g | GID analytical PCR after transformation priority | Designate analytical PCR as a Dashboard task for gID-derived transformations when the sample "analytical PCR priority" column is set to | Settings_QCgIDAnalyticalPCRAfterTransformationPriority |
| Minipreps m, Golden Gate g | GID analytical PCR mID priority | Designate analytical PCR as a Dashboard task for gID-derived mIDs when the sample "analytical gel priority" column is set to | Settings_QCgIDAnalyticalPCRmIDPriority |
| Oligos o | Cost base100nmol plate | Cost per base 100 nmol plate | Settings_oCostBase100nmolPlate |

| Tab(s) affected | Setting name | Description | Named Range designation in CC |
|---|---|---|---|
| Oligos o | Cost base100nmol tube | Cost per base 100 nmol tube | Settings_oCostBase100nmolTube |
| Oligos o | Cost base25nmol plate | Cost per base 25 nmol plate | Settings_oCostBase25nmolPlate |
| Oligos o | Cost base25nmol tube | Cost per base 25 nmol tube | Settings_oCostBase25nmolTube |
| Oligos o | Cost u100nmol plate | Cost per ideoxyU 100 nmol plate | Settings_oCostU100nmolPlate |
| Oligos o | Cost u100nmol tube | Cost per ideoxyU 100 nmol tube | Settings_oCostU100nmolTube |
| Oligos o | Cost u25nmol plate | Cost per ideoxyU 25 nmol plate | Settings_oCostU25nmolPlate |
| Oligos o | Cost u25nmol tube | Cost per ideoxyU 25 nmol tube | Settings_oCostU25nmolTube |
| Oligos o | Oligo max length | Length of oligo above which you are not willing to order (bases) | Settings_oOligoMaxLength |
| Oligos o | Oligo min length | Minimum length of oligos you can / are willing to order (bases) | Settings_oOligoMinLength |
| Oligos o | Oligo min order | What is the least number of oligos you want to order at once? (e.g. to fulfill the minimum order size for oligos in plates)? | Settings_oOligoMinOrder |
| Oligos o | Oligo synth scale change len | Length of oligo above which price / synthesis scale changes from your preferred vendor - oligos longer than this will be flagged (bases) | Settings_oOligoSynthScaleChangeLen |
| Oligos o | Oligo validation substring length diff | When checking for oligos that are substrings of each other, how different in length can the oligos be to consider whether they might be substring matches? (setting this too high will lead to huge numbers of substring matches for long oligos) | Settings_oOligoValidationSubstringLengthDiff |
| Oligos o | Secondary structure header text | For optional secondary structure entry, what temperature (or other conditions) and/or units should appear in the column header? (Choose whatever preferred design software provides) | Settings_oSecondaryStructureHeaderText |
| Oligos o | Working concentration | Oligo working solution concentration (for reference only), μM | Settings_oWorkingConcentration |
| Registry | Antibiotics abbrs | Abbreviations you want to display for the antibiotics you use | Settings_RegistryAntibioticsAbbrs |
| Registry | Antibiotics descriptions | Names and/or descriptions of the antibiotics you use | Settings_RegistryAntibioticsDescriptions |
| Registry | Expected min size backbone | Typical or average plasmid backbone size for Golden Gate donor plasmids (everything except the part; kb) | Settings_RegistryExpectedMinSizeBackbone |
| Registry | Month membership threshold to show | Show only departed members who participated for at least this many months (0 shows all departed members) | Settings_RegistryMonthMembershipThresholdToShow |
| Registry | Months to keep departed | Months after departure date to keep departed members on the roster for | Settings_RegistryMonthsToKeepDeparted |
| Registry | Origin compatibility group table | List of replication origin categories (compatibility groups) for plasmids you work with | Settings_RegistryOriginCompatibilityGroupTable |
| Registry | Sanger read average bases | Estimated average useable bases per read for Sanger sequencing (kbases) | Settings_RegistrySangerReadAverageBases |
| Registry | Sanger read cost | Sanger sequencing cost per read ($) | Settings_RegistrySangerReadCost |
| Registry | Warn if not pABC123 | Warn if registered construct names don't conform to the pattern pABC123 (except direct receipt constructs)? | Settings_RegistryWarnIfNotpABC123 |
| Registry | Whole plasmid seq cost | Whole plasmid sequencing cost ($) | Settings_RegistryWholePlasmidSeqCost |
| Sequencing | Primer volume | Primer volume per Sanger sequencing rxn (at default working concentration) | Settings_SequencingPrimerVolume |
| Sequencing | Require analytical PCR? | If analytical PCRs are queued for an mID, require that they be completed before marking it as ready to queue for sequencing? | Settings_SequencingRequireAnalyticalPCR? |
| Sequencing | Require gel? | Require gel data coded before marking mIDs as ready to queue for sequencing? | Settings_SequencingRequireGel? |
| Sequencing | Require QC? | Require QC data before marking dIDs as ready to mix for sequencing? | Settings_SequencingRequireQC? |
| Sequencing | Require quant? | Require mIDs to be quantitated (concentration determined) before marking them as ready to queue for sequencing? | Settings_SequencingRequireQuant? |
| Sequencing | Sanger plasmid default conc | Concentration to assume for unquantitated mIDs used in Sanger sequencing (ng/μL) | Settings_SequencingSangerPlasmidDefaultConc |
| Sequencing | Sanger plasmid targetng | Plasmid ng target per Sanger sequencing reaction | Settings_SequencingSangerPlasmidTargetng |
| Sequencing | Sanger scale | Sanger sequencing rxn total volume (μL) | Settings_SequencingSangerScale |
| Sequencing | Whole plasmid conc warning | Whole plasmid sequencing concentration minimum - warn if concentration is below this value: | Settings_SequencingWholePlasmidConcWarning |
| Sequencing | Whole plasmid high conc threshold | Concentration threshold above which to use "high concentration" volume | Settings_SequencingWholePlasmidHighConcThreshold |
| Sequencing | Whole plasmid high conc volume | Volume for whole plasmid sequencing for "high concentration" DNA | Settings_SequencingWholePlasmidHighConcVolume |
| Sequencing | Whole plasmid low conc volume | Volume for whole plasmid sequencing for "low concentration" DNA | Settings_SequencingWholePlasmidLowConcVolume |

155

| Tab(s) affected | Setting name | Description | Named Range designation in CC |
|---|---|---|---|
| Strains s | Storage plate size strains | How do you want your physical STRAINS to be stored? This setting affects only the Strains s tab.<br><br>Enter your choice using the following format:<br>"#w" = # well plate<br>"b#" = box with # spaces<br>"#r#c" = plate with # rows, # cols<br><br>If you're using custom-sized boxes or plates (b# or #r#c) that don't appear in the dropdown menu, you'll need to type in your desired plate/box size using the specified format and ignore the data validation error. | Settings_SampleStoragePlateSizeStrains |
| Transformations t | Default amount | Default volume to transform (µL) | Settings_TransformationsDefaultAmount |
| Transformations t | Default comp cell vol | Default comp cell volume (to go with default comp cell procedure/batch) (µL) | Settings_TransformationsDefaultCompCellVol |
| Transformations t | Default growth temp | Default growth temperature (° C) | Settings_TransformationsDefaultGrowthTemp |
| Transformations t | Default plate amount | Default volume to plate (µL) | Settings_TransformationsDefaultPlateAmount |
| Transformations t | Default procedure | Default procedure | Settings_TransformationsDefaultProcedure |
| Transformations t | Default recovery medium | Default recovery medium | Settings_TransformationsDefaultRecoveryMedium |
| Transformations t | Default recovery time | Default recovery time (min) | Settings_TransformationsDefaultRecoveryTime |
| Transformations t | Default recovery volume | Default recovery volume (µL) | Settings_TransformationsDefaultRecoveryVolume |
| Transformations t | Procedures list | List of procedures to show in Transformations dropdown menu | Settings_TransformationsProceduresList |

**Supplementary Text 3: CloneCoordinate Status Documentation (v0.98)**

**Portions reproduced with permission** from Jeon, E. Developing the CloneCoordinate Suite for Efficient and Informed Synthetic Bacterial Plasmid Construction. Senior Honors Thesis, Williams College (2024).

**Registry**

The entering of a new serial number indicates the creation of a distinct plasmid construct (an rID), with each construct corresponding to one entire row of the "Registry" spreadsheet. Once the rID has been certified as correctly registered, CloneCoordinate will check for any manual holds or "♥" verifications that have been entered by a user under the "What to do with this construct?" column. Both of these designations will stop any work being done on these constructs - holds, for whatever reason a user may need to place one on a construct, and "♥" verifications, which indicate that the construct's sequence has been verified as correct. Placing these checks before any other steps ensures that users won't find themselves doing unnecessary work on any rIDs. However, it also means that some operations may not appear as ready to do for some of the construct's constituent samples.

Any other manual designations entered under the "What to do with this construct?" column will also appear as the construct status. And any constructs without a properly formatted name - either a user typo or a non-construct entity - will also prompt a warning in their status.

Constructs which are direct receipts are generally treated as sequence-verified. Direct receipts which have an associated strain sample, or sID, but no miniprep, or mID, simply receive the status of "Complete ♥" as they are finished and stored samples. For direct receipts with mIDs, the construct status will tell users to wait until the associated minipreps have been purified and stored.

Of the constructs which are not direct receipts, those with mIDs that have been sequence-verified empirically also receive the "Complete ♥" status. Otherwise, constructs with some associated part (an oligomer oID, double-stranded DNA dID, assembly aID, or Golden Gate gID) will have a "Registered" status. All other constructs will be deemed simply "Queued," save for those recognized as duplicates of other entered constructs based on their Golden Gate rID composition.

**Ranges which Registry Status draws upon:**
Minipreps_m_Date_purified, Minipreps_m_Verified_construct?

**Oligos o**

The entering of a new serial number indicates the creation of a distinct DNA oligomer (Oligo, or oID) sample, with each sample corresponding to one entire row of the “Oligos o” spreadsheet. Clonecoordinate won’t return a status unless both a serial number and a nucleotide sequence have been entered for the oID.

If the oID has been certified as correctly queued, CloneCoordinate will then check for any designations containing the symbol "⛔" that have been entered by a user under the “What to do with this sample” column, as well as any manual holds. Placing these checks before any other steps ensures that users won’t find themselves doing unnecessary work on any oIDs. However, it also means that no operations whatsoever will be marked as ready to do for samples manually marked ⛔ or (hold).

At this point, the status will advance through the sequential steps of creating the physical oID sample - ordering the oligomer plate, receiving the plate shipment, and making the oligomer stock from the plate (at which point the oID will receive the “done” status).

Certain oID samples themselves can be designated as priority from the Registry by checking the “Flag all related samples as high priority?” column to prioritize all samples related to a construct. This will add them to a separate tally of priority experimental procedures on the CloneCoordinate dashboard and expedite the construction progress of their corresponding constructs. oIDs can also have “missing data?” appended to their status if columns corresponding to later operations have been filled out while earlier ones have not - a sign of inconsistent operation logging.

And finally, CloneCoordinate can also append “not yet used for dsDNA?” - dsDNA meaning double-stranded DNA samples. Excluding oIDs meant to be sequencing or analysis primers, this status tag applies to dsDNA/ultramer oIDs which haven’t been entered as a PCR template or as digest DNA for dsDNA, as well as other oIDs not yet entered as PCR primers for dsDNA.

**Ranges which Oligos Status draws upon:**
dsDNA_d_Input_DNA, dsDNA_d_DNA_to_digest, dsDNA_d_Oligo_1, dsDNA_d_Oligo_2

**dsDNA d**

The entering of a new serial number indicates the creation of a distinct double-stranded DNA (dsDNA, or dID) sample, with each sample corresponding to one entire row of the “dsDNA d” spreadsheet. dID samples may be one of three types: Anneal (annealed oligos, or oIDs), PCR (amplicons using two oID primers and a template), and Digest (treatment of input DNA, which could be a miniprep or another dsDNA d, with restriction enzymes or some other procedure).

If the dID has been certified as correctly queued, CloneCoordinate will then check for manual holds and any designations containing the symbol "⛔" that have been entered by a user under the “What to do with this sample?” column. Placing these checks before any other steps ensures that users won’t find themselves doing unnecessary work on any dIDs. However, it also means that no operations whatsoever will be marked as ready to do for samples manually marked (hold) or ⛔.

The main branching point in the status’s logic is whether the dID has been mixed and incubated. For those that have not, CloneCoordinate will make sure the component oIDs, if any, are in stock and appropriate for use (the oIDs are not lost, abandoned, or used up, signaled by a "⛔" in their status). At this point, dIDs of the Anneal type will receive the status "ready to set up."

PCR and Digest type dIDs will undergo several more checks of their input DNA (template DNA for PCRs, DNA to be digested for Digests) validity before also proceeding to construction. First, the input DNA needs to be an existing sample in CloneCoordinate - the status will search through every mID, oID, dID, rID, sID, and anything listed within the Registry to find a match. If the template DNA is listed in one of these places, then the template’s sample status is checked. If the input DNA is abandoned, further work on this dID would be pointless. If it is manually held, further work should not be done at this time. In these cases, the dID is marked with the status of the input DNA. If the input DNA’s status does not indicate that it is completed, the dID status will tell users to wait until the input DNA is ready.

dIDs that have been mixed and incubated will receive a status of “done.” These can now be marked as ready for any and all applicable quality control procedures (quantitation and/or gels). Anneals, by default, are not eligible for statuses relating to quantitation/gels unless settings_qcAnnealGelQuant? is changed to TRUE. All eligible samples must also have manual priority designations (in the “Quant priority” and “Gel priority” columns) at least as high as the adjustable thresholds in Settings & admin (at default “low” and “high” priority for quantitation and gels, respectively).

Quality control statuses will be appended to the main status because they are not mandatory to complete plasmid construction in most cases. Therefore, a sample will appear as “done” regardless of the condition of quality control steps, unless a quality control outcome shows the sample is not viable, in which case the “done” status is replaced with “failed”.

CloneCoordinate will flag any dIDs which have been measured to have concentrations too low to be viable for future plasmid construction, based on the appropriate settings_qcLowPCRConcCutoff (currently set at a default 15 ng/µl) or settings_qcLowDigestConcCutoff (default 10 ng/µl). Samples that have not been quantitated or that are above the thresholds are marked as “done”, signaling that they are available and appropriate to use for other operations.

In addition to the minimum concentrations for viability, there are warning thresholds (by default, settings_dMinVolumeToStillUse = 5 µL, settings_qcLowPCRConcWarn = 25 ng/µL, settings_qcLowDigestConcWarn = 20 ng/µL). Viable samples which find themselves below these levels will receive an appended warning status, as a suggestion for users to manually designate them for additional QC checks or be cautious about using them in other operations. A gel result that assesses band position as not matching the dID's expected size will result in a "gel problem?" appendage to the status. This suggests that the PCR amplicon or digest output may not be as intended..

dIDs that go on to be part of an assembly that generates a verified miniprep will also receive the "assembly success ♥" status appendage. This is a mark of some level of trustworthiness since the dID was, at worst, not a complete failure. If given the choice, users might want to use samples of this kind in their assemblies going forward.

Certain dID samples themselves can be designated as priority from the Registry by checking the "Flag all related samples as high priority?" column to prioritize all samples related to a construct. This will add them to a separate tally of priority experimental procedures on the CloneCoordinate dashboard and expedite the construction progress of their corresponding constructs. dIDs can also be deemed to have "no active uses" in their status if they have been added to the Registry list of abandoned dIDs. And dIDs will have "missing data?" appended to their status if columns corresponding to later operations have been filled out while earlier ones have not - a sign of inconsistent operation logging.

**Ranges which dsDNA Status draws upon:**
settings_qcLowPCRConcCutoff, settings_qcLowDigestConcCutoff, settings_dMinVolumeToStillUse, settings_qcAnnealGelQuant?, settings_QCPriorityList, settings_dMinVolumeWarn, settings_dMinVolumeToStillUse, settings_qcLowPCRConcWarn, settings_qcLowDigestConcWarn

Oligos_o_Status, Oligos_o_ID, Oligos_o_Stocks_made
Minipreps_m_Status, Minipreps_m_ID
dsDNA_d_Status, dsDNA_d_ID
Registry_Status, Registry_ID, Registry_Construct_name, Registry_List_of_abandoned_dIDs
Strains_s_Status, Strains_s_ID

**Assemblies a**

The entering of a new serial number indicates the creation of a distinct assembly (aID) sample, with each sample corresponding to one entire row of the “Assemblies a” spreadsheet. Clonecoordinate won’t return a status unless both a serial number and at least one assembly part have been entered for the aID.

The status will then run a check on the entered construct for the aID. The construct must also be registered in the “Registry” spreadsheet, and its respective status can’t be abandoned or have a manual hold. The status will check the aID itself for any designations containing the symbol "⛔" that have been entered by a user under the “What to do with this sample” column. Placing these checks before any other steps ensures that users won’t find themselves doing unnecessary work on any aIDs. However, it also means that no operations whatsoever will be marked as ready to do for samples manually marked (hold) or ⛔.

If the aID has been certified as correctly queued, the status will reach the first major branching point in its logic of whether the aID has been assembled. For those that have not, CloneCoordinate will run a series of checks on the calculated assembly conditions and the component parts. The status will alert the user if the entered part volumes for the assembly combined result in a negative volume of water (less than -0.3 µL). The status will also report whether any of the individual part statuses contain "⛔" - signifying some disqualifying issue regarding its part - or indicate incorrect/incomplete queueing or an incomplete part sample. If the assembly does not have a manual hold at this point, its status will mark it as ready to be assembled and list its overqueued parts, if any.

aIDs which have been assembled will first be checked for any manual hold. For those that have not been transformed before, individual part statuses are checked again for “⛔,” as part samples could have been lost or contaminated between the aID’s assembly and its transformation. Then, the status will check for any designations containing the phrase “don’t transform” that have been entered by a user under the “What to do with this sample?” column. This might be an instance where a user wants to simply assemble a plasmid construct without transformation and subsequent testing for verification. If the total volume queued for this transformation doesn’t exceed the original volume made of this aID, the status will mark the aID as ready to transform.

By this point, the remaining aIDs have all had a transformation attempted in the past. If an aID’s most recent transformation attempt was manually terminated (via a "⛔" designation in the Transformations spreadsheet), the aID status will prompt users to look over this last attempt to determine what the best course of action should be before queueing a new transformation attempt for the aID.

For aIDs which have been manually designated for retransformation (from the “What to do with this sample? column in the Assemblies spreadsheet) and those which have new transformation attempts queued but not completed, the status will once again check that the total queued volume for the transformation (including previous attempts) doesn’t exceed the original volume made of the aID. Now these aIDs are ready to be transformed, and statuses will mark transformed plates of aIDs as ready to score as well the day after their transformation, provided that they do not have a "⛔" designation. If a transformation yields no colonies at all, its corresponding aID status will detect that progress on this transformation has been stalled, and further steps such as plate scoring will not appear.

Once a construct has a “complete” status in the registry, its aID will read “done; construct complete” as plate scoring and confirmation of a correct phenotype is the final step of actually assembling a construct - procedures such as sequencing, analytical PCR, and strain storage are technically optional and carried out depending on the constructs’ final purpose.

Some aIDs might be preset to require an analytical PCR before a corresponding mID is queued (Settings_analytical_PCR_ aIDRequireQueueingAfterTransformation, not required by default), and the aID status will make this known to the user. For aIDs which have fulfilled or never had this requirement, the status will indicate that they are ready to have a miniprep queued. Again, the status will check whether any colonies grew from transformation, as well as if the transformation volume exceeded the original aID volume - unsuccessful transformations (those containing “don’t” or “can’t” in their transformation status, or simply grew incorrect colonies) of sufficient remaining aID volume will automatically be labeled for retransformation, while the other unsuccessful transformations will remain stalled and require manual intervention by users for the next steps.

The status will notify the user of any queued minipreps waiting to be inoculated, and if nothing else goes wrong with these minipreps, their aIDs will eventually attain a “done” status.

Certain aID samples themselves can be designated as priority from the Registry by checking the “Flag all related samples as high priority?” column to prioritize all samples related to a construct. This will add them to a separate tally of priority experimental procedures on the CloneCoordinate dashboard and expedite the construction progress of their corresponding constructs. aIDs can also be deemed to have a "♥ good mID" in their status if they have had a corresponding miniprep be empirically sequence-verified. The status will also append whether the aID volume has run out. Even if an aID has run out in this way, users will still be able to view whether it has resulted in a sequence-verified miniprep, as neither of these appended labels are mutually-exclusive. And aIDs will have “missing data?” appended to their status if columns corresponding to later operations have been filled out while earlier ones have not - a sign of inconsistent operation logging.

**Ranges which Assemblies Status draws upon:**

Settings_analytical_PCR_aIDRequireQueueingAfterTransformation

Registry_Status, Registry_Construct_name
Transformations_Transformed_amount, Transformations_Source_ID,
Transformations_What_to_do_with_this, Transformations_Date_transformed
Minipreps_m_Status, Minipreps_m_Source_ID, Minipreps_m_Date_purified
Lookup_lists_and_tabulations_Verified_mID_Sources

**Golden Gate g**

The entering of a new serial number indicates the creation of a distinct golden gate (gID) sample, with each sample corresponding to one entire row of the "Golden Gate g" spreadsheet. Clonecoordinate won't return a status unless a serial number has been entered for the gID.

If the gID has been certified as correctly queued and does not have any designation containing the symbol "⛔" entered by a user under the "What to do with this sample" column, the status will run a check on the entered construct for the gID - if the construct has been abandoned or completed based on its status in the Registry, as well as if it has a manual hold, the gID status will reflect this. Placing these checks before any other steps ensures that no operations whatsoever will be marked as ready to do for these samples so that users won't find themselves doing unnecessary work on any gIDs.

Now, the status will reach the first major branching point in its logic of whether the gID has been assembled. For those that have not, CloneCoordinate will run a series of checks on the calculated assembly conditions and the component parts. The status will alert the user if the entered part volumes for the assembly combined result in a negative volume of water (less than -0.3 µL). The status will also report on the state of the constructs from which the individual golden gate parts are derived. CloneCoordinate will detect whether any parts fail to be sourced to constructs, whether they have yet to be cloned, whether they need to be stocked and have their concentrations determined, and whether any have received a manual hold.

At this point, if the golden gate itself does not have a manual hold, the status will prompt the user to enter the part mIDs and volumes for the assembly. Any invalid mID inputs (where the mID does not correspond with the rID) will be pointed out, as well as whether the part junctions remain valid with these mIDs. Otherwise, the status will indicate that the golden gate is ready for assembly. If any part concentrations are lower than the preset threshold (settings_gMinNkuGG, by default 0.3 ng/kb/µL), CloneCoordinate will append a warning to the "ready to assemble" status. A list of overqueued parts, if any, will also be appended.

If there is no manual hold, the status will now advance to deal with gIDs which have been assembled. Users can preset the amount of time which must pass before assembled gIDs can be queued for transformation (settings_gWaitTimeBeforeTransforming, by default one day) as well as enter a "don't transform" designation under the "What to do with this sample" column, both of which will affect whether the gID will appear as "ready to transform." The status will check for and rule out transformations of gIDs for which more total volume has been queued for transformation than was made of the gID itself. And if a gID's most recent transformation attempt was manually terminated (via a "⛔" designation in the Transformations spreadsheet), the gID status will prompt users to look over this last attempt to determine what the best course of action should be before queueing a new transformation attempt for the gID. Otherwise, gIDs will be marked as ready to transform.

For gIDs which have been manually designated for retransformation (from the "What to do with this sample? column in the Golden Gate spreadsheet) and those which have new transformation attempts queued but not completed, the status will once again check that the total queued volume for the transformation (including previous attempts) doesn't exceed the original volume made of the gID. Now these gIDs are ready to be transformed, and statuses will mark transformed plates of gIDs as ready to score as well the day after their transformation, provided that they do not have a "⛔" designation. If a transformation yields no colonies at all, its corresponding gID status will detect that progress on this transformation has been stalled, and further steps such as plate scoring will not appear.

Once a construct has a “complete” status in the registry, its gID will read “done; construct complete” as plate scoring and confirmation of a correct phenotype is the final step of actually assembling a construct - procedures such as sequencing, analytical PCR, and strain storage are technically optional and carried out depending on the constructs’ final purpose.

The status will check for unsuccessful transformations (those containing “don’t” or “can’t” in their transformation status, or simply grew incorrect colonies or none at all) will be labeled as stalled and require manual intervention by users for the next steps. Some gIDs might be preset to require an analytical PCR before a corresponding mID is queued (Settings_analytical_PCR_ gIDRequireQueueingAfterTransformation, not required by default), and the gID status will make this known to the user. For gIDs which have fulfilled or never had this requirement, the status will indicate that they are ready to have a miniprep queued or, if queued, waiting to be inoculated. If nothing else goes wrong with these minipreps, their gIDs will eventually attain a “done” status.

Certain gID samples themselves can be designated as priority from the Registry by checking the “Flag all related samples as high priority?” column to prioritize all samples related to a construct. This will add them to a separate tally of priority experimental procedures on the CloneCoordinate dashboard and expedite the construction progress of their corresponding constructs. gIDs can also be deemed to have a "♥ good mID" in their status if they have had a corresponding miniprep be empirically sequence-verified. The status will also append whether the gID volume has run out. Even if an gID has run out in this way, users will still be able to view whether it has resulted in a sequence-verified miniprep, as neither of these appended labels are mutually-exclusive. And gIDs will have “missing data?” appended to their status if columns corresponding to later operations have been filled out while earlier ones have not - a sign of inconsistent operation logging.

**Ranges which Golden Gate Status draws upon:**

settings_gMinNkuGG, settings_gWaitTimeBeforeTransforming,
Settings_analytical_PCR_gIDRequireQueueingAfterTransformation

Registry_Status, Registry_Construct_name, Registry_GG_parts_display_name, Registry_5_overhang,
Registry_ID, Registry_3_overhang
Minipreps_m_Construct_name, Minipreps_m_ID
Transformations_Transformed_amount, Transformations_Source_ID,
Transformations_What_to_do_with_this, Transformations_Date_transformed
Minipreps_m_Status, Minipreps_m_Source_ID, Minipreps_m_Date_purified
Lookup_lists_and_tabulations_Verified_mID_Sources

**Transformations**

The entering of a new serial number indicates the creation of a distinct transformation attempt, with each transformation corresponding to one entire row of the “Transformations” spreadsheet. If the transformation has been certified as correctly queued, CloneCoordinate will then check that the corresponding construct of the source ID exists in the Registry spreadsheet, as well as whether this construct has been abandoned or placed on a manual hold by a user under the “What to do with this sample?” column in the Registry. Placing these checks before any other steps ensures that users won’t find themselves doing unnecessary work on any transformations for constructs of jeopardized status.

Transformations themselves, as well as their source IDs (aIDs, gIDs, and mIDs) can be given designations containing the symbol "⛔" by users under the “What to do with this sample?” column in their respective spreadsheets. All of these will similarly halt or terminate progress on their transformations. If no such designations are present, then we are ready to transform. The status will make note of whether this is a first-time transformation or a repeat, and whether previous transformation attempts have been carried out or simply queued.

The status will warn the user if the combined volume of queued transformation attempts for a source ID fall short of the preset default transformation volume (settings_transformationsDefaultAmount, set to 5 μL by default) - transformation attempts using extremely small volumes can result in suboptimal colony growth. The status will also warn again if any parts have received a "⛔" designation for transformations of aIDs. Both of these warnings are appended to the appropriate “ready to transform” status, as the transformation may still be carried out, just with suboptimal results.

Plates are to be scored a day after the transformations are done, and any plates with no colonies grown will be treated as a finished transformation attempt (a status of “done,” appended with “no colonies!”). Plates with colony growth but no correct phenotypes will also finish with an appendage noting their ineligibility for accurate miniprep queueing.

This status can also call for analytical PCR to be run following transformation. Source aIDs and gIDs each have their own preset priority threshold level for warranting an analytical PCR - settings_aIDColonyAnalyticalPCRThreshold and settings_gIDColonyAnalyticalPCRThreshold, respectively, each comprised of four levels (“do not do,” “low,” “normal,” and “high”) and set to “high” by default (mIDs, by default, are set between “do not do” and “low,” allowing analytical PCRs in general to be run but not qualifying them for any thresholds requiring at least a “low” priority). If a user manually enters an analytical PCR priority for a transformation which matches or exceeds these preset thresholds, that transformation will receive a status of "ready to queue analytical PCR.” The status will reflect ongoing progress on analytical PCRs until their completion, upon which their transformations will receive a “done” status.

Certain transformations themselves can be designated as priority from the Registry by checking the “Flag all related samples as high priority?” column to prioritize all samples related to a construct. This will add them to a separate tally of priority experimental procedures on the CloneCoordinate dashboard and expedite the construction progress of their corresponding constructs. Transformations will also have “missing data?” appended to their status if columns corresponding to later operations have been filled out while earlier ones have not - a sign of inconsistent operation logging.

**Ranges which Transformations Status draws upon:**

settings_transformationsDefaultAmount, settings_aIDColonyAnalyticalPCRThreshold, settings_gIDColonyAnalyticalPCRThreshold, settings_qcPriorityList

Registry_Status, Registry_Construct_name
Assemblies_a_What_to_do_with_this, Assemblies_a_ID
Golden_Gate_g_What_to_do_with_this, Golden_Gate_g_ID
Minipreps_m_What_to_do_with_this, Minipreps_m_ID
Transformations_Source_ID, Transformations_Date_transformed, Transformations_Transformed_amount

**Minipreps m**

The entering of a new serial number indicates the creation of a distinct miniprep (mID) sample, with each sample corresponding to one entire row of the “Minipreps m” spreadsheet. Clonecoordinate will return a blank status until a serial number has been entered for the mID, then will prompt the user until the mID has been certified as correctly queued.

Minipreps are a very central sample type within CloneCoordinate - they act as a link between aIDs and gIDs, their transformations (tIDs), and many quality control procedures. The mID status will first check for any discrepancies between the manual status designations entered for the mID (under the “What to do with this sample?” columns in both the Minipreps and Sequencing spreadsheets), then for any manual holds entered in either spreadsheet (specifying from which it comes from). Conflicting manual designations may be an indication of incorrect information at some point in CloneCoordinate, or they may be misinputs themselves. If quantitated, a miniprep with a concentration below the preset cutoff minimum (15 ng/μL, by default) will be given a failed status. Minipreps with corresponding constructs that have been abandoned or manually held will similarly be held, and those with no growth following inoculation will also be labeled as stalled. Placing these checks before any other steps ensures that users won’t find themselves doing unnecessary or incorrect procedures on any mIDs.

The status will label mIDs as stalled if the culture antibiotic does not match that of the construct marker - this will mean the growth from the culture may contain other bacteria than that of the desired plasmid. Then following a successful inoculation, the status will prompt users to prepare a miniprep, given that a day has passed since inoculation but not enough days to exceed the preset maximum time (Settings_mCultureAge, by default two days) to disqualify a culture as too old to miniprep.

Any manual status designation on a transformation (under the “What to do with this sample?” column in Transformations) will disqualify its associated mID from further steps until removed. CloneCoordinate also has a “Plate status” column which performs its own checks on the transformation plates for each mID - for mIDs derived from aIDs and gIDs, the mID status will call for a retransformation if no available transformation plate is detected, and it will also prompt users to manually examine the transformation plate if its status has deemed it to be of questionable age. mIDs with positive plate statuses will be labeled as ready for inoculation. For mIDs derived from sIDs at this stage, the mID status will wait until the corresponding strains have been stored before labeling the mID as ready for inoculation.

For any remaining mIDs which have not been purified, those with no source ID entered will return a blank status, and all others will appear as “stalled.”

As long as their volumes have not run below the preset minimum (Settings_mMinVolumeLeft, 5.00 μL by default), purified minipreps will receive a “done” status. Any manual hold (from either Minipreps or Sequencing) will be appended, specifying from which spreadsheet it comes from. The status will also append whether the miniprep growth was an unexpected phenotype/color.

There are many paths that a miniprep could take from this point, not necessarily in any order, so these minipreps will receive many status appendages for the different procedures they can receive. mIDs with manual priority designations for DNA quantitation and gel electrophoresis (in the “Quant priority” and “Gel priority” columns) at least as high as the adjustable thresholds 'Settings & admin'!$BH$29 and 'Settings & admin'!$BH$27 (both set to “normal” priority by default) will appear ready for these steps in their statuses. Minipreps of aIDs and gIDs for which users have entered analytical PCR priority at least as high as their preset thresholds (settings_aIDMiniprepAnalyticalPCRThreshold and settings_gIDMiniprepAnalyticalPCRThreshold, both set to “high” priority by default) will also appear ready for analytical PCR. The mID status will also advance through the stages of a gel - prepping the sample, running the gel, then coding the gel - as they are completed.

The mID status will also append whether a miniprep should be sequenced. This depends on a number of checks. The construct corresponding to the mID must have been checked off for sequencing by a user (under the “Currently needs sequencing?” column in the Registry). If quality control procedures (quantitation, gel, analytical PCR) have been preset as required before sequencing (via settings_SequencingRequireQuant?, settings_SequencingRequireGel?, and settings_SequencingRequireAnalyticalPCR?, all required by default), they must also be completed before “ready to queue sequencing” is appended to the mID status. The analytical PCR requirement for sequencing can also further be specified between mIDS derived from aIDs and gIDs (via settings_aIDMiniprepSequencingRequireAnalyticalPCR? and settings_gIDMiniprepSequencingRequireAnalyticalPCR?, both disabled by default as they are overridden by settings_SequencingRequireAnalyticalPCR?). There is also a final “Is sequencing appropriate?” column in Minipreps which rules out additional scenarios which would disqualify mIDs from sequencing (those with rID and sID sources, those from colonies of unexpected phenotype/color, etc.) - any result in this column containing “Do not” will disqualify mIDs from sequencing, while a result of “Check QC” will direct users to manually review the ambiguous quality control data and decide whether to manually queue the mIDs for sequencing.

The mID status will append warnings for mIDs which fall below the preset warning threshold, but above the preset working minimum, for volume (Settings_mWarnVolumeLeft and Settings_mMinVolumeLeft), as well as those below the preset warning threshold for concentration (settings_qcLowMiniprepConcWarn). CloneCoordinate also has a system for rating the quality and intensity of gel results as good, problematic, or ambiguous (via the pairs of settings_GelCodingQualityFlags with settings_GelCodingQuality and settings_GelCodingIntensityFlags with settings_GelCodingIntensity) - if any problematic results are detected, a warning will be appended to the mID status of this possibility.

The setting settings_mAutomaticReadyToTransformOption allows users to preset which newly sequence-verified mIDs should be marked as "ready to transform for strain storage" automatically - all of them, only new ones manually designated for storage, or none. While currently set to automatically mark all by default, the status will check for manual designations should the setting be adjusted. And those which have been transformed in this way will then be marked as ready to be inoculated for storage.

Certain mID samples themselves can be designated as priority from the Registry by checking the “Flag all related samples as high priority?” column to prioritize all samples related to a construct. This will add them to a separate tally of priority experimental procedures on the CloneCoordinate dashboard and expedite the construction progress of their corresponding constructs.

mIDs can also receive the "♥ verified" status appendage if they have been sequence-verified. This applies to mIDs derived from sIDs which have been designated to reprep their respective plasmids and do not have any "⛔" manual designation (under the “What to do with this sample?” columns in the

Minipreps, Sequencing, or Strains spreadsheets). All mIDs of rIDs will automatically appear as verified. mIDs of any remaining source ID types (such as aIDs and gIDs) will appear as verified if they have the verification indicator "♥" in a manual designation (under the "What to do with this sample?" columns in the Minipreps or Sequencing, specified if from Sequencing), with an appended note if the compiled sequencing data does not definitely indicate sequence verification (in the "Compiled sequencing results" column).

And mIDs will have "missing data?" appended to their status if columns corresponding to later operations have been filled out while earlier ones have not - a sign of inconsistent operation logging.

**Ranges which Minipreps Status draws upon:**

settings_qcLowMiniprepConcCutoff, Settings_mMinVolumeLeft,
settings_aIDMiniprepAnalyticalPCRThreshold, settings_gIDMiniprepAnalyticalPCRThreshold
settings_qcPriorityList, settings_SequencingRequireQuant?, settings_SequencingRequireGel?
settings_SequencingRequireAnalyticalPCR?, settings_gIDMiniprepSequencingRequireAnalyticalPCR?,
settings_aIDMiniprepSequencingRequireAnalyticalPCR?, Settings_mWarnVolumeLeft,
settings_qcLowMiniprepConcWarn, settings_GelCodingQualityFlags, settings_GelCodingQuality,
settings_GelCodingIntensityFlags, settings_GelCodingIntensity,
settings_mAutomaticReadyToTransformOption, Settings_mCultureAge

Registry_Manual_status_assignment, Registry_Construct_name, Registry_Needs_sequencing?
Strains_s_Status, Strains_s_Storage_strain?, Strains_s_Date_stored
Transformations_What_to_do_with_this, Transformations_Source_ID

**Analytical PCR**

The entering of a new serial number indicates the creation of a distinct analytical PCR sample, with each sample corresponding to one entire row of the “Analytical PCR” spreadsheet. If the analytical PCR has been certified as correctly queued, CloneCoordinate will then check for manual holds and any designations containing the phrase “don’t use” that have been entered by a user under the “What to do with this sample?” column. Placing these checks before any other steps ensures that users won’t find themselves doing unnecessary work on any analytical PCRs. However, it also means that no operations whatsoever will be marked as ready to do for samples manually marked (hold) or ⛔.

CloneCoordinate will also check the status of the construct corresponding to the analytical PCR. Abandoned constructs can be preset to never have analytical PCRs performed on their samples (via Settings_analytical_PCR_NoAbandonedConstructs, true by default), and constructs with manual holds will result in holds for their analytical PCRs as well.

Before an analytical PCR is run, its status will check through the statuses of its PCR components. Both the forward and reverse oligomer oIDs must be finished and without any designations containing the phrase “don’t use” entered by a user under the “What to do with this sample?” column on the “Oligos o” spreadsheet. The template DNA sample - aID, gID, mID, or sID - must also be finished. The analytical PCR can be run at this point, and afterward the status will proceed through the sequential steps of running, and then coding, a corresponding gel sample.

Analytical PCRs will have “missing data?” appended to their status if columns corresponding to later operations have been filled out while earlier ones have not - a sign of inconsistent operation logging.

**Ranges which Analytical PCR Status draws upon:**
Settings_analytical_PCR_NoAbandonedConstructs
Registry_Status, Registry_Construct_name
Oligos_o_Status, Oligos_o_ID

**Sequencing**

The entering of a serial number indicates the creation of a distinct sequencing sample, with each corresponding to one entire row of the "Sequencing" spreadsheet. Source IDs can be either dIDs or mIDs, and sequencing can be either Sanger or whole-plasmid.

If the sequencing has been certified as correctly queued, CloneCoordinate will then check for any designations containing the symbol "⛔" that have been entered by a user under the "What to do with this sample?" column, as well as manual holds. Placing these checks before any other steps ensures that users won't find themselves doing unnecessary work on any sequencing. However, it also means that no operations whatsoever will be marked as ready to do for samples manually marked ⛔ or (hold).

The main branching point in the status's logic is whether the sequencing sample has been mixed. For those that have not, CloneCoordinate will make sure its constituent source ID and sequencing primer oID samples don't have some disqualifying status containing "⛔," and that these samples have been finished. By default, CloneCoordinate also requires quality control data on dIDs before marking them as ready to mix for sequencing - this setting, settings_SequencingRequireQC?, can be toggled off as needed. oIDs satisfying all of these checks are ready to mix.

CloneCoordinate can now append a series of warnings to the status as applicable, starting with a possible redundancy warning for sequencing samples for completed, abandoned, or manually held template constructs. Whole-plasmid sequencing samples will also receive a warning if their template concentration falls below the preset warning threshold settings_SequencingWholePlasmidConcWarning (30ng/μl by default).

The appended warning checks for source ID volume are more extensive. The status will warn users if the current remaining volume of any source dID falls below the preset warning threshold settings_dMinVolumeWarn (10μl by default). And based on how much of the source ID has been queued in various samples across all of CloneCoordinate, the status will tell users whether the source ID volume is going to fall below the preset working minimums - settings_dMinVolumeToStillUse and Settings_mMinVolumeLeft for dIDs and mIDs, respectively, both 5μl by default - or below the warning threshold from above.

Of the sequencing samples that have been mixed, those that have been manually omitted from sequencing under the "What to do with this sample?" column will be stamped with a status of "done", so that no further steps will unnecessarily appear in their status. The remaining sequencing samples are then sequentially guided through the steps of assigning a well to the mixed sample, submitting the sample and receiving the sequencing read, and aligning and interpreting the read. Some alignments may be classified as not usable or not required to be interpreted, and these will be given a "done" status as well. The remaining alignments are then classified into "(easy)" and "(hard)" interpretations based on whether they completely match their expected nucleotide sequence.

Certain sequencing samples themselves can be designated as priority from the Registry by checking the "Flag all related samples as high priority?" column to prioritize all samples related to a construct. This will add them to a separate tally of priority experimental procedures on the CloneCoordinate dashboard and expedite the construction progress of their corresponding constructs. Sequencing samples can also have “missing data?” appended to their status if columns corresponding to later operations have been filled out while earlier ones have not - a sign of inconsistent operation logging.

**Ranges which Sequencing Status draws upon:**
settings_SequencingRequireQC?, settings_SequencingWholePlasmidConcWarning, settings_dMinVolumeWarn, settings_dMinVolumeToStillUse, Settings_mMinVolumeLeft

Registry_Status, Registry_Construct_name
Oligos_o_Status, Oligos_o_ID
dsDNA_d_Status, dsDNA_d_Volume_left
Minipreps_m_Status, Minipreps_m_Volume_left

**Strains s**

The entering of a new serial number indicates the creation of a distinct strain storage sample, with each sample corresponding to one entire row of the “Strains s” spreadsheet.  Unlike most other statuses, the strains status will first check whether this strain - corresponding to a position on a well or other physical storage setup - is unused and empty.  Prominently noting empty wells is important because strains are in cryostorage.  In terms of the spreadsheet, this would mean that the user has entered nothing under the columns for transformed DNA, plasmid name, organism, strain, and date stored.

If the strain has been certified as correctly queued, CloneCoordinate will then check for manual holds and any designations containing the symbol "⛔" that have been entered by a user under the “What to do with this sample?” column.  Placing these checks before any other steps ensures that users won’t find themselves doing unnecessary work on any strains. However, it also means that no operations whatsoever will be marked as ready to do for samples manually marked (hold) or ⛔.

The status will update as the DNA to be transformed is inoculated, then as this miniprep is successfully stored as a strain.

The user can select “Empirically verified correct ♥” under the “What to do with this sample?” column to append “♥ verified” to the status, an indication that the sequence of this stored strain has been verified through sequencing.

Certain strain storage samples can be designated as priority from the Registry by checking the “Flag all related samples as high priority?” column to prioritize all samples related to a construct. This will add them to a separate tally of priority experimental procedures on the CloneCoordinate dashboard and expedite the construction progress of their corresponding constructs.  And strains will have “missing data?” appended to their status if columns corresponding to later operations have been filled out while earlier ones have not - a sign of inconsistent operation logging.

**Ranges which Strains Status draws upon:**
Minipreps_m_Date_inoculated_for_storage

### Supplementary Text 4: Premix Documentation for Assistant Tabs

Several *in vitro* assembly tasks, as well as media preparation for culture inoculation, can benefit from premixing some of the components shared in common between multiple samples. CloneCoordinate has code to generate labor-saving (though not necessarily optimal) premixes for Asst: dsDNA, Asst: Assemblies, Asst: Golden Gate, and Asst: Minipreps. The Golden Gate premix code provides a good overview of the approach used. Premixes for other Assistant tabs are calculated along similar lines.

Golden Gate assemblies combine multiple plasmid and/or PCR product parts, known as Golden Gate donors, and assemble them scarlessly into a desired construct using a type IIS restriction enzyme and T4 DNA ligase. CloneCoordinate's Golden Gate assembly assistant sheet offers users the option to enhance efficiency and reduce pipetting errors by employing a premix during the assembly process. This premix contains standardized reaction components common to all assemblies being conducted. Importantly, the current premixing algorithm will only call for premixing components that have identical volumes in every currently selected assembly. To use a premix, the user must choose, by manually assigning sort numbers, only assemblies that share common components and ensure that they need identical volumes. Additional assemblies with a separate set of common components can then be carried out separately afterward. Implementation of a more robust and automatic premix selection algorithm is a future development goal.

Within the Golden Gate section of the settings/admin sheet, users can specify whether to activate the assistant's recommendation for using a premix (default = premixing enabled). If users opt against using a premix, the assistant will not enable premixing formulas. Otherwise, the assistant suggests premixing when the number of pipetting operations required without premixing exceeds those with a premix. The decision determines the practicality of premix utilization by comparing the sum of premixed components and assembly counts against the product of these two values. The former sum corresponds to the number of pipetting operations to (1) add each component to the premix and then (2) pipette premix into each reaction; the latter is the number of operations to put each component into each reaction separately. Users also have the option to input a custom premix bias in settings (Settings_gPremixBias, default value = 1, premixing recommended even if it requires 1 additional operation), recommending premixing even when it marginally increases pipetting operations. This flexibility acknowledges that the slight increase in pipetting may be outweighed by the improved accuracy and/or consistency that premixing allows. If premixing is calculated to be efficient based on these criteria, the assistant sheet systematically identifies components shared across assemblies and designates these as premix components. Enzymes and buffers are included in the premix provided their concentrations do not exceed a maximum fold concentration relative to their 1x final concentrations, a parameter inputted by the user in settings (Settings_gPremixMaxEnzymeConcFactor, default value = 2.0x). Moreover, the premix calculation accounts for the total number of assemblies utilizing the premix and determines the volume required, with an adjustable added overage volume configurable in the settings (Settings_gPremixOverageFloor default value = 5.00 µL) to allow for residual liquid adhering to tubes and tips during work. The total volume of premix dispensed into each assembly tube is the sum of individual component volumes within the premix. By streamlining the assembly process and minimizing human error through strategic premixing, CloneCoordinate's Golden Gate assembly assistant sheet enhances efficiency while accommodating user-specific preferences and settings.

The technical implementation of the premix calculations is as follows:

For each row, components ("Well, Part 1," "Well, Part 2," etc…) and their respective volumes ("µL 1," "µL 2," etc…) are concatenated in a list. Within this list, components and their accompanying volumes are delineated by a comma (,) while different components are separated by vertical bars (|), all stored in the "Component-volume list for this row" cell. Additionally, the type IIS restriction enzyme ("µL RE"), T4 DNA ligase ("µL T4 ligase"), and T4 DNA ligase buffer ("µL T4 ligase buffer") for each assembly are appended to the end of this cell in the same format.

In a new column, "Component sources to potentially premix, " the assistant sheet parses through each "Component-volume list for this row" cell, separating the list at each vertical bar (|). It then compares the formatted component and volume pairs across all the rows. If multiple assemblies use identical components with matching volumes, these shared components are listed in the "Component sources to potentially premix" column, with their respective volumes in the "Volumes to potentially premix" column.

To maintain enzyme and buffer concentrations within acceptable limits relative to their 1x final concentrations when added to the premix solution, a validation check is used. In the "Component sources to premix" column, components identified in the "Component sources to potentially premix" column are included unless they are enzymes or buffers. Enzymes and buffers are conditionally added based on user-defined maximum concentration parameters configured in settings (settings_gPremixMaxEnzymeConcFactor).

To determine the amount of each component to add into the premix, the volume of each component required for an individual assembly (as listed in the "Volumes to premix" column) is multiplied by the number of assemblies requiring that particular component. This product is further multiplied by an overage factor, determined by a custom setting ("settings_gPremixOverageFloor") within the Golden Gate section of the settings/admin sheet. Finally, the total premix volume ("µL Premix") for each assembly is calculated as the sum of volumes listed in the "Volumes to premix" column.

The Asst: dsDNA and Asst: Assemblies premix code operates largely analogously to the Golden Gate premix code, with separate configuration Settings. These premixes are simpler and so the decision about whether to create a premix is carried out separately for each sample, and the premix is called for if it reduces pipetting operations (accounting for the premix bias Setting). These Assistant tabs can recommend multiple premixes for a single set of samples. The Asst: Minipreps tab simply applies an overage factor to each antibiotic media type needed, and recommends a premix of media and antibiotic even for small numbers of wells/tubes needing the media type, since these are simple to account for manually.
